# Multi-scale Inference of Genetic Trait Architecture using Biologically Annotated Neural Networks

**DOI:** 10.1101/2020.07.02.184465

**Authors:** Pinar Demetci, Wei Cheng, Gregory Darnell, Xiang Zhou, Sohini Ramachandran, Lorin Crawford

## Abstract

In this article, we present Biologically Annotated Neural Networks (BANNs), a nonlinear probabilistic framework for association mapping in genome-wide association (GWA) studies. BANNs are feedforward models with partially connected architectures that are based on biological annotations. This setup yields a fully interpretable neural network where the input layer encodes SNP-level effects, and the hidden layer models the aggregated effects among SNP-sets. We treat the weights and connections of the network as random variables with prior distributions that reflect how genetic effects manifest at different genomic scales. The BANNs software uses variational inference to provide posterior summaries which allow researchers to simultaneously perform (*i*) mapping with SNPs and (*ii*) enrichment analyses with SNP-sets on complex traits. Through simulations, we show that our method improves upon state-of-the-art association mapping and enrichment approaches across a wide range of genetic architectures. We then further illustrate the benefits of BANNs by analyzing real GWA data assayed in approximately 2,000 heterogenous stock of mice from the Wellcome Trust Centre for Human Genetics and approximately 7,000 individuals from the Framingham Heart Study. Lastly, using a random subset of individuals of European ancestry from the UK Biobank, we show that BANNs is able to replicate known associations in high and low-density lipoprotein cholesterol content.

**Author Summary:** A common goal in genome-wide association (GWA) studies is to characterize the relationship between genotypic and phenotypic variation. Linear models are widely used tools in GWA analyses, in part, because they provide significance measures which detail how individual single nucleotide polymorphisms (SNPs) are statistically associated with a trait or disease of interest. However, traditional linear regression largely ignores non-additive genetic variation, and the univariate SNP-level mapping approach has been shown to be underpowered and challenging to interpret for certain trait architectures. While nonlinear methods such as neural networks are well known to account for complex data structures, these same algorithms have also been criticized as “black box” since they do not naturally carry out statistical hypothesis testing like classic linear models. This limitation has prevented nonlinear regression approaches from being used for association mapping tasks in GWA applications. Here, we present Biologically Annotated Neural Networks (BANNs): a flexible class of feedforward models with partially connected architectures that are based on biological annotations. The BANN framework uses approximate Bayesian inference to provide interpretable probabilistic summaries which can be used for simultaneous (*i*) mapping with SNPs and (*ii*) enrichment analyses with SNP-sets (e.g., genes or signaling pathways). We illustrate the benefits of our method over state-of-the-art approaches using extensive simulations. We also demonstrate the ability of BANNs to recover novel and previously discovered genomic associations using quantitative traits from the Wellcome Trust Centre for Human Genetics, the Framingham Heart Study, and the UK Biobank.

## Introduction

Over the two last decades, a considerable amount of methodological research in statistical genetics has focused on developing and improving the utility of linear models [1–13]. The flexibility and interpretability of linear models make them a widely used tool in genome-wide association (GWA) studies, where the goal is to test for statistical associations between individual single nucleotide polymorphisms (SNPs) and a phenotype of interest. In these cases, traditional variable selection approaches provide a set of *P*-values or posterior inclusion probabilities (PIPs) which lend statistical evidence on how important each variant is for explaining the overall genetic architecture of a trait. However, this univariate SNP-level mapping approach can be underpowered for “polygenic” traits which are generated by many mutations of small effect [14–19]. To mitigate this issue, more recent work has extended variable selection techniques to identify enriched gene or pathway-level associations, where groups of SNPs within a particular genomic region are combined (commonly known as a SNP-set) to detect biologically relevant disease mechanisms underlying the trait [20–27]. Still, the performance of standard SNP-set methods can be hampered by strict additive modeling assumptions; and the most powerful of these statistical approaches rely on algorithms that are computationally inefficient and unreliable for large-scale sets of data [28].

The explosion of large-scale genomic datasets has provided the unique opportunity to move beyond the traditional linear regression framework and integrate nonlinear modeling techniques as standard statistical tools within GWA analyses. Indeed, nonlinear methods such as neural networks are well known to be most powered in settings when large training data is available [29]. This includes GWA applications where consortiums have data sets that include hundreds of thousands of individuals genotyped at millions of markers and phenotyped for thousands of traits [30,31]. It is also well known that these nonlinear statistical approaches often exhibit greater predictive accuracy than linear models, particularly for complex traits with broad-sense heritability that is driven by non-additive genetic variation (e.g., gene-by-gene interactions) [32, 33]. One of the key characteristics that leads to better predictive performance from nonlinear approaches is the automatic inclusion of higher order interactions between variables being put into the model [34, 35]. For example, neural networks leverage activation functions between layers that implicitly enumerate all possible (polynomial) interaction effects [36]. While this is a partial mathematical explanation for model improvement, in many biological applications, we often wish to know precisely which subsets of variants are most important in defining the architecture of a trait. Unfortunately, the classic statistical idea of variable selection and hypothesis testing is lost within nonlinear methods since they do not naturally produce interpretable significance measures (e.g., *P*-values or PIPs) like traditional linear regression [35, 37].

In this work, we develop biologically annotated neural networks (BANNs), a nonlinear probabilistic framework for mapping and variable selection in high-dimensional genomic association studies (Fig. 1). BANNs are a class of feedforward Bayesian models with partially connected architectures that are guided by predefined SNP-set annotations (Fig. 1a). The interpretability of our approach stems from a combination of three key properties. First, the partially connected network architecture yields a hierarchical model where the input layer encodes SNP-level effects, and the single hidden layer models the effects among SNP-sets (Fig. 1b). Second, inspired by previous work in the Bayesian neural network literature [38–42], we treat the weights and connections of the network as random variables with sparse prior distributions, which flexibly allows us to model a wide range of sparse and polygenic genetic architectures (Fig. 1c). Third, we perform an integrative model fitting procedure where the enrichment of SNP-sets in the hidden layer are directly influenced by the distribution of associated SNPs with nonzero effects on the input layer. These three components collectively make for an effective nonlinear variable selection strategy for conducting association mapping and enrichment analyses simultaneously on complex traits. With detailed simulations, we assess the power of BANNs to identify significant SNPs and SNP-sets under a variety of genetic architectures, and compare its performance against multiple competing approaches [21,23,25–27,43–46]. We also apply the BANNs framework to six quantitative traits assayed in a heterogenous stock of mice from Wellcome Trust Centre for Human Genetics [47], and two quantitative traits in individuals from the Framingham Heart Study [48]. For the latter, we include an additional study where we independently analyze the same traits in a subset of individuals of European ancestry from the UK Biobank [31].

**Figure 1.**
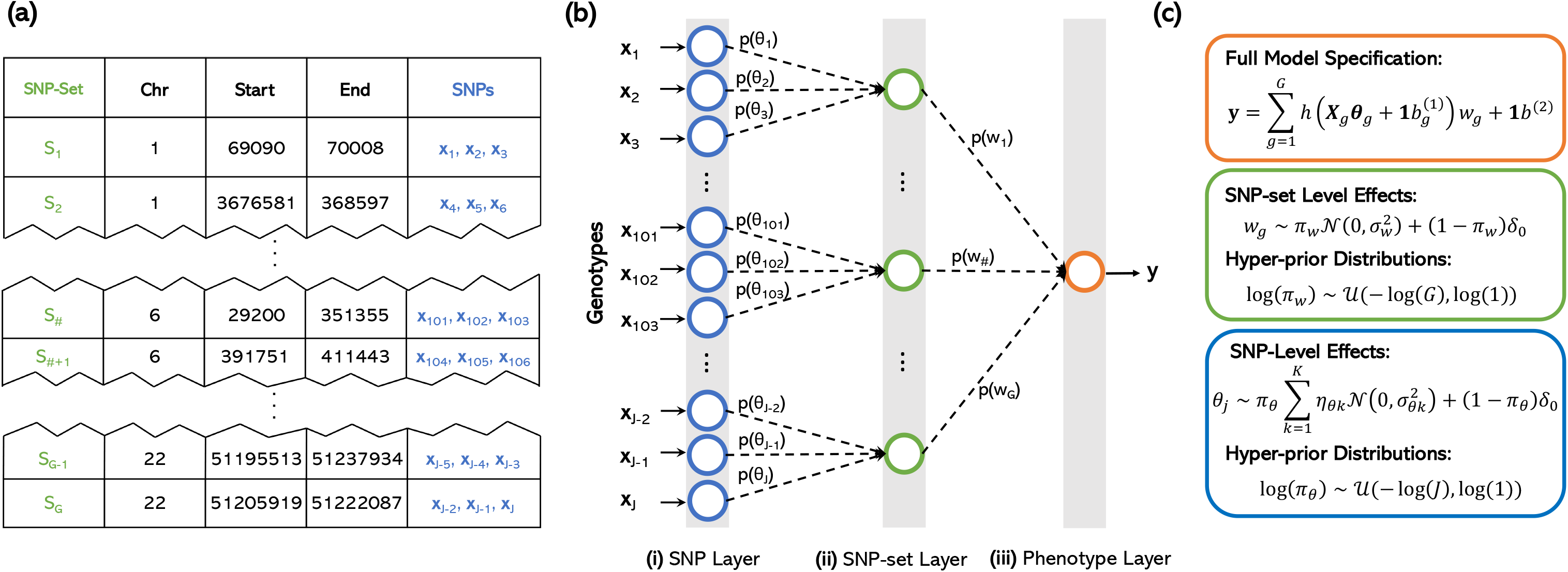
Biologically annotated neural networks (BANNs) allow for efficient multi-scale genotype-phenotype analyses in a unified probabilistic framework by leveraging the hierarchical nature of enrichment studies to define network architecture. **(a)** The BANNs framework requires an *N* × *J* matrix of individual-level genotypes **X** = [**x**_1_,…, **x**_*J*_], an *N*-dimensional phenotypic vector **y**, and a list of *G*-predefined SNP-sets 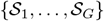. In this work, SNP-sets are defined as genes and intergenic regions (between genes) given by the NCBI’s Reference Sequence (RefSeq) database in the UCSC Genome Browser [51]. **(b)** A partially connected Bayesian neural network is constructed based on the annotated SNP groups. In the first hidden layer, only SNPs within the boundary of a gene are connected to the same node. Similarly, SNPs within the same intergenic region between genes are connected to the same node. Completing this specification for all SNPs gives the hidden layer the natural interpretation of being the “SNP-set” layer. **(c)** The hierarchical nature of the network is represented as nonlinear regression model. The corresponding weights in both the SNP (***θ***) and SNP-set (**w**) layers are treated as random variables with biologically motivated sparse prior distributions. Posterior inclusion probabilities PIP(*j*) ≡ Pr[*θ_j_* ≠ 0 | **y**, **X**] and PIP(*g*) ≡ Pr[*w_g_* ≠ 0 | **y**, **X**, ***θ***_*g*_] summarize associations at the SNP and SNP-set level, respectively. The BANNs framework uses variational inference for efficient network training and incorporates nonlinear processing between network layers for accurate estimation of phenotypic variance explained (PVE).

## Results

### BANNs Framework Overview

Biologically annotated neural networks (BANNs) are feedforward models with partially connected architectures that are inspired by the hierarchical nature of biological enrichment analyses in GWA studies (Fig. 1). The BANNs software takes in one of two data types: (*i*) individual-level data 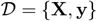 where **X** is an *N* × *J* matrix of genotypes with *J* denoting the number of single nucleotide polymorphisms (SNPs) encoded as {0, 1, 2} copies of a reference allele at each locus and **y** is an *N*-dimensional vector of quantitative traits (Fig. 1a); or (*ii*) GWA summary statistics 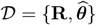 where **R** is a *J* × *J* empirical linkage disequilibrium (LD) matrix of pairwise correlations between SNPs and 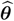 are marginal effect size estimates for each SNP computed using ordinary least squares (OLS) (Fig. S1). In both settings, the BANNs software also requires a predefined list of SNP-set annotations 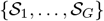 to construct partially connected network layers that represent different scales of genomic units. Structurally, sequential layers of the BANNs model represent different scales of genomic units. The first layer of the network takes SNPs as inputs, with each unit corresponding to information about a single SNP. The second layer of the network represents SNP-sets. All SNPs that have been annotated for the same SNP-set are then connected to the same neuron in the second layer (Fig. 1b).

In this section, we review the hierarchical probabilistic specification of the BANNs framework for individual data; however, note that extensions to summary statistics is straightforward and only requires substituting the genotypes **X** for the LD matrix **R** and substituting the phenotypes **y** for the OLS effect sizes 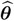 (see Material and Methods). Without loss of generality, let SNP-set *g* represent an annotated collection of SNPs 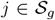 with cardinality 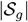. The BANNs framework is probabilistically represented as a nonlinear regression model

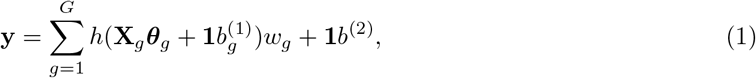

where 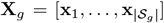 is the subset of SNPs annotated for SNP-set *g*; 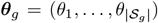 are the corresponding inner layer weights; *h*(•) denotes the nonlinear activations defined for neurons in the hidden layer; **w** = (*w*_1_,…, *w_G_*) are the weights for the *G*-predefined SNP-sets in the hidden layer; 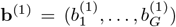 and *b*^(2)^ are deterministic biases that are produced during the network training phase in the input and hidden layers, respectively; and **1** is an *N*-dimensional vector of ones. Here, we define *h*(•) to be a Leaky rectified linear unit (Leaky ReLU) activation function [49], where *h*(***x***) = ***x*** if ***x*** > **0** and 0.01***x*** otherwise. Lastly, for convenience, we assume that the genotype matrix (column-wise) and trait of interest have been mean-centered and standardized.

In this work, we define SNP-sets as collections of contiguous regions of the genome that contain variants within some chromosomal window or neighborhood. More specifically, when studying real mice and human GWA data, we use gene annotations as defined by the Mouse Genome Informatics database [50] and the NCBI’s Reference Sequence (RefSeq) database in the UCSC Genome Browser [51], respectively (Materials and Methods). The BANNs framework flexibly allows for overlapping annotations. In this way, SNPs may be connected to multiple hidden layer units if they are located within the intersection of multiple gene boundaries. SNPs that are unannotated, but located within the same genomic region, are connected to their own units in the second layer and represent the intergenic region between two annotated genes. Given the natural biological interpretation of both layers, the partially connected architecture of the BANNs model creates a unified framework for comprehensibly understanding SNP and SNP-set level contributions to the broad-sense heritability of complex traits and phenotypes. Notably, this framework may be easily extended to other biological annotations and applications.

The framing of the BANNs methodology as a Bayesian nonlinear model helps facilitate our ability to perform classic variable selection (Fig. 1c; see Materials and Methods). Here, we leverage the fact that using nonlinear activation functions for the neurons in the hidden layer implicitly accounts for both additive and non-additive effects between SNPs within a given SNP-set (Supporting Information). Following previous work in the literature [38–42], we treat the weights and connections of the neural network as random variables with prior distributions that reflect how genetic effects are manifested at different genomic scales. For the input layer, we assume that the effect size distribution of non-null SNPs can take vastly different forms depending on both the degree and nature of trait polygenicity [28]. For example, polygenic traits are generated by many mutations of small effect, while other phenotypes can be driven by just a few clusters of SNPs with effect sizes much larger in magnitude [19]. To this end, we place a normal mixture prior on the input layer weights to flexibly estimate a wide range of SNP-level effect size distributions [10, 52–54]

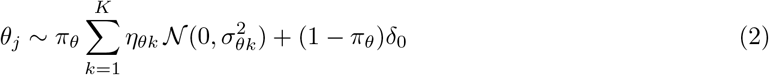

where *δ*_0_ is a point mass at zero; 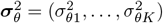 are variance of the *K*-nonzero mixture components; ***η***_*θ*_ = (*η*_*θ*1_,…, *η*_*θK*_) represents the marginal (unconditional) probability that a randomly selected SNP belongs to the *k*-th mixture component such that ∑_*k*_ *η*_*θk*_ = 1; and *π_θ_* denotes the total proportion of SNPs that have a nonzero effect on the trait of interest. Here, we fix *K* = 3 which emulates the hypothesis that SNPs can have large, moderate, and small nonzero effects on phenotypic variation [28]. Similarly, we follow other previous work and assume that enriched SNP-sets contain at least one SNP with a nonzero effect on the trait of interest [26]. This is formulated by placing a spike and slab prior distribution on the weights in the second layer

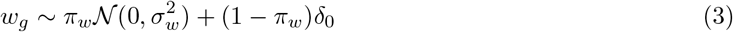

where, in addition to previous notation, *π_w_* denotes the total proportion of SNP-sets that have a nonzero effect on the trait of interest.

By using these point mass mixture distributions in Eqs. (2)–(3), we assume that each connection in the neural network has a nonzero weight with: (*i*) probability *π_θ_* for SNP-to-SNP-set connections, and (*ii*) probability *π_w_* for SNP-set-to-phenotype connections. By modifying a variational inference algorithm assuming point-normal priors in multiple linear regression [55, 56] to the neural network setting, we jointly infer posterior inclusion probabilities (PIPs) for SNPs and SNP-sets. These quantities are defined as the posterior probability that the weight of a given connection in the neural network is nonzero, PIP(*j*) ≡ Pr[*θ_j_* ≠ 0 | **y**, **X**] and PIP(*g*) ≡ Pr[*w_g_* ≠ 0 | **y**, **X**, ***θ**_g_*]. We use this information to prioritize statistically associated SNPs and SNP-sets that significantly contribute to the broad-sense heritability of the trait of interest. With biologically annotated units and the ability to perform statistical inference on explicitly defined parameters, our model presents a fully interpretable extension of neural networks to GWA applications. Additional details and derivations of the BANNs framework can be found in Materials and Methods and Supporting Information.

### Power to Detect SNPs and SNP-Sets in Simulation Studies

In order to assess the performance of models under the BANNs framework, we simulated complex traits under multiple genetic architectures using real genotype data on chromosome 1 from ten thousand randomly sampled individuals of European ancestry in the UK Biobank [31] (see Materials and Methods and previous work [9,28]). After quality control procedures, our simulations included 36,518 SNPs (Supporting Information). Next, we used the NCBI’s Reference Sequence (RefSeq) database in the UCSC Genome Browser [51] to annotate SNPs with the appropriate genes. Unannotated SNPs located within the same genomic region were labeled as being within the “intergenic region” between two genes. Altogether, this left a total of *G* = 2,816 SNP-sets to be included in the simulation study.

After the annotation step, we assume a linear model to generate quantitative traits while varying the following parameters: broad-sense heritability (modestly set to *H*^2^ = 0.2 and 0.6); the proportion of broad-sense heritability that is being contributed by additive effects versus pairwise *cis*-interaction effects (*ρ* =1 and 0.5); and the percentage of enriched SNP-sets that influence the trait (set to 1% for sparse and 10% for polygenic architectures, respectively). We use the parameter *ρ* to assess the neural network’s robustness in the presence of non-additive genetic effects between causal SNPs. To this end, *ρ* =1 represents the limiting case where the variation of a trait is driven by solely additive effects. For *ρ* = 0.5, the additive and pairwise interaction effects are assumed to equally contribute to the phenotypic variance.

In each simulation scenario, we consider traits being generated with and without additional population structure (Materials and Methods, and Supporting Information). To do so, we consider two different data compositions with individuals from the UK Biobank. In the first, we simulate synthetic traits only using individuals who self-identify as being of “white British” ancestry. In the second, we simulate traits by randomly subsampling 3,000 individuals who self-identify as being of “white British” ancestry, 3,000 individuals who self-identify as being of “white Irish” ancestry, and 4,000 individuals who identify as being of “any other white background”. Note that the latter composition introduces additional population structure into the problem. In the main text and Supporting information, we refer to these datasets as the “British” and “European” cohorts, respectively.

Throughout this section, we assess the performance for two versions of the BANNs framework. The first takes in individual-level genotype and phenotype data; while, the second models GWA summary statistics (hereafter referred to as BANN-SS). For the latter, GWA summary statistics are computed by fitting a single-SNP univariate linear model (via ordinary least squares) after quality control to obtain: effect size estimates, standard errors, and *P*-values for all SNPs in the data. We also use the in-sample genotypes to compute the LD matrix between SNPs. All results are based on 100 different simulated phenotypes for each parameter combination (Supporting Information).

The main utility of the BANNs framework is having the ability to detect associated SNPs and enriched SNP-sets simultaneously. Therefore, we compare the performance of BANNs to state-of-the-art SNP and SNP-set level approaches [21, 23, 25–27, 43–46], with the primary idea that our method should be competitive in both settings. For each method, we assess the empirical power and false discovery rates (FDR) for identifying either the correct causal SNPs or the correct SNP-sets containing causal SNPs (Tables S1-S8). Frequentist approaches are evaluated at a Bonferroni-corrected threshold for multiple hypothesis testing (e.g., *P* = 0.05/36518 = 1.37 × 10^−6^ at the SNP-level and *P* = 0.05/2816 = 1.78 × 10^−5^ at the SNP-set level, respectively); while, Bayesian methods are evaluated according to the median probability model (PIPs and posterior enrichment probability ≥ 0.5) [57]. We also compare each method’s ability to rank true positives over false positives via receiver operating characteristic (ROC) and precision-recall curves (Fig. 2 and Figs. S2-S16). Specific results about these analyses are given below.

**Figure 2.**
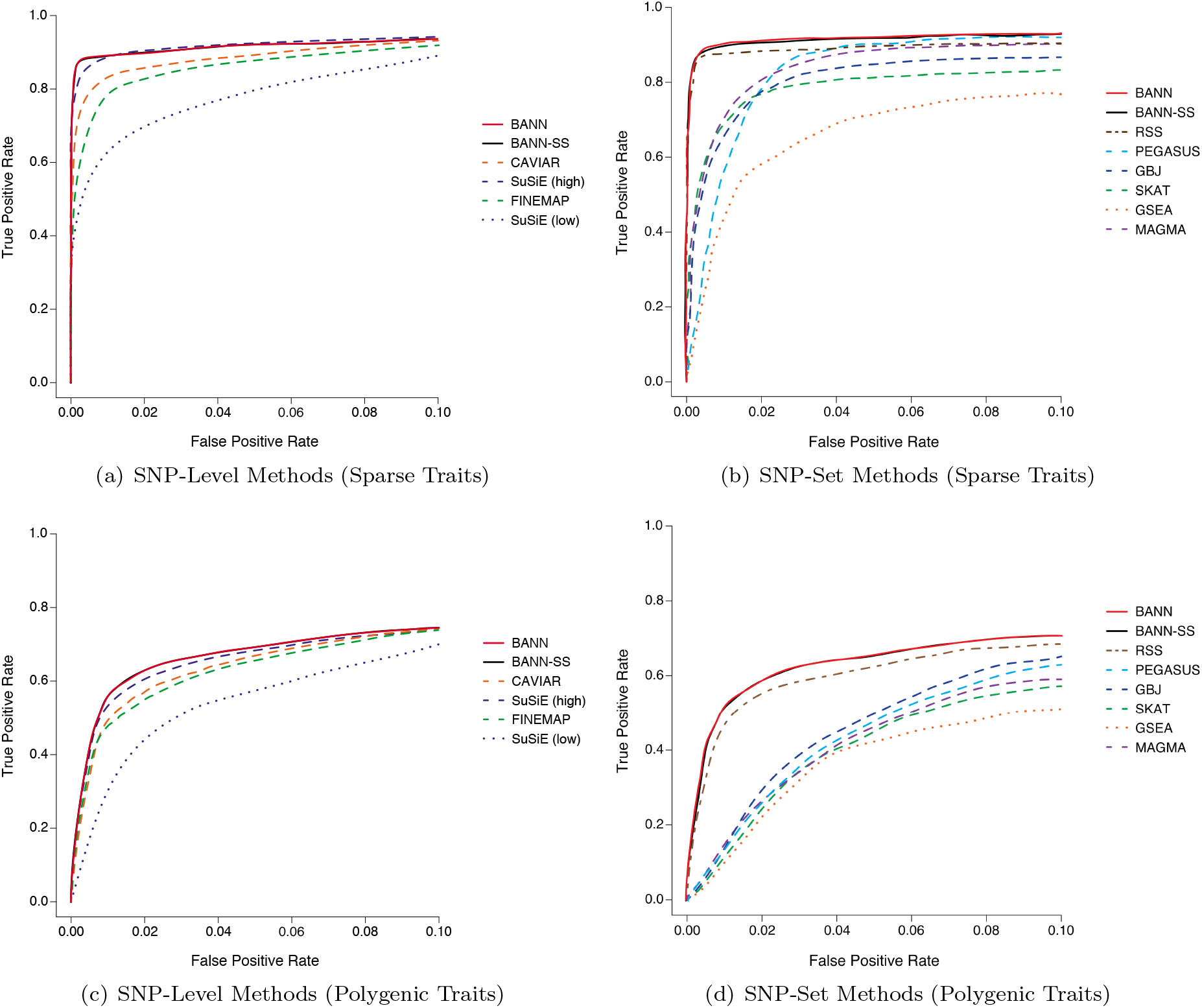
Receiver operating characteristic (ROC) curves comparing the performance of the BANNs (red) and BANN-SS (black) models with competing SNP and SNP-set mapping approaches in simulations (British cohort). Here, quantitative traits are simulated to have broadsense heritability of *H*^2^ = 0.6 with only contributions from additive effects set (i.e., *ρ* =1). We show power versus false positive rate for two different trait architectures: **(a, b)** sparse where only 1% of SNP-sets are enriched for the trait; and **(c, d)** polygenic where 10% of SNP-sets are enriched. We set the number of causal SNPs with nonzero effects to be 1% and 10% of all SNPs located within the enriched SNP-sets, respectively. To derive results, the full genotype matrix and phenotypic vector are given to the BANNs model and all competing methods that require individual-level data. For the BANN-SS model and other competing methods that take GWA summary statistics, we compute standard GWA SNP-level effect sizes and *P*-values (estimated using ordinary least squares). **(a, c)** Competing SNP-level mapping approaches include: CAVIAR [45], SuSiE [46], and FINEMAP [44]. The software for SuSiE requires an input *ℓ* which fixes the maximum number of causal SNPs in the model. We display results when this input number is high (*ℓ* = 3000) and when this input number is low (*ℓ* = 10). **(b, d)** Competing SNP-set mapping approaches include: RSS [26], PEGASUS [25], GBJ [27], SKAT [21], GSEA [43], and MAGMA [23]. Note that the upper limit of the x-axis has been truncated at 0.1. All results are based on 100 replicates (see Supporting Information).

#### Mapped SNP-Level Results

For SNP-level comparisons, we used three fine-mapping methods as benchmarks: CAVIAR [45], SuSiE [46], and FINEMAP [44]. Each of these methods implement Bayesian variable selection strategies, in which different sparse prior distributions are placed on the “true” effect sizes of each SNP and posterior inclusion probabilities (PIPs) are used to summarize their statistical relevance to the trait of interest. Notably, both CAVIAR (exhaustively) and FINEMAP (approximately) search over different models to find the best combination of associated SNPs with nonzero effects on a given phenotype. On the other hand, the software for SuSiE requires an input *ℓ* which fixes the maximum number of causal SNPs to include in the model. In this section, we consider results when this input number is high (*ℓ* = 3000) and when this input number is low (*ℓ* = 10). While SuSiE is applied to individual-level data, both CAVIAR and FINEMAP require summary statistics where marginal z-scores are treated as a phenotype and modeled with in-sample estimate of the LD matrix.

Overall, BANNs, BANN-SS, and SuSiE (with high *ℓ* = 3000) generally achieve the greatest empirical power and lowest FDR across all genetic architectures we considered (Tables S1-S8). These three approaches also stand out in terms of true-versus-false positive rates and precision-versus-recall (Fig. 2 and Figs. S2-S16). Notably, the choice of the *ℓ* parameter largely influenced the performance of SuSiE, as it was consistently the worst performing method when we underestimated the number of causal SNPs with nonzero effects *a priori* (i.e., *ℓ* = 10). Importantly, these performance gains come with a cost: the computational run time of SuSiE becomes much slower as *ℓ* increases (Table S9). For more context, an analysis on just 4,000 individuals and 10,000 SNPs takes the BANNs methods an average of 319 seconds to run on a CPU; while, SuSiE can take up to nearly twice as long to complete as *ℓ* increases (e.g., average runtimes of 23 and 750 seconds for *ℓ* =10 and 3000, respectively).

Training BANNs on individual-level data relatively becomes the best approach when the broad-sense heritability of complex traits is partly made up of pairwise genetic interaction effects between causal SNPs (e.g., *ρ* = 0.5; see Figs. S5-S8 and S13-S16)—particularly when traits have low heritability with polygenic architectures (e.g., *H*^2^ = 0.2). A direct comparison of the PIPs derived by BANNs and SuSiE shows that the proposed neural network training procedure enables the ability to identify associated SNPs even in these more complex phenotypic architectures (Fig. 3 and Figs. S17-S23). It is important to note that the inclusion probabilities were not perfectly calibrated for either BANNs or SuSiE in our simulations (Fig. S24), despite FDR still being reasonably well controlled for both methods (Tables S1-S8). We hypothesize that the quality of PIP calibration for BANNs is a direct consequence of its variational inference algorithm which tends to favor sparse solutions and can lead to greater type II versus type I error rates [46,55]. To investigate how choices in the BANNs model setup contributed to improved variable selection over SuSiE, we also performed an “ablation analysis” [58,59] where we modified parts of the algorithm independently and observed their direct effect on method performance (Fig. S25). Ultimately, these results for BANNs were enabled by a combination of (*i*) using ReLU activation functions in the hidden layers of the BANNs framework, which implicitly enumerates the interactions between SNPs within a given SNP-set, and (*ii*) using model averaging to estimate the inclusion probabilities for the network weights (Supporting Information). Note the absence of the nonlinear activation function only affected the power of BANNs in simulations where there were non-additive genetic effects (e.g., Figs. S25c and S25d).

**Figure 3.**
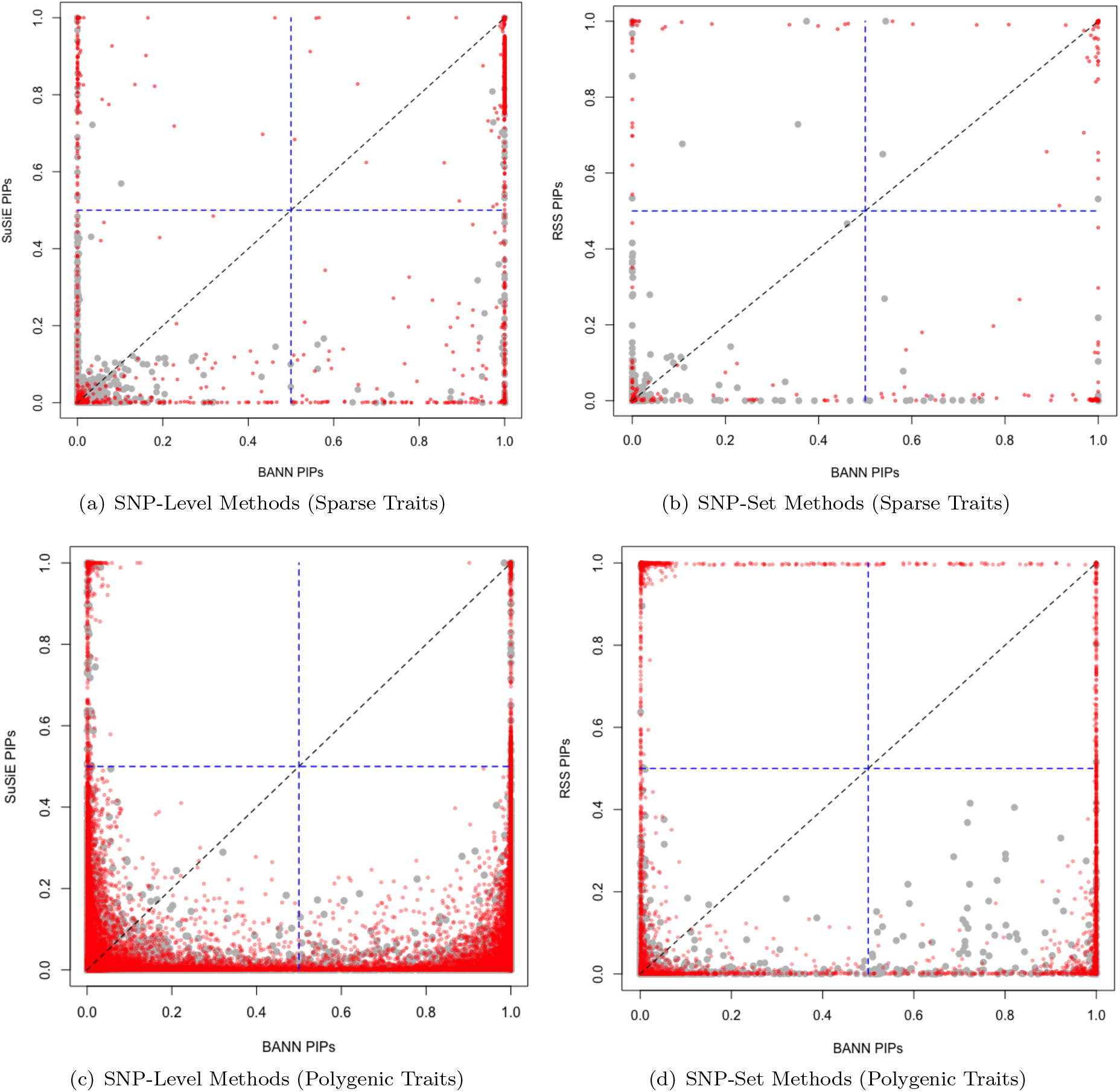
Scatter plots comparing how the integrative neural network training procedure enables the ability to identify associated SNPs and enriched SNP-sets in simulations (British cohort). Quantitative traits are simulated to have broad-sense heritability of *H*^2^ = 0.6 with only contributions from additive effects set (i.e., *ρ* = 1). We consider two different trait architectures: **(a, b)** sparse where only 1% of SNP-sets are enriched for the trait; and **(c, d)** polygenic where 10% of SNP-sets are enriched. We set the number of causal SNPs with nonzero effects to be 1% and 10% of all SNPs located within the enriched SNP-sets, respectively. Results are shown comparing the posterior inclusion probabilities (PIPs) derived by the BANNs model on the x-axis and **(a, c)** SuSiE [46] and **(b, d)** RSS [26] on the y-axis, respectively. Here, SuSie is fit while assuming a high maximum number of causal SNPs (*ℓ* = 3000). The blue horizontal and vertical dashed lines are marked at the “median probability criterion” (i.e., PIPs for SNPs and SNP-sets greater than 0.5) [57]. True positive causal variants used to generate the synthetic phenotypes are colored in red, while non-causal variants are given in grey. SNPs and SNP-sets in the top right quadrant are selected by both approaches; while, elements in the bottom right and top left quadrants are uniquely identified by BANNs and SuSie/RSS, respectively. Each plot combines results from 100 simulated replicates (see Supporting Information).

As a final comparison, the BANN-SS, CAVIAR, and FINEMAP methods see a decline in performance for these same scenarios with genetic interactions. Assuming that the additive and non-additive genetic effects are uncorrelated, this result is also expected since summary statistics are often derived from simple linear additive regression models that (in theory) partition or marginalize out proportions of the phenotypic variance that are contributed by nonlinearities [9,13].

#### Enriched SNP-Set Level Results

For comparisons between SNP-set level methods, we consider six gene or SNP-set enrichment approaches including: RSS [26], PEGASUS [25], GBJ [27], SKAT [21], GSEA [43], and MAGMA [23]. SKAT, VEGAS, and PEGASUS fall within the same class of frequentist approaches, in which SNP-set GWA *P*-values are assumed to be drawn from a correlated chi-squared distribution with covariance estimated using an empirical LD matrix [60]. MAGMA is also a frequentist approach in which gene-level *P*-values are derived from distributions of SNP-level effect sizes using an *F*-test [23]. GBJ attempts to improve upon the previously mentioned methods by generalizing the Berk-Jones statistic to account for complex correlation structures and adaptively adjust the size of annotated SNP-sets to only SNPs that maximize power [61]. Lastly, RSS is a Bayesian linear regression method which places a likelihood on the observed SNP-level GWA effect sizes (using their standard errors and LD estimates), and assumes a spike-and-slab shrinkage prior on the true SNP effects to derive a probability of enrichment for genes or other annotated units [62]. It is worth noting that, while RSS and the BANNs framework are conceptually different, the two methods utilize very similar variational approximation algorithms for posterior inference [55] (Materials and Methods, and Supporting Information).

Similar to the conclusions drawn during the SNP-level assessments, both the BANNs and BANN-SS implementations had among the best tradeoffs between true and false positive rates for detecting enriched SNP-sets across all simulations—once again, including those scenarios which also considered pairwise interactions between causal SNPs (Fig. 2, Figs. S2-S16, and Tables S1-S8). Since RSS is an additive model, it sees a decline in performance for the more complex genetic architectures that we simulated. A direct comparison between the PIPs from BANNs and RSS can be found in Fig. 3 and Figs. S17-S24. Once again, training BANNs on individual-level data becomes the best approach when the broad-sense heritability of complex traits is partly made up of non-additive genetic variation. Our ablation analysis results suggest that the nonlinear activation function plays an important role here (Fig. S25). While RSS also performs generally well for the additive trait architectures, the algorithm for the model often takes twice as long than either of the BANNs implementations to converge (Table S10). PEGASUS, GBJ, SKAT, and MAGMA are score-based methods and, thus, are expected to take the least amount of time to run. BANNs and RSS are hierarchical regression-based methods and the increased computational burden of these approaches results from their need to do (approximate) Bayesian posterior inference. Previous work has suggested that, when using GWA summary statistics to identify genotype-phenotype associations at the SNP-set level, having the ability to adaptively account for possibly inflated SNP-level effect sizes and/or *P*-values is crucial [28]. Therefore, it is understandable why the score-based methods consistently struggle relative to the regression-based approaches even in the simplest simulation cases where traits are generated to have high broad-sense heritability, sparse phenotypic architectures that are dominated by additive genetic effects, and total phenotypic variance that is not confounded by additional population structure (Fig. 2 and Figs. S2-S16). Both the BANN-SS and RSS methods use shrinkage priors to correct for potential inflation in GWA summary statistics and recover estimates that are better correlated with the true generative model for the trait of interest.

### Estimating Total Phenotypic Variance Explained in Simulation Studies

While our main focus is on conducting multi-scale inference of genetic trait architecture, because the BANNs framework provides posterior estimates for all weights in the neural network, we are able to also provide an estimate of phenotypic variance explained (PVE). Here, we define PVE as the total proportion of phenotypic variance that is explained by genetic effects, both additive and non-additive, collectively [16]. Within the BANNs framework, this estimation can be done on both the SNP and SNP-set level while using either genotype-phenotype data or summary statistics (Supporting Information). As a reminder, for our simulation studies, the true PVE is set to *H*^2^ = 0.2 and 0.6, respectively. We assess the ability of BANNs to recover these true estimates using root mean square error (RMSE) (Figs. S26 and S27). In order to be successful at this task, the neural network needs to accurately estimate both the individual effects of causal SNPs in the input layer, as well as their cumulative effects for SNP-sets in the outer layer. BANNs and BANN-SS exhibit the most success with traits have additive sparse architectures (with and without additional population structure)—achieving PVE estimates with RMSEs as low as 4.54 × 10^−3^ and 4.78 × 10^−3^ on the SNP and SNP-set levels for highly heritable phenotypes, respectively. However, both models underestimate the total PVE in polygenic traits and traits with pairwise SNP-by-SNP interactions. Therefore, even though the BANNs framework is still able to correctly prioritize the appropriate SNPs and SNP-sets, in these more complicated settings, we misestimate the approximate posterior means for the network weights and overestimate the variance of the residual training error (Supporting Information). Similar observations have been noted when using variational inference [63,64]. Results from other work also suggest that the sparsity assumption on the SNP-level effects can lead to the underestimation of the PVE [16, 65].

### Mapping Genomic Enrichment in Heterogenous Stock of Mice

We apply the BANNs framework to individual-level genotypes and six quantitative traits in a heterogeneous stock of mice dataset from the Wellcome Trust Centre for Human Genetics [47]. This data contains approximately 2,000 individuals genotyped at approximately 10,000 SNPs—with specific numbers varying slightly depending on the quality control procedure for each phenotype (Supporting Information). For SNP-set annotations, we used the Mouse Genome Informatics database (http://www.informatics.jax.org) [50] to map SNPs to the closest neighboring gene(s). Unannotated SNPs located within the same genomic region were labeled as being within the “intergenic region” between two genes. Altogether, a total of 2,616 SNP-sets were analyzed. The six traits that we consider are grouped based on their category and include: body mass index (BMI) and body weight; percentage of CD8+ cells and mean corpuscular hemoglobin (MCH); and high-density and low-density lipoprotein (HDL and LDL, respectively). We choose to analyze these particular traits because their architectures represent a realistic mixture of the simulation scenarios we detailed in the previous section (i.e., varying different values of *ρ*). Specifically, the mice in this study are known to be genetically related with population structure and these particular traits have been shown to have various levels of broad-sense heritability with different contributions from both additive and non-additive genetic effects [35, 37, 47, 66–68].

For each trait, we provide a summary table which lists the PIPs for SNPs and SNP-sets after fitting the BANNs model to the individual-level genotypes and phenotype data (Tables S11-S16). We use Manhattan plots to visually display the variant-level mapping results across each of the six traits, where chromosomes are shown in alternating colors for clarity and associated SNPs with PIPs above the median probability model threshold are highlighted (Fig. S28). As a comparison, we also report the corresponding SNP and SNP-set level PIPs after running SuSiE [46] and RSS [26] on these same data, respectively. Across all traits, BANNs identified 71 associated SNPs and 57 enriched SNP-sets (according to the median probability model threshold). In comparison, SuSiE identified 22 associated SNPs (11 of which were also identified by BANNs) and RSS identified 14 enriched SNP-sets (6 of which were also identified by BANNs). Importantly, many of the candidate genes and intergenic regions selected by the BANNs model have been previously discovered by past publications as having some functional relationship with the traits of interest (Table 1). For example, BANNs reports the genes *Btbd9* and *hlb156* as being enriched for the percentage of CD8+ cells in mice (PIP = 0.87 and 0.72 versus RSS PIP = 0.02 and 0.68, respectively). This same chromosomal region on chromosome 17 was also reported in the original study as having highly significant quantitative trait loci and contributing non-additive variation for CD8+ cells (bootstrap posterior probability equal to 1.00) [47]. Similarly, the X chromosome is well known to strongly influence adiposity and metabolism in mice [66]. As expected, in body weight and BMI, our approach identified significant enrichment in this region—headlined by the dystrophin gene *Dmd* in both cases [69]. Finally, we note that including intergenic regions in our analyses allows us to discover trait relevant genomic associations outside the immediate gene annotations provided by the Mouse Genome Informatics database. This proved important for BMI where BANNs reported the region between *Gm22219* and *Mc4r* on chromosome 18 as having a relatively high PIP of 0.74 (versus an RSS PIP = 1 × 10^−3^ for reference). Recently, a large-scale GWA study on individuals from the UK Biobank showed that variants around *MC4R* protect against obesity in humans [70].

**Table 1.**
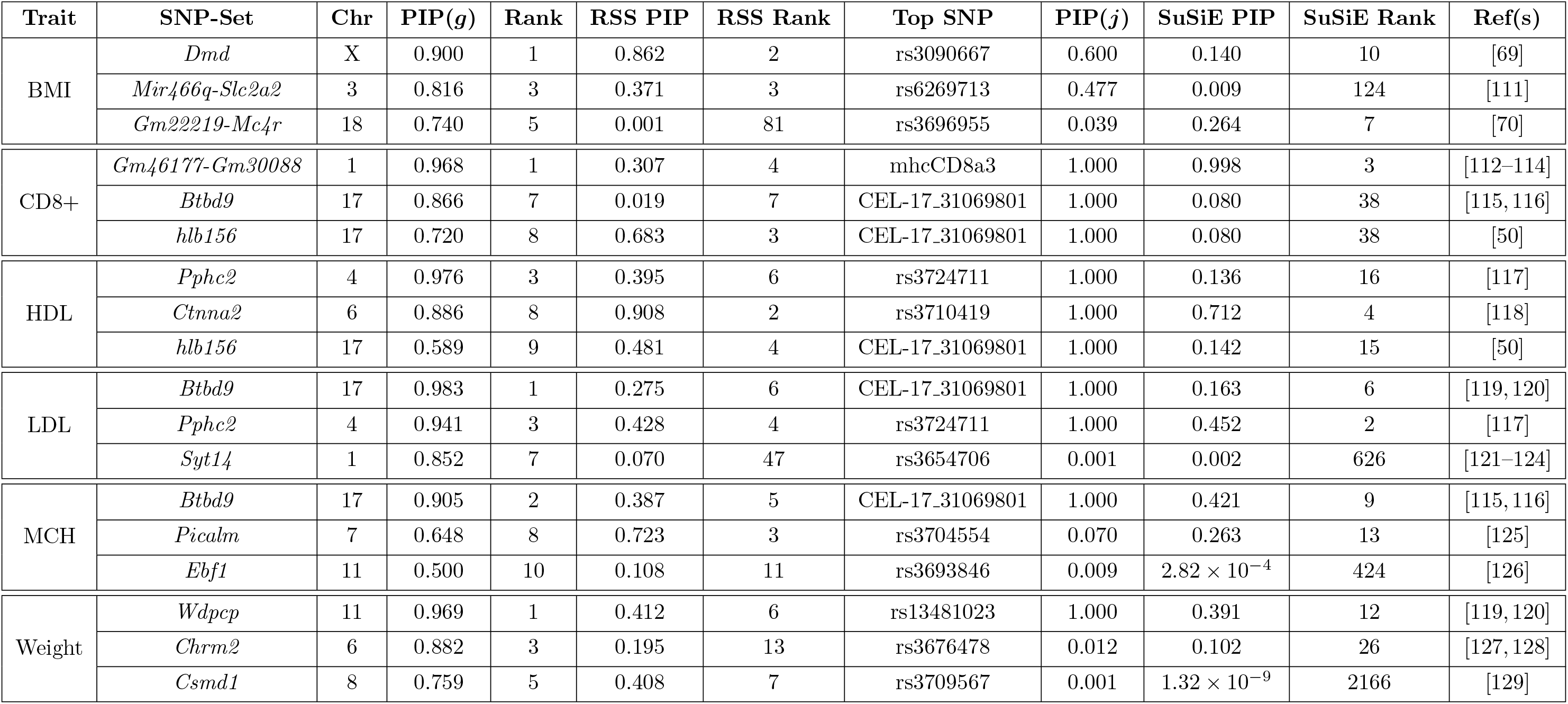
Notable enriched SNP-sets after applying the BANNs framework to six quantitative traits in heterogenous stock of mice from the Wellcome Trust Centre for Human Genetics. [47]. The traits include: body mass index (BMI), percentage of CD8+ cells, high-density lipoprotein (HDL), low-density lipoprotein (LDL), mean corpuscular hemoglobin (MCH), and body weight. Here, SNP-set annotations are based on gene boundaries defined by the Mouse Genome Informatics database (see URLs). Unannotated SNPs located within the same genomic region were labeled as being within the “intergenic region” between two genes. These regions are labeled as *Gene1-Gene2* in the table. Posterior inclusion probabilities (PIP) for the input and hidden layer weights are derived by fitting the BANNs model on individual-level data. A SNP-set is considered enriched if it has a PIP(*g*) ≥ 0.5 (i.e., the “median probability model” threshold [57]). We report the “top” associated SNP within each region and its corresponding PIP(*j*). We also report the corresponding SNP and SNP-set level results after running SuSiE [46] and RSS [26] on these same traits, respectively. The last column details references and literature sources that have previously suggested some level of association or enrichment between the each genomic region and the traits of interest. See Tables S11-S16 for the complete list of SNP and SNP-set level results.

Overall, the results from this smaller GWA study highlight three key characteristics resulting from the sparse probabilistic assumptions underlying the BANNs framework. First, the variational spike and slab prior placed on the weights of the neural network will select no more than a few variants in a given LD block [55]. This is important since traditional naïve SNP-set methods will often exhibit high false positive rates due to many of these correlated regions along the genome [28]. Second, we see that the enrichment of a SNP-set is influenced by the relative posterior distribution of zero and nonzero SNP-level effect sizes within its annotated genomic window (Tables S11-S16). In other words, a SNP-set is not guaranteed to have a high inclusion probability just because it contains one SNP with a large nonzero effect; however, BANNs will report a SNP-set as insignificant if the total ratio of non-causal SNPs within the set heavily outweighs the number of causal SNPs that have been annotated for the same region. To this end, in the presence of large SNP-sets, the BANNs framework will favor preserving false discovery rates at the expense of having slightly more false negatives. Lastly, the careful modeling of the SNP-level effect size distributions and considering genetic interactions enhances our ability to conduct multi-scale genomic inference. In this particular study, we show the power to still find trait relevant SNP-sets with variants that are not marginally strong enough to be detected individually, but have notable genetic signal when their weights are aggregated together (again see Table 1 and Fig. S28).

### Analyzing Lipoproteins in the Framingham Heart Study

Next, we apply the BANNs framework to two continuous plasma trait measurements — high-density lipoprotein (HDL) and low-density lipoprotein (LDL) cholesterol — assayed in 6,950 individuals from the Framingham Heart Study [48] genotyped at 394,174 SNPs genome-wide. Following quality control procedures, we regressed out the top ten principal components of the genotype data from each trait to control for population structure (Supporting Information). Next, we used the gene boundaries listed in the NCBI’s RefSeq database from the UCSC Genome Browser [51] to define SNP-sets. In this analysis, we define genes with boundaries in two ways: (*a*) we use the UCSC gene boundary definitions directly, or (*b*) we augment the gene boundaries by adding SNPs within a ±500 kilobase (kb) buffer to account for possible regulatory elements. Genes with only 1 SNP within their boundary were excluded from either analysis. Unannotated SNPs located within the same genomic region were labeled as being within the “intergenic region” between two genes. Altogether, a total of *G* = 18,364 SNP-sets were analyzed—which included 8,658 intergenic SNP-sets and 9,706 annotated genes—using the UCSC boundaries. When including the 500kb buffer, a total of *G* = 35,871 SNP-sets were analyzed.

For each trait, we again fit the BANNs model to the individual-level genotype-phenotype data and used the median probability model threshold as evidence of statistical significance for all weights in the neural network (Tables S17-S19). We also again report the corresponding SNP and SNP-set level PIPs after running SuSiE and RSS on these same data. Note that while BANNs is run on the genome-wide data jointly, for computational considerations, SuSiE and RSS are run on a chromosome-by-chromosome basis. A complete breakdown of the overlap of findings between BANNs, SuSiE, and RSS can be found on the first page of Table S20. In Fig. 4, we show Manhattan plots of the variant-level association mapping results for BANNs, where each significant SNP is color coded according to its SNP-set annotation. As an additional validation step, we took the enriched SNP-sets identified by BANNs in each trait and used the gene set enrichment analysis tool Enrichr [71,72] to identify the categories that they overrepresent in the database of Genotypes and Phenotypes (dbGaP) and the NHGRI-EBI GWAS Catalog (Figs. S29 and S30). Similar to our results in the previous section, the BANNs framework identified many SNPs and SNP-sets that have been shown to be associated with cholesterol-related processes in past publications (Table 2 with UCSC gene boundary definitions and Table S17 with augmented buffer). For example, in HDL, BANNs identified an enriched intergenic region between the genes *HERPUD1* and *CETP* (PIP = 1.00 versus RSS PIP = 0.78) which has been also replicated in multiple GWA studies with diverse cohorts [73–76]. The Enrichr analyses were also consistent with published results (Figs. S29 and S30). For example, the top ten significant enriched categories in the GWAS Catalog (i.e., Bonferroni-correct threshold *P*-value < 1 × 10^−5^ or *Q*-value < 0.05) for HDL-associated SNP-sets selected by the BANNs model are either directly related to lipoproteins and cholesterol (e.g., “Alpolipoprotein A1 levels”, “HDL cholesterol levels”) or related to metabolic functions (e.g., “Lipid metabolism phenotypes”, “Metabolic syndrome”).

**Figure 4.**
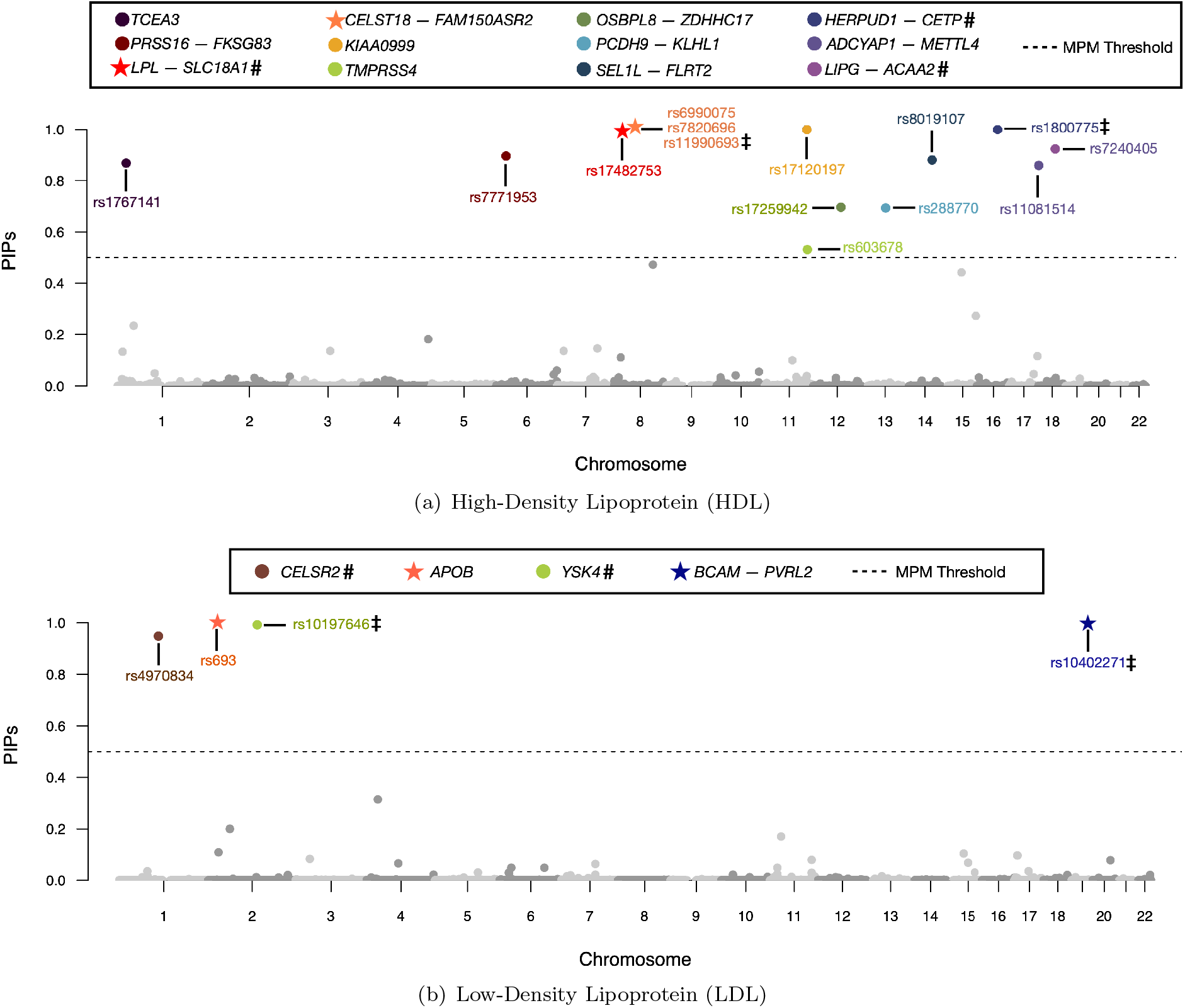
Manhattan plot of variant-level association mapping results for high-density and low-density lipoprotein (HDL and LDL, respectively) traits in the Framingham Heart Study [48]. Posterior inclusion probabilities (PIP) for the neural network weights are derived from the BANNs model fit on individual-level data and are plotted for each SNP against their genomic positions. Chromosomes are shown in alternating colors for clarity. The black dashed line is marked at 0.5 and represents the “median probability model” threshold [57]. SNPs with PIPs above that threshold are color coded based on their SNP-set annotation. Here, SNP-set annotations are based on gene boundaries defined by the NCBI’s RefSeq database in the UCSC Genome Browser [51]. Unannotated SNPs located within the same genomic region were labeled as being within the “intergenic region” between two genes. These regions are labeled as *Gene1-Gene2* in the legend. Double daggers (**‡**) denote SNPs that are also identified when using SuSiE [46] to analyze the same traits, and hashtag symbols (**#**) denote SNP-sets that are identified by RSS [26]. Stars (⋆) denote SNPs and SNP-sets identified by BANNs that replicate in our analyses of HDL and LDL using ten thousand randomly sampled individuals of European ancestry from the UK Biobank [31]. Gene set enrichment analyses for these SNP-sets identified by BANNs can be found in Figs. S29 and S30. A complete list of PIPs for all SNPs and SNP-sets computed in these two traits can be found in Tables S18 and S19. Results for the additional study with the independent UK Biobank dataset [31] are illustrated in Figures S31-S33 and full results are listed in Tables S21 and S22.

**Table 2.**
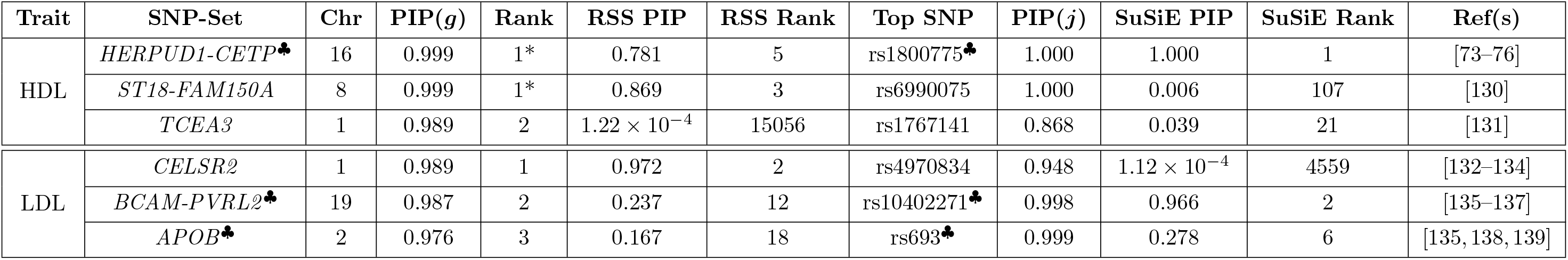
Top three enriched SNP-sets after applying the BANNs framework to high-density and low-density lipoprotein (HDL and LDL, respectively) traits in the Framingham Heart Study [48]. Here, SNP-set annotations are based on gene boundaries defined by the NCBI’s RefSeq database in the UCSC Genome Browser [51]. Unannotated SNPs located within the same genomic region were labeled as being within the “intergenic region” between two genes. These regions are labeled as *Gene1-Gene2* in the table. Posterior inclusion probabilities (PIP) for the input and hidden layer weights are derived by fitting the BANNs model on individual-level data. A SNP-set is considered enriched if it has a PIP(*g*) ≥ 0.5 (i.e., the “median probability model” threshold [57]). We report the “top” associated SNP within each region and its corresponding PIP(*j*). We also report the corresponding SNP and SNP-set level results after running SuSiE [46] and RSS [26] on these same traits, respectively. The last column details references and literature sources that have previously suggested some level of association or enricvariants and pathwayshment between the each genomic region and the traits of interest. See Tables S18 and S19 for the complete list of SNP and SNP-set level results. *: Multiple SNP-sets were tied for this ranking. ♣: SNPs and SNP-sets replicated in an independent analysis of ten thousand randomly sampled individuals of European ancestry from the UK Biobank [31].

As in the previous analysis, the results from this analysis also highlight insight into complex trait architecture enabled by the variational inference used in the BANNs software. SNP-level and SNP-set results remain consistent with the qualitative assumptions underlying our probabilistic hierarchical model. For instance, previous studies have estimated that rs599839 (chromosome 1, bp: 109822166) and rs4970834 (chromosome 1, bp: 109814880) explain approximately 1% of the phenotypic variation in circulating LDL levels [77]. Since these two SNPs are physically closed to each other and sit in a high LD block (*r*^2^ ≈ 0.63 with *P* < 1 × 10^−4^ [78]), the spike and slab prior in the BANNs framework will maintain the nonzero weight for one and penalize the estimated effect of the other. Indeed, in our analysis, rs4970834 was reported to be associated with LDL (PIP = 0.95 versus SuSiE PIP = 1.12 × 10^−4^), while the effect size of rs599839 was shrunk towards 0 (PIP = 1 × 10^−4^ versus SuSiE PIP = 0.99). A similar issue can occur in correctly identifying enriched SNP-sets when nearby sets contain SNPs in tight LD. For example, when augmenting the boundary of SNP-set annotations by a ±500 kilobase buffer, BANNs tends to shrink the PIP of at least one member of overlapping or correlated sets. Due to the variational approximations utilized by BANNs (Materials and Methods, and Supporting Information), if two SNPs or SNP-sets are in strong LD, the model will tend to select just one of them [26, 55].

### Independent Lipoprotein Study using the UK Biobank

To further validate our results from the Framingham Heart Study, we also independently apply BANNs to analyze HDL and LDL cholesterol traits in ten thousand randomly sampled individuals of European ancestry from the UK Biobank [31]. Here, we filter the imputed genotypes from the UK Biobank to keep only the same 394,174 SNPs that were used in the Framingham Heart Study analyses from the previous section. We then apply BANNs, SuSiE, and RSS to the individual-level data and in-sample derived summary statistics using the same (*a*) 18,364 SNP-set annotations based on the NCBI’s RefSeq database from the UCSC Genome Browser [51] and (*b*) 35,849 SNP-sets when applying the augmented ±500 kilobase buffer. It is important to note that we restrict this analysis to just ten thousand individuals due to computational considerations for BANNs and SuSiE since they take in individual level data. In Fig. S31, we show the BANNs variant-level Manhattan plots for the independent UK Biobank cohort with significant SNPs color coded according to their SNP-set annotation. Once again, we use the median probability model threshold to determine statistical significance for all weights in the neural network, and a complete breakdown of the overlap of findings between BANNs, SuSiE, and RSS between the traits can be found in Table S20. Lastly, Tables S21 and S22 give the complete list of all SNP and SNP-set level results in this additional UK Biobank study.

Despite the UK Biobank being a completely independent dataset, we found that BANNs was able to replicate two SNPs and two SNP-sets in HDL and two SNPs and one SNP-set that we observed in the Framingham Heart Study analysis (see specially marked rows in Table 2 and Table S17, as well as the overlap summary given in Table S20). For example, in HDL, both the variants rs1800775 (PIP = 1.00 versus SuSiE PIP = 1.00) and rs17482753 (PIP = 1.00 versus SuSiE PIP = 0.73) were replicated. BANNs also identified the corresponding intergenic region between the genes *HERPUD1* and *CETP* as being enriched (PIP = 1.00 versus RSS PIP = 1.00). In our analysis of LDL, BANNs replicated two out of the four associated SNPs: rs693 within the *APOB* gene, and rs10402271 which falls within the intergenic region between genes *BCAM* and *PVRL2*.

There were a few scenarios where a given SNP-set was replicated but the leading SNP in that region differed between the two studies. For instance, while the intergenic region between *LIPG* and *ACAA2* was enriched in both cohorts, the variant rs7240405 was found to be most associated with HDL in the Framingham Heart Study; a different SNP, rs7244811, was identified in the UK Biobank (Fig. 4 and Fig. S31) Similarly, in the analysis with the ±500 kilobase buffer for SNP-set annotations, rs4939883 in the intergenic region between *LIPG* and *ACAA2* was found to be significant for HDL in the UK Biobank instead of rs7244811 which was selected in the Framingham Heart Study. These discrepancies at the variant level are likely due to: (*i*) the sparsity assumption imposed by BANNs, which lead the model to select one of two variants in high LD; and (*ii*) ancestry differences among individuals from the two studies likely also generate different LD structures in the same genomic region.

As a final step, we took the enriched SNP-sets identified by BANNs in the UK Biobank and used Enrichr [71,72] to ensure that we were still obtaining trait relevant results (Figs. S32 and S33). Indeed, for both HDL and LDL, the most overrepresented categories in dbGaP and the GWAS Catalog (i.e., Bonferroni-correct threshold *P*-value < 1 × 10^−5^ or *Q*-value < 0.05) was consistently the trait of interest— followed by other functionally related gene sets such as “Metabolic syndrome” and “Cholesterol levels”. This story remained largely consistent even when augmenting SNP-set annotations with a ±500 kilobase buffer (Figs. S32 and S33). Overall, the sensible results from performing mapping on the variant-level and enrichment analyses on the SNP-set level in two different independent datasets, only further enhances our confidence about the potential impact of the BANNs framework in GWA studies.

## Discussion

Recently, nonlinear approaches have been applied in biomedical genomics for prediction-based tasks, particularly using GWA datasets with the objective of predicting phenotypes [79–83]. However, since the classical idea of variable selection and hypothesis testing is lost within these statistical algorithms, they have not been widely used for association mapping where the goal is to identify significant SNPs or genes underlying complex traits. Here, we present Biologically Annotated Neural Networks (BANNs): a class of feedforward probabilistic models which overcome this limitation by incorporating partially connected architectures that are guided by predefined SNP-set annotations. This creates an interpretable and integrative framework where the first layer of the neural network encodes SNP-level effects and the neurons within the hidden layer represent the different SNP-set groupings. We frame the BANNs methodology as a Bayesian nonlinear regression model and use sparse prior distributions to perform variable selection on the network weights. By modifying a well established variational inference algorithm, we are able to derive posterior inclusion probabilities (PIPs) which allows researchers to carry out SNP-level mapping and SNP-set enrichment analyses, simultaneously. While we focus on genomic motivations in this study, the concept of partially connected neural networks may extend to any scientific application where annotations can help guide the groupings of variables.

Through extensive simulation studies, we demonstrate the utility of the BANNs framework on individual-level data (Fig. 1) and GWA summary statistics (Fig. S1). Here, we showed that both implementations are consistently competitive with commonly used SNP-level association mapping methods and state-of-the-art SNP-set enrichment methods in a wide range of genetic architectures (Figs. 2–3, Figs. S2-S23, and Tables S1-S8). The advantage of our approach was most clear when the broad-sense heritability of the complex traits included pairwise genetic interactions. In two real GWA datasets, we demonstrated the ability of BANNs to prioritize trait relevant SNPs and SNP-sets that have been identified by previous publications and functional validation studies (Fig. 4, Figs. S28-S30, Tables 1–2, and Tables S11-S19). Lastly, using a third real dataset, we assess the ability of BANNs to statistically replicate a subset of these findings in an independent cohort (Figs. S31-S33 and Tables S21-S22).

The current implementation of the BANNs framework offers many directions for future development and applications. Perhaps the most obvious limitation is that ill-annotated SNP-sets can bias the interpretation of results and lead to misplaced scientific conclusions (i.e., might cause us to highlight the “wrong” gene [84, 85]). This is a common issue in most enrichment methods [28]; however, similar to other hierarchical methods like RSS [26], BANNs is likely to rank SNP-set enrichments that are driven by just a single SNP as less reliable than enrichments driven by multiple SNPs with nonzero effects. Another current limitation for the BANNs model comes from the fact that it uses a variational inference to estimate its parameters. While the current implementation works reasonably well for large datasets (Tables S9 and S10), we showed that our sparse prior assumption combined with the variational expectation-maximization algorithm can lead to slightly miscalibrated PIPs (Figs. S24), underestimated approximations of the PVE (Figs. S27 and S28), and will occasionally miss causal SNPs if they are in high LD with other non-causal SNPs in the dataset. For example, in the application to the Framingham Heart Study, BANNs estimates the PVE for HDL and LDL to be 0.11 and 0.04, respectively. Similarly, in the UK Biobank study, BANNs estimates the PVE for HDL and LDL to be 0.12 and 0.06, respectively. In general, these values are lower than what is typically reported in the literature for these complex phenotypes (PVE ≥ 27% for HDL and PVE ≥ 21% for LDL, respectively) [86]. Exploring different prior assumptions and considering other (more scalable) ways to carry out approximate Bayesian inference is something to consider for future work [87]. For example, the Bayesian sparse linear mixed modeling (BSLMM) framework [16, 88, 89] extends the traditional spike-and-slab prior and could provide a useful, yet alternative, hierarchical specification for BANNs.

There are several other potential extensions for the BANNs framework. First, in the current study, we only consider a single hidden layer based on the annotations of gene boundaries and intergenic region between genes. One natural direction for future work would be to a take more of a deep learning approach by including additional hidden layers to the neural network where genes are grouped based on signaling pathways or other functional ontologies (e.g., transcription factor binding). This would involve integrating information from curated databases such as MSigDB [90,91] or CADD [92]. Second, while BANNs is able to account for nonlinear genetic effects, it cannot be used to directly identify the component (i.e., linear vs. nonlinear) that is driving individual SNP or SNP-set associations. A key part of our future work is learning how to disentangle this information and provide detailed summaries of variant-level and gene-by-gene interaction effects [93]. Third, the current BANNs model only takes in genetic information and, in its current form, ignores unobserved environmental covariates (and potential gene-by-environment or G × E interactions) that explain variation in complex traits. In the future, we would like to expand the framework to also take in covariates as fixed effects in the model. Fourth, we have only focused on analyzing one phenotype at a time in this study. However, many previous studies have extensively shown that modeling multiple phenotypes can often dramatically increase power [94]. Therefore, it would be interesting to extend the BANNs framework to take advantage of phenotype correlations to identify pleiotropic epistatic effects. Modeling strategies based on the multivariate linear mixed model (mvLMM) [95] and matrix variate Gaussian process (mvGP) [96] could be helpful here.

As a final avenue for future work, we only focused on applying BANNs to quantitative traits. For studies interested in extending this approach to binary traits (i.e., case-control studies), one might be tempted to simply place a sigmoid or logistic link function on the penultimate layer of the neural network. Indeed, this would allow the BANNs framework to be expressed as a (nonlinear) logistic classification model which is an approach that has been well-established in the statistics literature [97–99]. Unfortunately, it is not straightforward to define broad-sense heritability under the traditional logistic regression framework. As one alternative, we could implement a penalized quasi-likelihood approach [100] which has been shown to enable effective heritability estimation and differential analyses using the generalized linear mixed model framework. As a second alternative, the liability threshold model avoids issues by assuming that binary traits can be modeled via continuous latent liability scores [101–103]. Therefore, a potentially effective way to extend BANNs to case-control studies would be to develop a two-step algorithmic procedure where: in the first step, we find the posterior mean of the liability scores be using existing software packages and then, in the second step, treat those empirical liability estimates as observed traits in the neural network. Regardless of the modeling strategy, new algorithms are likely needed to maximize the appropriateness of BANNs for non-continuous phenotypes.

## URLs

Biologically annotated neural networks (BANNs) software, https://github.com/lcrawlab/BANNs; UK Biobank, https://www.ukbiobank.ac.uk; Database of Genotypes and Phenotypes (dbGaP), https://www.ncbi.nlm.nih.gov/gap; Framingham Heart Study (FHS), https://www.ncbi.nlm.nih.gov/gap; NHGRI-EBI GWAS Catalog, https://www.ebi.ac.uk/gwas/; UCSC Genome Browser, https://genome.ucsc.edu/index.html; Enrichr software, http://amp.pharm.mssm.edu/Enrichr/; Wellcome Trust Centre for Human Genetics, http://mtweb.cs.ucl.ac.uk/mus/www/mouse/index.shtml; Mouse Genome Informatics database, http://www.informatics.jax.org; CAusal Variants Identification in Associated Regions (CAVIAR) software, http://genetics.cs.ucla.edu/caviar/; Efficient variable selection using summary data from GWA studies (FINEMAP) software, http://www.christianbenner.com; Generalized Berk-Jones (GBJ) test for set-based inference software, https://cran.r-project.org/web/packages/GBJ/; Gene Set Enrichment Analysis (GSEA) software, https://www.nr.no/en/projects/software-genomics; SNP-set (Sequence) Kernel Association Test (SKAT) software, https://www.hsph.harvard.edu/skat; Sum of Single Effects (SuSiE) variable selection software, https://github.com/stephenslab/susieR; Multi-marker Analysis of GenoMic Annotation (MAGMA) software, https://ctg.cncr.nl/software/magma; Precise, Efficient Gene Association Score Using SNPs (PEGASUS) software, https://github.com/ramachandran-lab/PEGASUS; and Regression with Summary Statistics (RSS) enrichment software, https://github.com/stephenslab/rss.

## Acknowledgements

This research was conducted in part using computational resources and services at the Center for Computation and Visualization at Brown University. This research was conducted using the UK Biobank Resource under Application Number 22419. This research was also conducted in part using data and resources from the Framingham Heart Study of the NHLBI and Boston University School of Medicine, which was partially supported by the NHLBI Framingham Heart Study (Contract No. N01-HC-25195) and its contract with Affymetrix, Inc for genotyping services (Contract No. N02-HL-6-4278). We thank all participants and staff from the Framingham Heart Study.

## Author Contributions

LC conceived the methods. PD and WC developed the software and carried out the analyses. All authors wrote and reviewed the manuscript.

## Material and Methods

### Annotations

We used the NCBI’s Reference Sequence (RefSeq) database in the UCSC Genome Browser [51] to annotate SNPs with appropriate SNP-sets. In the main text, we define genes with boundaries in two ways: (*a*) we use the UCSC gene boundary definitions directly, or (*b*) we augment the gene boundaries by adding SNPs within a ±500 kilobase (kb) buffer to account for possible regulatory elements. Genes with only 1 SNP within their boundary were excluded from either analysis. Unannotated SNPs located within the same genomic region were labeled as being within the “intergenic region” between two genes. Altogether, a total of *G* = 28,644 SNP-sets were kept for analysis using the UCSC boundaries and a total of *G* = 35,849 SNP-sets were kept for analysis when including the 500kb buffer.

### Biologically Annotated Neural Networks

Consider a genome-wide association (GWA) study with *N* individuals. We have an *N*-dimensional vector of quantitative traits **y**, an *N* × *J* matrix of genotypes **X**, with *J* denoting the number of single nucleotide polymorphisms (SNPs) encoded as {0, 1, 2} copies of a reference allele at each locus, and a list of *G*-predefined SNP-sets 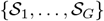 (Fig. 1a). Let each SNP-set *g* represent a known collection of annotated SNPs 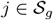 with cardinality 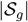. For example, 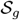 may include SNPs within the regulatory region of a gene. The BANNs framework assumes a partially connected Bayesian neural network architecture based on SNP-set annotations to learn the phenotype of interest for each observation in the data (Fig. 1b). Formally, we specify this network as a nonlinear regression model (Fig. 1c)

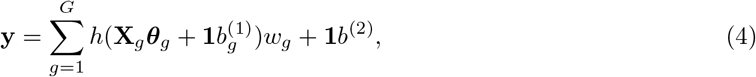

where 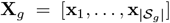 is the subset of SNPs annotated for SNP-set *g*; 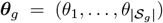 are the corresponding inner layer weights; *h*(●) denotes the nonlinear activations defined for neurons in the hidden layer; **w** = (*w*_1_,…, *w_G_*) are the weights for the *G*-predefined SNP-sets in the hidden layer; 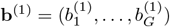 and *b*^(2)^ are deterministic biases that are produced during the network training phase in the input and hidden layers, respectively; and **1** is an *N*-dimensional vector of ones. For convenience, we assume that the genotype matrix (column-wise) and trait of interest have been mean-centered and standardized. In the main text, *h*(●) is defined as a Leaky rectified linear unit (Leaky ReLU) activation function [49], where *h*(***x***) = ***x*** if ***x*** > **0** and 0.01***x*** otherwise. Note that Eq. (4) can be seen as a nonlinear take on classic integrative and structural regression models [22, 26, 104–107] frequently used in GWA analyses.

A key methodological aspect in the BANNs framework is to treat the weights of the input (*θ_j_*) and hidden layers (*w_g_*) as random variables. This, in part, enables us to perform interpretable association mapping on both SNPs and SNP-sets, simultaneously. For the weights on the input layer, our goal is to approximate a wide range of possible SNP-level effect size distributions underlying complex traits. To this end, we assume that SNP-level effects follow a *K*-mixture of normal distributions [10, 52–54]

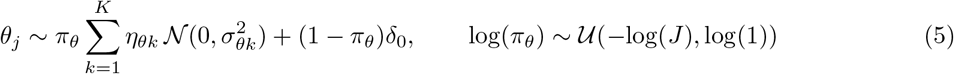

where *δ*_0_ is a point mass at zero; 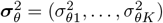 are variance of the *K*-nonzero mixture components; ***η***_*θ*_ = (*η*_*θ*1_,…, *η*_*θK*_) represents the marginal (unconditional) probability that a randomly selected SNP belongs to the *k*-th mixture component such that ∑_*k*_ *η*_*θk*_ = 1; and *π_θ_* denotes the total proportion of SNPs that have a nonzero effect on the trait of interest. We allow sequential fractions of SNPs (*η_θ_*_1_,…, *η_θK_*) to correspond to distinctly smaller effects 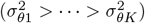 [53]. Intuitively, specifying a larger *K* allows the neural network to learn general SNP effect size distributions spanning over a diverse class of trait architectures. For results in the main text, we fix *K* = 3 for computational reasons. This corresponds to the hypothesis that SNPs can have large, moderate, and small nonzero effects on phenotypic variation [28]. We assume a uniform prior on log *π_θ_* to coincide with the observation that the number of SNPs in each of these categories can vary greatly depending on how heritability is distributed across the genome [16,65] (see Supporting Information).

For inference on the hidden layer, we assume that enriched SNP-sets contain at least one SNP with a nonzero effect. This criterion is formulated by placing a spike and slab prior on the hidden layer weights

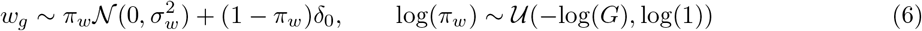

where, in addition to previous notation, the parameter *π_w_* denotes the total proportion of annotated SNP-sets that are enriched for the trait of interest. Given the structural form of the joint likelihood in Eq. (4), the magnitude of association for a SNP-set will be directly influenced by the effect size distribution of the SNPs it contains.

We use a variational Bayesian algorithm to estimate all model parameters (Supporting Information). As the BANNs model is trained, the posterior mean for the weights of non-associated SNP and SNP-sets will trend towards zero as the neural network attempts to identify a subset of neurons that are associated with the phenotype. We use posterior inclusion probabilities (PIPs) as a general summaries of evidence for SNPs and SNP-sets being associated with phenotypic variation. Here, we respectively define

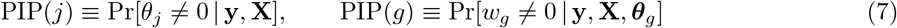

where, again for the latter, the enrichment of SNP-sets is conditioned on the association of individual SNPs. Overall, the Bayesian formulation in the BANNs framework enables network sparsity to be targeted for GWA applications through contextually motivated sparse shrinkage prior distributions in Eqs. (5)–(6). Moreover, posterior inference on PIP(*j*) and PIP(*g*) detail the degree to which nonzero weights occur.

### Posterior Computation with Variational Inference

We combine the likelihood in Eq. (4) and the prior distributions in Eqs. (5)–(7) to perform Bayesian inference. With the size of high-throughput GWA datasets, it is less feasible to implement traditional Markov Chain Monte Carlo (MCMC) algorithms due to the large dimensionality of the parameter space. For model fitting, we modify a previously established variational expectation-maximization (EM) algorithm [55,56] for integrative neural network parameter estimation. The overall goal of variational inference is to approximate the true posterior distribution for network parameters with a “best match” distribution from an approximating family [63]. The EM algorithm we use aims to minimize the Kullback-Leibler divergence between the exact and approximate posterior distributions.

To compute the variational approximations, we make the mean-field assumption that the true posterior can be “fully-factorized” [108]. The algorithm then follows three general steps. First, we assign exchangeable uniform hyper-priors over a grid of values on the log-scale for *π_θ_* and *π_w_* [55]. Next, we iterate through each combination of hyper-parameter values and compute variational updates for the other parameters using co-ordinate ascent. Lastly, we empirically compute (approximate) posterior values for the network connections (***θ***, **w**) and their corresponding inclusion probabilities by marginalizing over the different hyper-parameter combinations. This final step can be viewed as an analogy to Bayesian model averaging where marginal distributions are estimated via a weighted average of conditional distributions multiplied by importance sampling weights [109]. Throughout the model fitting procedure, we assess two different lower bounds for the input and hidden layers to check convergence of the algorithm. The first lower bound is maximized with respect to the SNP-level effects on the observed trait of interest; while, the second lower bound focuses on the SNP-set level enrichments. The software code first iterates over the “inner” lower bound until convergence and then uses those weights to compute the hidden neurons and maximize the “outer” lower bound. Detailed steps in the variational EM algorithm, explicit co-ordinate ascent updates for network parameters, and pseudocode are given in Supporting Information.

Parameters in the variational EM algorithm are initialized by taking a random draws from their assumed prior distributions. Iterations in the algorithm are terminated when either one of two stopping criteria are met: (*i*) the difference between the lower bound of two consecutive updates are within some small range (specified by argument *ϵ*), or (*ii*) a maximum number of iterations is reached. For the simulations and real data analyses ran in this paper, we set *ϵ* = 1 × 10^−4^ for the first criterion and used a maximum of 10,000 iterations for the second.

### Extensions to Summary Statistics

The BANNs framework also models summary statistics in the event that individual-level genotype and phenotype data are not accessible. Here, the software takes alternative inputs: GWA marginal effect size estimates 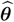 as the response variable, and an empirical linkage disequilibrium (LD) matrix **R** as the design matrix. In the main text, we refer to this version of the method as the BANN-SS model. We assume that GWA summary statistics are derived from the following generative linear model for complex traits

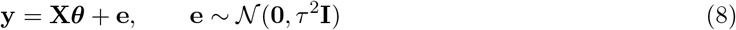

where **e** is a normally distributed error term with mean zero and scaled variance *τ*^2^, and **I** is an *N* × *N* identity matrix. For every *j*-th SNP, the ordinary least squares (OLS) estimates are based on the generative model 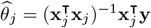, where **x**_*j*_ is the *j*-th column of the individual-level genotype matrix **X** and 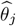 is the *j*-th entry of the vector 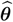. In practice, the LD matrix **R** can be empirically estimated directly from the in-sample GWA study data or from external data (e.g., using an LD map from a population with genomic ancestry similar to individuals in the orginial study). Note that all results presented in the main text are based on estimating **R** with the in-sample genotype data. The BANN-SS model treats the observed OLS estimates and LD matrix as “proxies” for the unobserved phenotype and genotypes, respectively. Specifically, for large sample size *N*, we consider the asymptotic relationship between the expectation of the observed GWA effect size estimates 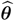 and the true coefficient values ***θ*** is [28, 45, 53, 110]

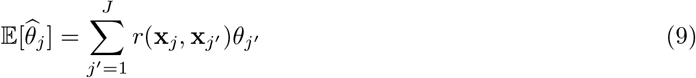

where *r*(**x**_*j*_, **X**_*j*_′) denotes the correlation coefficient between SNPs **x**_*j*_ and **x**_*j*_′. The above resembles a highdimensional regression model with the OLS effect sizes 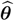 as the response variables, the LD matrix **R** as the design matrix, and the true coefficients ***θ*** being the SNP-level effects that generated the phenotype. Note that this observation is also utilized by other GWA summary-level statistical methods (e.g., CAVIAR [45] and RSS [26,62]). With this relationship in mind, the BANN-SS framework implements the following sparse nonlinear regression for inferring multi-scale genomic effects from summary statistics (Fig. S1)

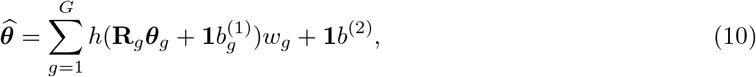

where, in addition to previous notation, **R**_*g*_ is the subset of the LD matrix involving all SNPs annotated for the *g*-th SNP-set. Using the rewritten joint likelihood in Eq. (10), posterior Bayesian inference for the parameters in the BANN-SS model directly mirrors the procedure used when we have access to individual-level data (i.e., as described previously in Eqs. (5)–(7) and given in detail in the Supporting Information). Again, we use measurements PIP(*j*) and PIP(*g*) to summarize whether the true SNP-level effects and aggregated effects on the SNP-set level are statistically associated with the trait of interest.

### Simulation Studies

We implement a simulation scheme to generate quantitative traits under multiple genetic architectures by using real genotype data on chromosome 1 from individuals of European ancestry in the UK Biobank. First, we randomly select a subset of associated SNP-sets (i.e., collections of genomic regions) and assume that complex traits are generated via the linear regression model

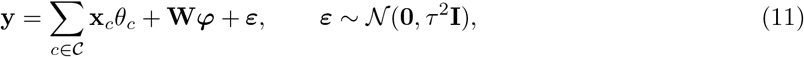

where **y** is an *N*-dimensional vector containing all the phenotypes; 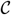 represents the set of causal SNPs contained within the associated SNP-sets; **x**_*c*_ is the genotype for the *c*-th causal SNP encoded as 0, 1, or 2 copies of a reference allele; *θ_c_* is the additive effect size for the *c*-th SNP; **W** is an *N* × *E* matrix which holds all pairwise interactions between the causal SNPs with corresponding effects ***φ***; and ***ε*** is an *N*-dimensional vector of environmental noise. The total phenotypic variance is assumed 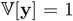. The additive and interaction effect sizes of SNPs in associated SNP-sets are randomly drawn from standard normal distributions and then rescaled so they explain a fixed proportion of the broad-sense heritability 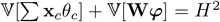. Lastly the environment noise is rescaled such that 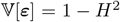. The full genotype matrix and phenotypic vector are given to the BANNs model and all other competing methods that require individual-level data. For the BANN-SS model and other competing methods that take GWA summary statistics, we fit a single-SNP univariate linear model via ordinary least squares (OLS) to obtain: coefficient estimates 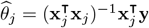, standard errors 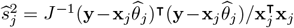, and *P*-values for all SNPs in the data. We also obtain an empirical estimate of the linkage disequilibrium (LD) matrix for these methods **R**, which we compute directly from the full in-sample genotype matrix. Given different model parameters, we simulate data mirroring a wide range of genetic architectures (Supporting Information).

### Data and Software Availability

Source code (with versions in both `R` and `Python` 3) and tutorials for implementing biologically annotated neural networks (BANNs) is publicly available online at https://github.com/lcrawlab/BANNs. All software for competing methods were fit using the default settings, unless otherwise stated in the main text. Links to competing methods, WTCHG mice data, and other relevant sources are also provided (See URLs). Data from the UK Biobank Resource [31] (https://www.ukbiobank.ac.uk) was made available under Application Number 22419. The FHS genotype and phenotype data is available in dbGaP [48] (https://www.ncbi.nlm.nih.gov/gap) with accession number phs000007.

