## Supplementary Text, Figures, and Tables for "Multi-scale Inference of Genetic Trait Architecture using Biologically Annotated Neural Networks"

### 1 Overview of Partially Connected Bayesian Neural Networks

Biologically annotated neural networks (BANNs) are feedforward Bayesian models with have partially connected architectures that are inspired by the hierarchical nature of biological enrichment analyses in GWA studies. The BANNs software takes in one of two data types from genome-wide association (GWA) studies: ($i$) individual-level data $\mathcal{D} = \{\mathbf{X}, \mathbf{y}\}$ where $\mathbf{X}$ is an $N \times J$ matrix of genotypes with $J$ denoting the number of single nucleotide polymorphisms (SNPs) encoded as $\{0, 1, 2\}$ copies of a reference allele at each locus and $\mathbf{y}$ is an $N$-dimensional vector of quantitative traits (see Fig. 1 in the main text); or ($ii$) GWA summary statistics $\mathcal{D} = \{\mathbf{R}, \widehat{\boldsymbol{\theta}}\}$ where $\mathbf{R}$ is a $J \times J$ empirical linkage disequilibrium (LD) matrix of pairwise correlations between SNPs and $\widehat{\boldsymbol{\theta}}$ are marginal effect size estimates for each SNP computed using ordinary least squares (OLS) (see Fig. S1). In either setting, the BANNs software also requires a predefined list of SNP-set annotations $\{\mathcal{S}_1, \ldots, \mathcal{S}_G\}$ to construct partially connected network layers that represent different scales of genomic units. In this section, we review the hierarchical probabilistic specification of the BANNs framework for individual data; however, note that extensions to summary statistics is straightforward and only requires substituting the genotypes $\mathbf{X}$ for the LD matrix $\mathbf{R}$ and substituting the phenotypes $\mathbf{y}$ for the OLS effect sizes $\widehat{\boldsymbol{\theta}}$.

Without loss of generality, let SNP-set $g$ represent an annotated collection of SNPs $j \in \mathcal{S}_g$ with cardinality $|\mathcal{S}_g|$. The BANNs framework is probabilistically represented as a nonlinear regression model

$$\mathbf{y} = \sum_{g=1}^{G} h(\mathbf{X}_g \boldsymbol{\theta}_g + \mathbf{1} b_g^{(1)}) w_g + \mathbf{1} b^{(2)}, \tag{1}$$

where $\mathbf{X}_g = [\mathbf{x}_1, \ldots, \mathbf{x}_{|\mathcal{S}_g|}]$ is the subset of SNPs annotated for SNP-set $g$; $\boldsymbol{\theta}_g = (\theta_1, \ldots, \theta_{|\mathcal{S}_g|})$ are the corresponding inner layer weights; $h(\bullet)$ denotes the nonlinear activations defined for neurons in the hidden layer; $\mathbf{w} = (w_1, \ldots, w_G)$ are the weights for the $G$-predefined SNP-sets in the hidden layer; $\mathbf{b}^{(1)} = (b_1^{(1)}, \ldots, b_G^{(1)})$ and $b^{(2)}$ are deterministic biases that are produced during the network training phase in the input and hidden layers, respectively; and $\mathbf{1}$ is an $N$-dimensional vector of ones. For convenience, we assume that the genotype matrix (column-wise) and trait of interest have been mean-centered and standardized. In the main text, $h(\bullet)$ is defined as a Leaky rectified linear unit (Leaky ReLU) activation function [1], where $h(\boldsymbol{x}) = \boldsymbol{x}$ if $\boldsymbol{x} > 0$ and $0.01\boldsymbol{x}$ otherwise. Throughout this Supporting Information, we will equivalently write Eq. (1) in matrix notation as

$$\mathbf{y} = \mathbf{H}(\boldsymbol{\theta})\mathbf{w} + \mathbf{1} b^{(2)},$$

where $\mathbf{H}(\boldsymbol{\theta}) = [h(\mathbf{X}_1\boldsymbol{\theta}_1 + \mathbf{1}b_1^{(1)}), \ldots, h(\mathbf{X}_G\boldsymbol{\theta}_G + \mathbf{1}b_G^{(1)})]$ denotes the matrix of nonlinear neurons in the hidden layer which are empirically computed given estimates the input layer weights. The hierarchical structure of the joint likelihood can be seen as a nonlinear take on classical integrative and structural regression models frequently used in GWA analyses [2–8].

As explained in the main text, we treat the weights of the input ($\boldsymbol{\theta}$) and hidden layers ($\mathbf{w}$) as random variables which allows for multi-scale genomic inference on both SNPs and SNP-sets, simultaneously. We assume that SNP-level effects follow a sparse $K$-mixture of normal distributions

$$\theta_j \sim \pi_\theta \sum_{k=1}^{K} \eta_{\theta k} \mathcal{N}(0, \sigma_{\theta k}^2) + (1 - \pi_\theta)\delta_0 \tag{2}$$

where $\delta_0$ is a point mass at zero; $\boldsymbol{\sigma}_\theta^2 = (\sigma_{\theta 1}^2, \ldots, \sigma_{\theta K}^2)$ are variance of the $K$ nonzero mixture components; $\boldsymbol{\eta}_\theta = (\eta_{\theta 1}, \ldots, \eta_{\theta K})$ represents the marginal (unconditional) probability that a randomly selected SNP belongs to the $k$-th mixture component such that $\sum_k \eta_{\theta k} = 1$; and $\pi_\theta$ denotes the total proportion of SNPs that have a nonzero effect on the trait of interest. Notice that we write the mixture prior in this form as a way to simplify updates in the algorithm for posterior inference. For reference, one can think of the delta mass as a normal distribution with fixed variance set to zero. Intuitively, specifying a larger $K$ allows the neural network to learn general SNP effect size distributions spanning over a diverse class of trait architectures. For example, one can take a nonparametric approach and allow $K \to \infty$ such that Eq. (2) mirrors a Dirichlet process Gaussian mixture [9]. For results in the main text, we follow previous work and fix $K = 3$ [10–12]. This corresponds to the general hypothesis that SNPs can have large, moderate, and small nonzero effects on phenotypic variation [13]. For inference on the hidden layer, we assume that enriched SNP-sets contain at least one SNP with a nonzero effect. This simpler criterion is formulated by placing a spike and slab prior on the hidden weights

$$w_g \sim \pi_w \mathcal{N}(0, \sigma_w^2) + (1 - \pi_w)\delta_0. \tag{3}$$

where, due to the integrative form of the likelihood in Eq. (1), the magnitude of association for a SNP-set will be directly influenced by the effect size distribution of the SNPs it contains.

For the hyper-parameters in the model, we assume the following prior distributions

$$\log(\pi_\theta) \sim \mathcal{U}(-\log(J), \log(1)), \qquad \log(\pi_w) \sim \mathcal{U}(-\log(G), \log(1)). \tag{4}$$

Following previous work, relatively uninformative uniform priors are assumed over $\log \pi_\theta$ and $\log \pi_w$ to reflect our lack of knowledge *a priori* about the proportion on associated SNP and SNP-sets with nonzero weights [14–16]. To facilitate posterior computation and interpretable inference, we also introduce two vectors of binary indicator variables $\boldsymbol{\gamma}_\theta = (\gamma_{\theta 1}, \dots, \gamma_{\theta J}) \in \{0,1\}^J$ and $\boldsymbol{\gamma}_w = (\gamma_{w1}, \dots, \gamma_{wG}) \in \{0,1\}^G$ where we implicitly assume *a priori* that

$$\Pr[\gamma_{\theta j} = 1] = \Pr[\theta_j \neq 0] = \pi_\theta, \qquad \Pr[\gamma_{wg} = 1] = \Pr[w_g \neq 0] = \pi_w. \tag{5}$$

Alternatively, we say $\gamma_{\theta j}$ and $\gamma_{wg}$ take values of 1 when weights $\theta_j$ and $w_g$ are drawn from the normal "slabs" of Eqs. (2) and (3), respectively; they take values of 0 otherwise. In the main text, we refer to these indicators as inclusion probabilities [17] and we use the marginal posterior means of these quantities as general summaries of evidence that SNPs and SNP-sets are statistically associated with phenotypic variation.

### 2 Variational Expectation-Maximization (EM) Algorithm

We modify a previously developed variational expectation-maximization (EM) algorithm to estimate the posterior distribution of parameters in the BANNs framework. The derivations in this section largely follow those developed in previous work [7, 9, 18–20]. As mentioned in the main text, the overall goal of variational inference is to approximate the true posterior distribution for network parameters with a similar distribution from an approximating family [21–25]. The EM algorithm we use aims to minimize the Kullback-Leibler divergence between the exact and approximate posterior distributions, respectively. To begin, we assign exchangeable uniform hyper-priors over a grid of values on the log-scale for $\pi_\theta$ and $\pi_w$ [15]. We then run the EM algorithm while iterating through each combination of these values. In the E-step, we use co-ordinate ascent to update the free parameters of the approximate variational posterior. In the M-step, we derive updates for the model hyper-parameters by solving for the roots of their gradients. Finally, in the last step, we empirically compute (approximate) posterior values for the network connection weights $(\boldsymbol{\theta}, \mathbf{w})$ and their corresponding inclusion probabilities by marginalizing over the different model combinations for $\pi_\theta$ and $\pi_w$ with normalized importance weights [19, 20]. A complete overview of the algorithm is given below. Again, note that extensions to summary statistics is straightforward and only requires substituting the genotypes $\mathbf{X}$ for an LD matrix $\mathbf{R}$ and substituting the phenotypes $\mathbf{y}$ for estimated OLS effect sizes $\widehat{\boldsymbol{\theta}}$ (see Material and Methods in the main text).

Given the formulation of the BANNs model and the partially connected neural network architecture, the weights in the second layer are conditionally independent of the weights in the input layer given the activations (or outputs) from the first layer. This means that we can break up the model fitting procedure into two integrative parts and assess two different lower bounds for the input and hidden layer weights, respectively, to ensure convergence. Specifically estimates on the SNP-level are first maximized with respect to the trait of interest; while, parameters corresponding to the SNP-set level are maximized with respect to the observed trait. The software code iterates between the "inner" lower bound and the "outer" lower bound each step of the algorithm until convergence. Iterations in the algorithm are terminated when either one of two stopping criteria are met: ($i$) the difference between the lower bound of two consecutive updates are within some small range (specified by tolerance argument $\epsilon$), or ($ii$) a maximum number of iterations is reached. For the simulations and real data analyses ran in this paper, we set $\epsilon = 1 \times 10^{-4}$ for the first criterion and used a maximum of 10,000 iterations for the second.

#### 2.1 Input Layer (SNP-Level) Updates

For the SNP-level effects in the input layer, we aim to find a distribution $q(\boldsymbol{\theta}, \boldsymbol{\gamma}_\theta)$ that approximates the true posterior distribution $p(\boldsymbol{\theta}, \boldsymbol{\gamma}_\theta \,|\, \mathcal{D})$, where $\boldsymbol{\theta} = (\boldsymbol{\theta}_1, \ldots, \boldsymbol{\theta}_G)$ and $\mathcal{D}$ is used to denote the individual-level data and all relevant hyper-parameters. The similarity between these two distributions is maximized by minimizing the Kullback-Leibler (KL) divergence between them. This is formulated by

$$\mathrm{KL}(q(\boldsymbol{\theta}, \boldsymbol{\gamma}_\theta) \,\|\, p(\boldsymbol{\theta}, \boldsymbol{\gamma}_\theta \,|\, \mathcal{D})) = \int \log \left[ \frac{q(\boldsymbol{\theta}, \boldsymbol{\gamma}_\theta)}{p(\boldsymbol{\theta}, \boldsymbol{\gamma}_\theta \,|\, \mathcal{D})} \right] q(\boldsymbol{\theta}, \boldsymbol{\gamma}_\theta) \, \mathrm{d}\boldsymbol{\theta} \, \mathrm{d}\boldsymbol{\gamma}_\theta. \tag{6}$$

Once again, to facilitate posterior inference on this layer, we follow previous work [9] by introducing another vector of binary indicator variables $\varphi_{jk} \in \{0, 1\}$ to indicate which of the $K$-normal components $\theta_j$ belongs to in the prior specified in Eq. (2) such that $\Pr[\varphi_{jk} = 1] = \eta_{\theta k}$. This leads to the following natural expression for the variational mixture distribution on each of the individual SNP-level weights

$$q(\theta_j, \gamma_{\theta j}; \phi_j) = \begin{cases} \sum_{k=1}^{K} \alpha_{jk} \, \mathcal{N}(m_{jk}, s_{jk}^2) & \text{if} \quad \gamma_{\theta j} = 1 \\ (1 - \sum_{k=1}^{K} \alpha_{jk}) \, \delta_0 & \text{if} \quad \gamma_{\theta j} = 0 \end{cases} \tag{7}$$

where, in addition to previous notation, we assume $q[\varphi_{jk} = 1] = \alpha_{jk}$ for the binary indicators. Additionally, we will let $\phi_j = \{\alpha_{jk}, m_{jk}, s_{jk}^2\}_{k=1}^K$ denote a collection of free parameters and $\boldsymbol{\phi} = (\phi_1, \ldots, \phi_J)$ will be used to compute the approximations [26, 27]. The basic idea behind the variational approximation is to formulate a lower bound to the marginal likelihood, then to iteratively adjust the free parameters in $\boldsymbol{\phi}$ so that this bound becomes as tight as possible [19, 20, 25]. Finding the "best" factorized variational distribution amounts to finding the free parameters $\boldsymbol{\phi}$ that make the Kullback-Leibler divergence in Eq. (6) as small as possible. Taking moments with respect to the specific class of variational distributions in Eq. (7) yields the following analytical expression for the lower bound on the inner layer (or SNP-level)

$$\begin{aligned} \mathrm{LB}(\pi_\theta, \boldsymbol{\sigma}_\theta^2, \tau_\theta^2) = &-\frac{N}{2}\log(2\pi\tau_\theta^2) - \frac{1}{2\tau_\theta^2}\|\mathbf{y} - \mathbf{X}\boldsymbol{\beta}_\theta\|_2^2 - \frac{1}{2}\sum_{j=1}^{J}(\mathbf{X}^\intercal\mathbf{X})_{jj}\mathbb{V}[\theta_j] \\ &- \sum_{j=1}^{J}\sum_{k=1}^{K}\alpha_{jk}\log\left(\frac{\alpha_{jk}}{\pi_\theta}\right) + \frac{1}{2}\sum_{j=1}^{J}\sum_{k=1}^{K}\alpha_{jk}\left[1 + \log\left(\frac{s_{jk}^2}{\tau_\theta^2\sigma_{\theta k}^2}\right) - \frac{s_{jk}^2 + m_{jk}^2}{\tau_\theta^2\sigma_{\theta k}^2}\right] \end{aligned} \tag{8}$$

where $\tau_\theta^2 \approx \mathbb{V}[\mathbf{y} - \mathbf{X}\boldsymbol{\beta}_\theta]$ approximates the variance of residual training error in the input layer; $\| \bullet \|_2$ is the Euclidean norm; $\boldsymbol{\beta}_\theta$ is a $J$-dimensional estimate of the posterior mean for $\boldsymbol{\theta}$ with individual elements $\beta_{\theta j} = \sum_{k=1}^K \alpha_{jk} m_{jk}$; the term $(\mathbf{X}^\intercal\mathbf{X})_{jj}$ is the $j$-th diagonal component of the matrix $(\mathbf{X}^\intercal\mathbf{X})$; and $\mathbb{V}[\theta_j] = \sum_{k=1}^K \alpha_{jk}(m_{jk}^2 + s_{jk}^2) - (\sum_{k=1}^K \alpha_{jk} m_{jk})^2$ is the variance of the $j$-th weight under the approximating distribution in Eq. (7). We now describe the expectation and maximization steps of the approximate EM algorithm below (see Software Details in Supporting Information, Section 10). Again, the derivations in this section largely follow those developed in previous work and other theoretical derivations can be found in those corresponding sources [7, 9, 18–20].

1. **E-Step: Update the Variational Free Parameters.** As done in Carbonetto and Stephens (2012) [19], for the E-step of the algorithm, we take partial derivatives of the variational lower bound in Eq. (8) with respect to the free parameters and conditioned on hyperparameters $(\pi_\theta, \boldsymbol{\sigma}_\theta^2, \tau_\theta^2)$. Next, we set these partial derivatives to zero and solve for $m_{jk}$, $s_{jk}^2$, and $\alpha_{jk}$ which yields

$$m_{jk} = \frac{s_{jk}^2}{\tau_\theta^2}\left[(\mathbf{X}^\intercal\mathbf{y})_j - \sum_{l\neq j}(\mathbf{X}^\intercal\mathbf{X})_{jl}\beta_{\theta l}\right] \tag{9}$$

$$s_{jk}^2 = \tau_\theta^2\left[(\mathbf{X}^\intercal\mathbf{X})_{jj} + \frac{1}{\sigma_{\theta k}^2}\right]^{-1} \tag{10}$$

$$\alpha_{jk} = \text{Sigmoid}\left(\log\left(\frac{\pi_\theta}{1-\pi_\theta}\right) + \log\left(\frac{s_{jk}}{\sigma_{\theta k}\tau_\theta}\right) + \frac{m_{jk}^2}{2s_{jk}^2}\right) \tag{11}$$

where $\sum_k \alpha_{jk} \approx \Pr[\gamma_{\theta j} = 1 \,|\, \mathbf{y}, \mathbf{X}, \pi_\theta, \boldsymbol{\sigma}_\theta^2, \tau_\theta^2]$ and the sigmoid function is set to be the standard logistic function. Intuitively, the E-step of the algorithm for the input layer produces a collection of $\boldsymbol{\alpha}_j = (\alpha_{j1}, \ldots, \alpha_{jK})$ values to determine whether each SNP has a nonzero effect on the phenotypic variance.

2. **M-Step: Update the Variance Hyper-Parameters.** In the M-step of the algorithm, we fix the newly estimated values of the variational free parameters $\boldsymbol{\phi}$ from the E-step and maximize the lower bound in Eq. (8) with respect to $\sigma_{\theta k}^2$ and $\tau_\theta^2$. This done in the usual way of solving for roots $\sigma_{\theta k}^2$ and $\tau_\theta^2$ of the gradient which yields [20]

$$\sigma_{\theta k}^2 = \left[\sum_{j=1}^J \alpha_{jk}(m_{jk}^2 + s_{jk}^2)\right] \Bigg/ \left(\tau_\theta^2 \sum_{j=1}^J \alpha_{jk}\right) \tag{12}$$

$$\tau_\theta^2 = \left(N + \sum_{j=1}^J\sum_{k=1}^K \alpha_{jk}\right)^{-1}\left[\|\mathbf{y} - \mathbf{X}\boldsymbol{\beta}_\theta\|_2^2 + \sum_{j=1}^J (\mathbf{X}^\intercal\mathbf{X})_{jj}\mathbb{V}[\theta_j] + \sum_{j=1}^J\sum_{k=1}^K \frac{\alpha_{jk}}{\sigma_{\theta k}^2}(m_{jk}^2 + s_{jk}^2)\right] \tag{13}$$

where $N$ is equal to the dimensionality of the trait vector (i.e., the sample size when modeling individual-level data).

Following previous work [7,14–16,19,20], we incorporate Eq. (4) and account for our lack of *a priori* knowledge about the "correct" proportion of associated SNPs with nonzero effects by placing an exchangeable uniform hyper-prior distribution over an $L$-valued grid of possible values where $\{\pi_\theta^{(1)}, \ldots, \pi_\theta^{(L)}\} \in [1/J, 1]$. We then use the lower bound to the likelihood in Eq. (8) to approximate the posterior distribution of $\pi_\theta$. Formally, we approximate $\Pr[\pi_\theta = \pi_\theta^{(l)} \,|\, \mathbf{y}, \mathbf{X}]$ with the normalized importance weights

$$\lambda_\theta^{(l)} = \frac{\text{LB}(\pi_\theta^{(l)}, \boldsymbol{\sigma}_\theta^2, \tau_\theta^2)}{\sum_{l'=1}^L \text{LB}(\pi_\theta^{(l')}, \boldsymbol{\sigma}_\theta^2, \tau_\theta^2)}. \tag{14}$$

As a final step in the model fitting procedure, we empirically compute (approximate) SNP-level posterior inclusion probabilities by marginalizing over the different grid combinations for $\boldsymbol{\pi}_\theta$. Namely,

$$\text{PIP}(j) \equiv \Pr[\gamma_{\theta j} = 1 \,|\, \mathbf{y}, \mathbf{X}] \approx \sum_{l=1}^L \lambda_\theta^{(l)} \Pr[\gamma_{\theta j} = 1 \,|\, \mathbf{y}, \mathbf{X}, \pi_\theta^{(l)}, \boldsymbol{\sigma}_\theta^2, \tau_\theta^2]. \tag{15}$$

This final step can be viewed as an analogy to Bayesian model averaging where marginal distributions are estimated via a weighted average of conditional distributions multiplied by importance weights [19,20,28].

#### 2.2 Outer Layer (SNP-Set Level) Updates

In this section, we detail the posterior computation for parameters in the outer layer of the partially connected neural network. We are now interested in finding a distribution $q(\mathbf{w}, \boldsymbol{\gamma}_w)$ that approximates the true posterior $p(\mathbf{w}, \boldsymbol{\gamma}_w \mid \mathcal{D}, \boldsymbol{\theta})$. Here, it is important to note that, due to the integrative setup of the joint likelihood used in the BANNs framework, the true posterior for the weights in the outer layer is conditionally dependent upon the posterior estimates for the weights in the input layer. Since we assume that enriched SNP-sets contain at least one SNP with a nonzero effect, we consider a simpler family of variational distributions

$$q(w_g, \gamma_{wg}; \psi_g) = \begin{cases} \alpha_g \, \mathcal{N}(m_g, s_g^2) & \text{if} \quad \gamma_{wg} = 1 \\ (1-\alpha_g)\, \delta_0 & \text{if} \quad \gamma_{wg} = 0 \end{cases} \tag{16}$$

where $\psi_g = (\alpha_g, m_g, s_g^2)$ is used to describe a new set free parameters for the $g$-th SNP-set. Once again, our goal is to find the "best" factorized variational distribution amounts with free parameters $\boldsymbol{\psi}$ that minimize the Kullback-Leibler divergence between the exact and approximate posteriors. The specific class of variational distributions in Eq. (16) yields the following analytical expression for the lower bound on the outer layer (or SNP-set level)

$$\begin{aligned} \mathrm{LB}(\pi_w, \sigma_w^2, \tau_w^2 \mid \boldsymbol{\theta}) = &- \frac{N}{2}\log(2\pi\tau_w^2) - \frac{1}{2\tau_w^2}\|\mathbf{y} - \mathbf{H}(\boldsymbol{\theta})\boldsymbol{\beta}_w\|_2^2 - \frac{1}{2}\sum_{g=1}^{G}\{\mathbf{H}(\boldsymbol{\theta})^\intercal\mathbf{H}(\boldsymbol{\theta})\}_{gg}\,\mathbb{V}[w_g] \\ &- \sum_{g=1}^{G}\alpha_g \log\left(\frac{\alpha_g}{\pi_w}\right) - \sum_{g=1}^{G}(1-\alpha_g)\log\left(\frac{1-\alpha_g}{1-\pi_w}\right) \\ &+ \frac{1}{2}\sum_{g=1}^{G}\alpha_g\left[1 + \log\left(\frac{s_g^2}{\sigma_w^2\tau_w^2}\right) - \frac{m_g^2 + s_g^2}{\sigma_w^2\tau_w^2}\right] \end{aligned} \tag{17}$$

where, similar to the input layer updates, $\tau_w^2 \approx \mathbb{V}[\mathbf{y} - \mathbf{H}(\boldsymbol{\theta})\boldsymbol{\beta}_w]$ estimates the variance of residual training error in the outer layer; the term $\boldsymbol{\beta}_w$ is a $G$-dimensional estimate of the posterior mean for $\mathbf{w}$ with elements $\beta_{wg} = \alpha_g m_g$ for the $g$-th SNP-set; the matrix $\mathbf{H}(\boldsymbol{\theta})^\intercal\mathbf{H}(\boldsymbol{\theta})$ is deterministically computed given posterior estimates of the weights $\boldsymbol{\theta}$ from the input layer, and $\mathbb{V}[w_g] = \alpha_g(m_g^2 + s_g^2) + \alpha_g^2 m_g^2$ is the variance of the $g$-th weight under the approximating distributional family in Eq. (16). We describe the explicit expectation and maximization steps of the approximate EM algorithm for the outer layer below.

1. **E-Step: Update the Variational Free Parameters.** In the E-step of the algorithm, we this time take the partial derivatives of the lower bound in Eq. (17) with respect to the free parameters in $\boldsymbol{\psi}$ and set them equal to zero. Solving for $m_g$, $s_g^2$, and $\alpha_g$ yields the following updates

$$m_g = \frac{s_g^2}{\tau_w^2}\left[(\mathbf{H}(\boldsymbol{\theta})^\intercal\mathbf{y})_g - \sum_{l \neq g}\{\mathbf{H}(\boldsymbol{\theta})^\intercal\mathbf{H}(\boldsymbol{\theta})\}_{lg}\,\beta_{wl}\right] \tag{18}$$

$$s_g^2 = \tau_w^2\left[\{\mathbf{H}(\boldsymbol{\theta})^\intercal\mathbf{H}(\boldsymbol{\theta})\}_{gg} + \frac{1}{\sigma_w^2}\right]^{-1} \tag{19}$$

$$\alpha_g = \mathrm{Sigmoid}\left(\log\left(\frac{\pi_w}{1-\pi_w}\right) + \log\left(\frac{s_g}{\sigma_w\tau_w}\right) + \frac{m_g^2}{2s_g^2}\right) \tag{20}$$

where $\alpha_g \approx \Pr[\gamma_{wg} = 1 \,|\, \mathbf{y}, \mathbf{X}, \boldsymbol{\theta}, \pi_w, \sigma_w^2, \tau_w^2]$ and, again, we set the sigmoid function to be the standard logistic function.

2. **M-Step: Update the Variance Hyper-Parameters.** In the M-step of the algorithm, we fix values of the variational free parameters and maximize the lower bound in Eq. (17) with respect to $\sigma_w^2$ and $\tau_w^2$. Once again, this is done by solving for the roots $\sigma_w^2$ and $\tau_w^2$ of the gradient which then yields the following updates

$$\sigma_w^2 = \left[\sum_{g=1}^{G} \alpha_g (m_g^2 + s_g^2)\right] \Bigg/ \left(\tau_w^2 \sum_{g=1}^{G} \alpha_g\right) \tag{21}$$

$$\tau_w^2 = \left(N + \sum_{g=1}^{G} \alpha_g\right)^{-1} \left[\|\mathbf{y} - \mathbf{H}(\boldsymbol{\theta})\boldsymbol{\beta}_w\|_2^2 + \sum_{g=1}^{G} \{\mathbf{H}(\boldsymbol{\theta})^{\intercal}\mathbf{H}(\boldsymbol{\theta})\}_{gg}\, \mathbb{V}[w_g] + \frac{1}{\sigma_w^2} \sum_{g=1}^{G} \alpha_g (m_g^2 + s_g^2)\right] \tag{22}$$

where, again, $N$ is equal to the dimensionality of the phenotypic response vector $\mathbf{y}$.

Similar to the algorithmic updates in the input layer, we account for our lack of *a priori* knowledge about the "correct" proportion of enriched SNP-sets by placing another exchangeable uniform hyper-prior distribution over an $L$-valued grid of possible values where $\{\pi_w^{(1)}, \ldots, \pi_w^{(L)}\} \in [1/G, 1]$. Here, we now use the variational lower bound in Eq. (17) to approximate the posterior distribution of $\pi_w$. As a final step in the model fitting procedure, we again conduct a Bayesian model averaging-like procedure by integrating over the different grid combinations for $\pi_w$ and computing marginal posterior inclusion probabilities for each of the $G$-annotated SNP-sets as the following

$$\text{PIP}(g) \equiv \Pr[\gamma_{wg} = 1 \,|\, \mathbf{y}, \mathbf{X}, \boldsymbol{\theta}] \approx \sum_{l=1}^{L} \lambda_w^{(l)} \Pr[\gamma_{wg} = 1 \,|\, \mathbf{y}, \mathbf{X}, \boldsymbol{\theta}, \pi_w^{(l)}, \sigma_w^2, \tau_w^2]. \tag{23}$$

where each importance weight $\lambda_w^{(l)}$ takes on a form similar to the normalized ratio described in Eq. (13).

### 3 Accounting for Non-Additive Genetic Effects

As mentioned in the main text, the BANNs framework models the proportion of phenotypic variance that is explained by sparse genetic effects (both additive and non-additive) [15]. This primarily done through the inclusion of the nonlinear Leaky ReLU activation function $h(\bullet)$ in the hidden layer [1]. In other areas of statistical genetics, similar nonlinear functions have been used to model non-additive effects that contribute to phenotypic variation [29–35]. For example, it has been shown that the Taylor series expansion of the Gaussian kernel function enumerates higher-order interaction terms between SNPs [36–39], thus alleviating potential combinatorial concerns with exhaustive searches [40]. The nonlinear ReLU function family shares this same property [41–43]. To see this, consider the general ReLU function for SNPs in the $g$-th SNP-set which is defined as

$$h(\mathbf{X}_g\boldsymbol{\theta}_g) = \max\left\{0, \sum_{j=1}^{|\mathcal{S}_g|} \mathbf{x}_j\theta_j + \mathbf{1}b_g^{(1)}\right\}. \tag{24}$$

Under this formulation, it becomes clear that the effect that any one SNP has in determining the output of the ReLU activation jointly depends on the effects of all other variants in the SNP-set — therefore, capturing the dependence or interaction between the inputs in the function. The SNP-set specific bias terms $b_g^{(1)}$ allow each node in the hidden layer to change slope for different combinations of genotypes and provides more flexible estimation of broad-sense heritability. Theoretically, as more nodes and hidden layers are added to the network architecture, the model will have an even greater ability to account for non-additive genetic effects (acting similarly to classic Gaussian process regression methods) [44]. Through our simulation studies, we demonstrate the capability to accurately prioritize/rank associated SNPs and enriched SNP-sets in the BANNs framework, even in the presence of pairwise SNP-by-SNP interactions and population structure.

### 4 Estimating Phenotypic Variance Explained (PVE)

As described in the main text, we are able to provide an estimate of phenotypic variance explained (PVE) within the BANNs framework as the total proportion of phenotypic variance that is explained by sparse genetic effects (both additive and non-additive) [15]. Given the true values of the neural network parameters, we define this proportion on the SNP-level in the inner layer and SNP-set level in the outer layer as the following

$$\mathrm{PVE}(\boldsymbol{\theta}) \approx \frac{\mathbb{V}[\mathbf{X}\boldsymbol{\theta}]}{\mathbb{V}[\mathbf{y}]}, \qquad \mathrm{PVE}(\mathbf{w}) \approx \frac{\mathbb{V}[\mathbf{H}(\boldsymbol{\theta})\mathbf{w}]}{\mathbb{V}[\mathbf{y}]}, \tag{25}$$

where, as a reminder, $\mathbb{V}[\bullet]$ is the variance function and $\mathbf{H}(\boldsymbol{\theta}) = [h(\mathbf{X}_1\boldsymbol{\theta}_1 + \mathbf{1}b_1^{(1)}), \dots, h(\mathbf{X}_G\boldsymbol{\theta}_G + \mathbf{1}b_G^{(1)})]$ denotes the matrix of deterministic nonlinear neurons in the hidden layer given estimates of the input layer weights. In practice, we estimate PVE using posterior values of the network parameters derived from the variational EM algorithm described in the previous section. Specifically, after averaging over the grid of different models, we use the (approximate) marginal posterior means $\boldsymbol{\beta}_\theta$ and $\boldsymbol{\beta}_w$ for the input and outer layer weights from Eqs. (8) and (16), respectively. We also approximate the variance of residual error that is observed during the training phase of both layers with estimates of $\tau_\theta^2$ and $\tau_w^2$ from Eqs. (13) and (22). This yields the following empirical estimate for the PVE of complex traits

$$\mathrm{PVE}(\boldsymbol{\theta}) \approx \frac{\mathbb{V}[\mathbf{X}\boldsymbol{\beta}_\theta]}{\mathbb{V}[\mathbf{X}\boldsymbol{\beta}_\theta] + \tau_\theta^2}, \qquad \mathrm{PVE}(\mathbf{w}) \approx \frac{\mathbb{V}[\mathbf{H}(\boldsymbol{\beta}_\theta)\boldsymbol{\beta}_w]}{\mathbb{V}[\mathbf{H}(\boldsymbol{\beta}_\theta)\boldsymbol{\beta}_w] + \tau_w^2}, \tag{26}$$

where the matrix hidden neurons is empirically estimated as $\mathbf{H}(\boldsymbol{\beta}_\theta) = [h(\mathbf{X}_1\boldsymbol{\beta}_{\theta 1} + b_1^{(1)}), \dots, h(\mathbf{X}_G\boldsymbol{\beta}_{\theta G} + b_G^{(1)})]$. Note that this formula is similar to the traditional form used for estimating PVE, except here we also consider the contribution of non-additive genetic effects through the nonlinear Leaky ReLU activation function $h(\bullet)$ [1]. Through various simulations, we demonstrate the ability to accurately estimate PVE in the BANNs framework under additive sparse architectures (see Figs. S26 and S27). We underestimate PVE in both polygenic traits and traits with pairwise SNP-by-SNP interactions, which we believe is caused by either ($i$) a misestimation of the approximate posterior mean for network weights or ($ii$) an over-estimation of the residual error variance during the variational EM algorithm. Similar observations have been noted when using variational inference [9,19,25,45]. Results from other work also suggest that the sparsity assumption on the SNP-level effects can lead to the underestimation of the PVE [14,15].

### 5 Ablation Test and Analysis

To investigate how choices in the model setup contribute to variable selection, we performed an "ablation analysis" where we modified parts of the BANNs framework independently and observed their direct effect on model performance. We considered two different modifications to our model: (1) removing the activation function and training a fully linear hierarchical model, and (2) removing the approximate Bayesian model averaging approach and updating the probabilities $\pi_\theta$ and $\pi_w$ as additional parameters in the variational EM algorithm. In the normal BANNs setup, we initialize $L$ different models with varying priors for inclusion probabilities specified over a grid $\{\pi_\theta^{(1)}, \ldots, \pi_\theta^{(L)}\} \in [1/J, 1]$ and $\{\pi_w^{(1)}, \ldots, \pi_w^{(L)}\} \in [1/G, 1]$, respectively. However, in the case of the latter ablation modification, we initialize $\pi_\theta = 1/J$ and $\pi_w = 1/G$ as an analogy to the "single causal variant" assumption frequently used in fine mapping [64]. Next, we update their values in the M-step of the algorithm according to the following analytic expressions

$$\frac{\pi_\theta}{1-\pi_\theta} = \frac{\sum_j \sum_k \alpha_{jk}}{\sum_j \sum_k (1-\alpha_{jk})}, \qquad \frac{\pi_w}{1-\pi_w} = \frac{\sum_g \alpha_g}{\sum_g (1-\alpha_g)}. \tag{27}$$

Results presented in the main text and Supporting Information (see Fig. S25) are shown using simulations with the self-identified "white British" ancestry cohort from the UK Biobank on synthetic traits that have broad-sense heritability $H^2 = 0.6$. Each plot combines results from 100 simulated replicates (see Section 9 for details on the setup for the simulation study).

### 6 Data Quality Control Procedures for Stock of Mice

Some of the real data analysis results in this work made use of GWA data from the Wellcome Trust Centre for Human Genetics (`http://mtweb.cs.ucl.ac.uk/mus/www/mouse/index.shtml`). This study contains $N$ = 1,814 heterogenous stock of mice from 85 families (all descending from eight inbred progenitor strains) [46], and 131 quantitative traits that are classified into 6 broad categories including behavior, diabetes, asthma, immunology, haematology, and biochemistry (`http://mtweb.cs.ucl.ac.uk/mus/www/GSCAN/index.shtml/index.old.shtml`). In the main text, we focused on six specific phenotypes from these categories including: body mass index (BMI) (`Obesity.BMI`), body weight (`Glucose.BodyWeight`), percentage of CD8+ cells (`Imm.PctCD8`), mean corpuscular hemoglobin (MCH) (`Haem.MCH`), high-density lipoprotein content (`Biochem.HDL`), and low-density lipoprotein content (`Biochem.LDL`). All phenotypes were previously corrected for sex, age, body weight, season, year, and cage effects [46]. For individuals with missing genotypes, we imputed values by the mean genotype of that SNP in their corresponding family. Only polymorphic SNPs with minor allele frequency above 5% were kept for the analyses. This left a total of $J$ = 10,227 autosomal SNPs that were available for all mice. For annotations, we used the Mouse Genome Informatics database (`http://www.informatics.jax.org`) to map SNPs to the closest neighboring gene(s). Here, pseudogenes, quantitative trait loci (QTL), and genes with only 1 annotated SNP within their boundary were excluded from the analyses. Unannotated SNPs located within the same genomic region were labeled as being within the "intergenic region" between two genes. Altogether, a total of $G$ = 1,925 SNP-sets were analyzed.

### 7 Data Quality Control Procedures for Framingham Heart Study

The other real data analysis results made use of human GWA data from the Framingham Heart Study (`https://www.ncbi.nlm.nih.gov/gap`) [47]. This study originally contains $N$ = 6,950 individuals and $J$ = 394,174 SNPs. For quality control on these data, we removed ($i$) SNPs with minor allele frequency less than 2.5%, ($ii$) SNPs not in Hardy-Weinberg Equilibrium (Fisher's exact test $P > 1 \times 10^{-4}$), and ($iii$) proximal SNPs in high linkage disequilibrium (using the flag `--indep-pairwise 50 5 0.8` with PLINK 1.9 [48]). This resulted in a final dataset containing $J$ = 372,131 SNPs, where any missing values for a given SNP were imputed by using the estimated mean genotype of that SNP. Next, we used the NCBI's Reference Sequence (RefSeq) database in the UCSC Genome Browser [49] to annotate SNPs with appropriate genes. Recall that in the real data analysis, we define genes with boundaries in two ways: ($a$) we use the UCSC gene boundary definitions directly, or ($b$) we augment the gene boundaries by adding SNPs within a ±500 kilobase (kb) buffer to account for possible regulatory elements. Genes with only 1 SNP within their boundary were excluded from either analysis. Unannotated SNPs located within the same genomic region were labeled as being within the "intergenic region" between two genes. Altogether, a total of $G = 18{,}364$ SNP-sets were analyzed—which included 8,658 intergenic SNP-sets and 9,706 annotated genes—using the UCSC boundaries. When including the 500kb buffer, a total of $G = 35{,}871$ SNP-sets were analyzed.

### 8 Data Quality Control Procedures for UK Biobank

The simulation results and additional lipoprotein study presented in the main text made use of imputed data released from the UK Biobank [50]. Quality control procedures for these data are as follows. First, we only studied individuals who self-identified as being of European ancestry. From this cohort, we further excluded individuals identified by the UK Biobank to have high heterozygosity, excessive relatedness, or aneuploidy (1,550 individuals removed). We also removed individuals whose kinship coefficient was greater than 0.0442 (i.e., close relatives). Next, we removed (*i*) monomorphic SNPs, (*ii*) SNPs with minor allele frequency less than 2.5%, (*iii*) SNPs not in Hardy-Weinberg Equilibrium (Fisher's exact test $P > 1 \times 10^{-6}$), (*iv*) SNPs with missingness greater than 1%, and (*v*) SNPs in high linkage disequilibrium (using the flag `--indep-pairwise 50 5 0.9` with PLINK 1.9 [48]). After all QC steps, we had a final dataset of $N = 349{,}414$ individuals and $J = 1{,}070{,}306$ SNPs. Next, we used the NCBI's Reference Sequence (RefSeq) database in the UCSC Genome Browser [49] to annotate SNPs with appropriate genes. Again, in the real data analysis, we define genes with boundaries in two ways: (*a*) we use the UCSC gene boundary definitions directly, or (*b*) we augment the gene boundaries by adding SNPs within a $\pm 500$ kilobase (kb) buffer to account for possible regulatory elements. Genes with only 1 SNP within their boundary were excluded from either analysis. Unannotated SNPs located within the same genomic region were labeled as being within the "intergenic region" between two genes. Altogether, a total of $G = 28{,}644$ SNP-sets were kept for analysis using the UCSC boundaries and a total of $G = 35{,}849$ SNP-sets were kept for analysis when including the 500kb buffer.

### 9 Simulation Setup and Scenarios

In our simulation studies, we used the following general simulation scheme to generate quantitative traits using real genotype data on chromosome 1 from ten thousand randomly sampled individuals of European ancestry in the UK Biobank [50]. We consider two different data compositions. In the first, we simulate synthetic traits only using individuals who self-identify as being of "white British" ancestry. In the second, we simulate phenotypes by random subsampling 3,000 individuals who self-identify as being of "white British" ancestry, 3,000 individuals who self-identify as being of "white Irish" ancestry, and 4,000 individuals who identify as being of "any other white background". Note that the latter composition introduces additional, yet cryptic, population structure into the problem. In the main text and Supporting information (i.e., Figs. 2-3, Figs. S2-S27, and Tables S1-S8), we refer to these datasets as the "British" and "European" cohorts, respectively.

The setup to generate synthetic traits follows mostly from previous studies [13,38–40]. We will denote this genotype matrix as $\mathbf{X}$, with $\mathbf{x}_j$ denoting the genotypic vector for the $j$-th SNP. Following quality control procedures detailed in the previous section, our simulations included $J = 36{,}518$ SNPs distributed across genome. Again, we used the NCBI's RefSeq database in the UCSC Genome Browser to assign SNPs to genes which resulted in 1,408 genes to be used in the simulation study. We also consider the unnanotated SNPs between two genes to be located within intergenic regions. Altogether, a total of $G = 2{,}816$ SNP-sets were analyzed.

After the annotation step, we assume that all simulated traits have been standardized such that $\mathbb{V}[\mathbf{y}] = 1$ and that all observed genetic effects explain a fixed proportion of this value (i.e., broad-sense heritability, $H^2$). To be explicit, one can equate the total PVE in these simulations to $H^2$. Next, we use the $N \times J$ matrix of genotypes $\mathbf{X}$ to generate real-valued phenotypes that mirror genetic architectures affected by a combination of linear (additive) and interaction (epistatic) effects. We randomly select a certain percentage of truly associated SNP-sets and denote the SNPs that they contain as $\mathcal{C}$. Within $\mathcal{C}$, we select causal SNPs in a way such that each associated SNP-set contains at least two SNPs with non-zero effects. The additive effect size for all causal SNPs are assumed to come from a standard normal distribution, $\boldsymbol{\theta} \sim \mathcal{N}(\mathbf{0}, \mathbf{I})$. Next, we create a separate matrix $\mathbf{W}$ which holds the pairwise interactions between the causal SNPs in enriched SNP-sets. This is done by taking the Hadamard (element-wise) product between genotypic vectors of SNPs within $\mathcal{C}$. The corresponding interaction effect sizes are drawn as $\boldsymbol{\varphi} \sim \mathcal{N}(\mathbf{0}, \mathbf{I})$. We scale both the additive and pairwise genetic effects so that collectively they explain a fixed proportion of genetic variance. Namely, the additive effects make up $\rho\%$ while the pairwise interactions make up the remaining $(1-\rho)\%$. Alternatively, the proportion of the heritability explained by additivity is said to be $\mathbb{V}[\sum \mathbf{x}_c \theta_c] = \rho H^2$, while the proportion detailed by genetic interactions is given as $\mathbb{V}[\mathbf{W}\boldsymbol{\varphi}] = (1-\rho)H^2$. We consider two choices for the parameter $\rho = \{0.5, 1\}$. Intuitively, $\rho = 1$ represents the limiting case where the variation of a trait is driven by solely additive effects. For $\rho = 0.5$, the additive and pairwise interaction effects are assumed to equally contribute to the phenotypic variance. Once we obtain the final effect sizes for all causal variants, we draw normally distributed random errors as $\boldsymbol{\varepsilon} \sim \mathcal{N}(\mathbf{0}, \mathbf{I})$ to make up the remaining percentage of the total variance. Quantitative continuous traits are then generated under the following general linear model:

$$\mathbf{y} = \sum_{c \in \mathcal{C}} \mathbf{x}_c \theta_c + \mathbf{W}\boldsymbol{\varphi} + \boldsymbol{\varepsilon}. \tag{28}$$

Given the simulation procedure above, we randomly sample $N = 10{,}000$ individuals and simulate a wide range of scenarios for comparing the performance of both SNP and SNP-set level association methods. Here, we vary the following simulation parameters:

- Broad-sense heritability: $H^2 = 0.2$ and 0.6;
- Contribution of interaction effects: $(1-\rho) = 0$ and 0.5;
- Percentage of associated SNP-sets: 1% (sparse architecture) and 10% (polygenic architecture);

Lastly, we set the number of causal SNPs with non-zero effects to be some fixed percentage of all SNPs located within the selected associated SNP-sets. We set this percentage to be 1% in the 1% associated SNP-set case, and 10% in the 10% associated SNP-set case. All performance comparisons are based on 100 different simulated runs for each parameter combination. For evaluating the performance of each method, we assessed the following:

- The power and false discovery rates when identifying causal SNPs or associated SNP-sets at a Bonferroni-corrected threshold for frequentist approaches ($P = 0.05/36518 = 1.37 \times 10^{-6}$ at the SNP-level and $P = 0.05/2816 = 1.78 \times 10^{-5}$ at the SNP-set level) or according to the median probability model for Bayesian methods (posterior enrichment probability $> 0.5$) [51];
- The ability to rank true positive (TP) genes over false positives (FP) via receiver operating characteristic (ROC) and precision-recall curves.

All figures and tables show the mean performances (and standard errors) across all simulated replicates.

### 10 Software Details

Source code for the BANNs framework is freely available at `https://github.com/lcrawlab/BANNs` and is licensed under the GNU General Public License (version 3.0). We have released two versions of the

BANNs software: one implemented within `Python 3` (release version 3.7.7) and other within `R` (compatible with versions 3.3.2 through 3.6.3). The BANNs GitHub repository includes example data, documentation, and instructions for how to execute the code within both coding languages. Results in the main text and Supporting Information are based on the `Python 3` implementation which depends on the `pandas` library (version 1.0.1) [52] for automatically creating partial neural network architectures based on the biological annotations provided by the user; the `NumPy` (version 1.18-19) [53] and `Numba` (version 0.48.0) [54] packages for efficient matrix operations; and the `multiprocessing` library (version 2.6) [55] for parallelizing posterior computation over multiple threads and providing faster execution. Training, estimation of the network parameters, and optimization was done by using an Adam optimizer [56] in `TensorFlow` (version 1.5). While the software can be run directly using the source code, it can also be installed as a package through `pip` with the command: `pip3 install BANNs`. All dependencies are also automatically installed with the package.

The `R` implementation uses the `dplyr` package (version 0.8.5) [57] for automatically creating partial neural network architectures based on the biological annotations provided by the user; the `Matrix` package (version 1.2-18) [58] for efficient matrix operations; and the `doParallel` (version 1.0.15) [59], `forEach` (version 1.4.8) [60], `iterators` (version 1.0.12) [61], and standard `parallel` packages for parallelized execution of the variational expectation-maximization algorithm. Similarly, the `R` implementation of the software can be run by directly downloading the source code or it can be installed using `devtools` [62] with the commands: `devtools::install("lcrawlab/BANNs")` and `library(BANNs)`.

**Software Details for Competing Approaches.** In this work, comparisons to SNP-level association mapping methods were made using software for CAVIAR (version 2.0.0; `http://genetics.cs.ucla.edu/caviar/`), FINEMAP (version 1.4; `http://www.christianbenner.com`), and SuSiE (version 0.9.0; `https://github.com/stephenslab/susieR`). Comparisons to SNP-set mapping methods were made using software for GBJ (version 0.5.3; `https://cran.r-project.org/web/packages/GBJ/`), GSEA (`https://www.nr.no/en/projects/software-genomics`), MAGMA (version 1.07b; `https://ctg.cncr.nl/software/magma`), PEGASUS (version 1.3.0; `https://github.com/ramachandran-lab/PEGASUS`), RSS (version 1.0.0; `https://github.com/stephenslab/rss`), and SKAT (version 1.3.2.1; `https://www.hsph.harvard.edu/skat`), which are also publicly available. All software for competing methods were fit using the default settings, unless otherwise stated in the main text and Supporting Information.

### 11 Pseudocode for Biologically Annotated Neural Networks

**Algorithm 1** BANNs Model with Individual Level Data

Input genotype data $\mathbf{X}$, continuous trait $\mathbf{y}$, and annotations $\{\mathcal{S}_1, \ldots, \mathcal{S}_G\}$.<br>
Choose the number of models $L$, number of maximum iterations $T$, and tolerance parameter $\epsilon$.<br>
Set up the $L$-grid of possible values $\{\pi_\theta^{(1)}, \ldots, \pi_\theta^{(L)}\} \in [1/J, 1]$ and $\{\pi_w^{(1)}, \ldots, \pi_w^{(L)}\} \in [1/G, 1]$ for the inner and outer layer, respectively.<br>
Randomly initialize variational parameters $\{\alpha_{jk}, m_{jk}, s_{jk}^2\}_{k=1}^K$, $\{\sigma_{\theta k}^2\}_{k=1}^K$, and $\tau_\theta^2$ for the inner layer.<br>
Randomly initialize variational parameters $\{\alpha_g, m_g, s_g^2\}$, $\sigma_w^2$, and $\tau_w^2$ for the outer layer.<br>
**for each** $\pi_\theta^{(l)} \in \{\pi_\theta^{(1)}, \ldots, \pi_\theta^{(L)}\}$ and $\pi_w^{(l)} \in \{\pi_w^{(1)}, \ldots, \pi_w^{(L)}\}$ **do**<br>
    Compute inner lower bound `LB_inner_new`.<br>
    **for** $t = 1 \to T$ **do** ▷ Inner Layer Updates<br>
        Set `LB_inner` = `LB_inner_new`.<br>
        Update variational parameters $\{\alpha_{jk}, m_{jk}, s_{jk}^2\}_{k=1}^K$ for $j = 1, \ldots, J$ SNPs. ▷ E-Step<br>
        Update hyper-parameters parameters $\{\sigma_{\theta k}^2\}_{k=1}^K$ and $\tau_\theta^2$. ▷ M-Step<br>
        Update lower bound `LB_inner_new`.<br>
        **if** `LB_inner_new` $-$ `LB_inner` $\leq \epsilon$ **then**<br>
            Save `LB_inner` = `LB_inner_new`.<br>
            Break<br>
        **end if**<br>
    **end for**<br>
    Compute hidden layer neurons $\mathbf{H}(\boldsymbol{\theta})$.<br>
    Compute outer lower bound `LB_outer_new`.<br>
    **for** $t = 1 \to T$ **do** ▷ Outer Layer Updates<br>
        Set `LB_outer` = `LB_outer_new`.<br>
        Update variational parameters $\{\alpha_g, m_g, s_g^2\}$ for $g = 1, \ldots, G$ SNP-sets. ▷ E-Step<br>
        Update hyper-parameters parameters $\sigma_w^2$ and $\tau_w^2$. ▷ M-Step<br>
        Update lower bound `LB_outer_new`.<br>
        **if** `LB_outer_new` $-$ `LB_outer` $\leq \epsilon$ **then**<br>
            Save `LB_outer` = `LB_inner_outer`.<br>
            Break<br>
        **end if**<br>
    **end for**<br>
**end for each**<br>
Compute normalized importance weights $\lambda_\theta^{(l)}$ and $\lambda_g^{(l)}$ for $l = 1, \ldots, L$ models.<br>
Compute (marginal) posterior means $\boldsymbol{\beta}_\theta$ and $\boldsymbol{\beta}_w$ for network weights $\boldsymbol{\theta}$ and $\mathbf{w}$, respectively.<br>
Compute (marginal) posterior inclusion probabilities $\text{PIP}(\boldsymbol{\theta})$ and $\text{PIP}(\mathbf{w})$.<br>
Compute the phenotypic variance explained by the input and hidden layers $\text{PVE}(\boldsymbol{\theta})$ and $\text{PVE}(\mathbf{w})$.<br>
**Return** $\{\boldsymbol{\beta}_\theta, \boldsymbol{\beta}_w, \text{PIP}(\boldsymbol{\theta}), \text{PIP}(\mathbf{w}), \text{PVE}(\boldsymbol{\theta}), \text{PVE}(\mathbf{w})\}$.

**Algorithm 2** BANN-SS Model with GWA Summary Statistics

Input LD matrix $\mathbf{R}$, OLS effect size estimates $\widehat{\boldsymbol{\theta}}$, and annotations $\{\mathcal{S}_1, \ldots, \mathcal{S}_G\}$.
Choose the number of models $L$, number of maximum iterations $T$, and tolerance parameter $\epsilon$.
Set up the $L$-grid of possible values $\{\pi_\theta^{(1)}, \ldots, \pi_\theta^{(L)}\} \in [1/J, 1]$ and $\{\pi_w^{(1)}, \ldots, \pi_w^{(L)}\} \in [1/G, 1]$ for the inner and outer layer, respectively.
Randomly initialize variational parameters $\{\alpha_{jk}, m_{jk}, s_{jk}^2\}_{k=1}^K$, $\{\sigma_{\theta k}^2\}_{k=1}^K$, and $\tau_\theta^2$ for the inner layer.
Randomly initialize variational parameters $\{\alpha_g, m_g, s_g^2\}$, $\sigma_w^2$, and $\tau_w^2$ for the outer layer.
**for each** $\pi_\theta^{(l)} \in \{\pi_\theta^{(1)}, \ldots, \pi_\theta^{(L)}\}$ and $\pi_w^{(l)} \in \{\pi_w^{(1)}, \ldots, \pi_w^{(L)}\}$ **do**
    Compute inner lower bound `LB_inner_new`.
  **for** $t = 1 \to T$ **do** ▷ Inner Layer Updates
      Set `LB_inner` = `LB_inner_new`.
      Update variational parameters $\{\alpha_{jk}, m_{jk}, s_{jk}^2\}_{k=1}^K$ for $j = 1, \ldots, J$ SNPs. ▷ E-Step
      Update hyper-parameters parameters $\{\sigma_{\theta k}^2\}_{k=1}^K$ and $\tau_\theta^2$. ▷ M-Step
      Update lower bound `LB_inner_new`.
    **if** `LB_inner_new` − `LB_inner` $\leq \epsilon$ **then**
        Save `LB_inner` = `LB_inner_new`.
        Break
    **end if**
  **end for**
    Compute hidden layer neurons $\mathbf{H}(\boldsymbol{\theta})$.
    Compute outer lower bound `LB_outer_new`.
  **for** $t = 1 \to T$ **do** ▷ Outer Layer Updates
      Set `LB_outer` = `LB_outer_new`.
      Update variational parameters $\{\alpha_g, m_g, s_g^2\}$ for $g = 1, \ldots, G$ SNP-sets. ▷ E-Step
      Update hyper-parameters parameters $\sigma_w^2$ and $\tau_w^2$. ▷ M-Step
      Update lower bound `LB_outer_new`.
    **if** `LB_outer_new` − `LB_outer` $\leq \epsilon$ **then**
        Save `LB_outer` = `LB_inner_outer`.
        Break
    **end if**
  **end for**
**end for each**
Compute normalized importance weights $\lambda_\theta^{(l)}$ and $\lambda_g^{(l)}$ for $l = 1, \ldots, L$ models.
Compute (marginal) posterior means $\boldsymbol{\beta}_\theta$ and $\boldsymbol{\beta}_w$ for network weights $\boldsymbol{\theta}$ and $\mathbf{w}$, respectively.
Compute (marginal) posterior inclusion probabilities $\mathrm{PIP}(\boldsymbol{\theta})$ and $\mathrm{PIP}(\mathbf{w})$.
Compute the phenotypic variance explained by the input and hidden layers $\mathrm{PVE}(\boldsymbol{\theta})$ and $\mathrm{PVE}(\mathbf{w})$.
**Return** $\{\boldsymbol{\beta}_\theta, \boldsymbol{\beta}_w, \mathrm{PIP}(\boldsymbol{\theta}), \mathrm{PIP}(\mathbf{w}), \mathrm{PVE}(\boldsymbol{\theta}), \mathrm{PVE}(\mathbf{w})\}$.

### 12 Supplementary Figures

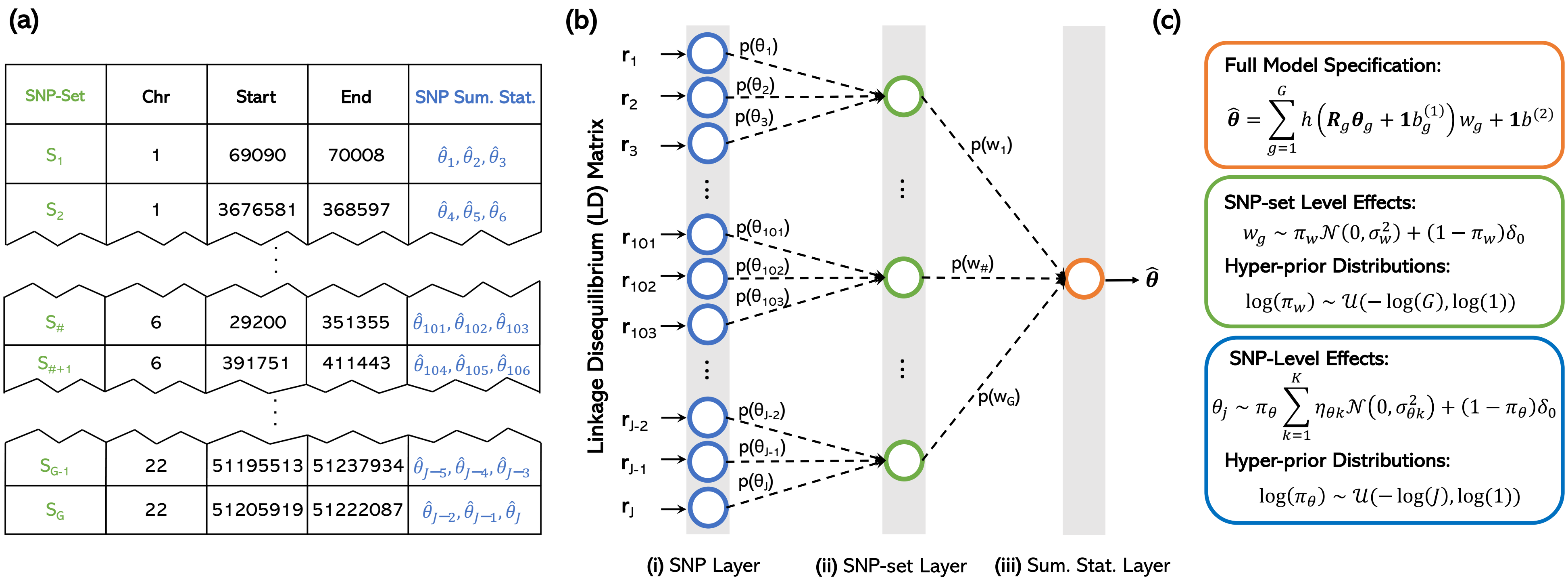

**Figure S1. Biologically annotated neural networks also take in GWA summary statistics (BANN-SS) for multi-scale genotype-phenotype by specifying a partially connected architecture based on the hierarchical nature of enrichment studies. (a)** The BANN-SS framework requires a $J$-dimensional vector of SNP-level GWA marginal effect size (OLS) estimates $\widehat{\boldsymbol{\theta}} = (\widehat{\theta}_1, \ldots, \widehat{\theta}_J)$; an empirical $J \times J$ linkage disequilibrium (LD) matrix $\mathbf{R} = [\mathbf{r}_1, \ldots, \mathbf{r}_J]$, where $\mathbf{r}_j = [r(\mathbf{x}_j, \mathbf{x}_1), \ldots, r(\mathbf{x}_j, \mathbf{x}_J)]$ is a vector of correlation coefficients between the $j$-th SNP and all other SNPs in the study; and a list of $G$-predefined SNP-sets $\{\mathcal{S}_1, \ldots, \mathcal{S}_G\}$. In this work, SNP-sets are defined as genes and intergenic regions (between genes) given by the NCBI's Reference Sequence (RefSeq) database in the UCSC Genome Browser [49]. **(b)** A partially connected Bayesian neural network is constructed based on the annotated SNP groups. In the first hidden layer, only SNPs within the boundary of a gene are connected to the same node. Similarly, SNPs within the same intergenic region between genes are connected to the same node. Completing this specification for all SNPs gives the hidden layer the natural interpretation of being the "SNP-set" layer. **(c)** The hierarchical nature of the network is represented as nonlinear regression model. The corresponding weights in both the SNP ($\boldsymbol{\theta}$) and SNP-set ($\mathbf{w}$) layers are treated as random variables with biologically motivated sparse prior distributions. Posterior inclusion probabilities $\mathrm{PIP}(j) \equiv \Pr[\theta_j \neq 0 \,|\, \mathbf{y}, \mathbf{X}]$ and $\mathrm{PIP}(g) \equiv \Pr[w_g \neq 0 \,|\, \mathbf{y}, \mathbf{X}, \boldsymbol{\theta}_g]$ summarize associations at the SNP and SNP-set level, respectively. The BANN-SS framework uses the same variational inference procedure that is used when we have access to individual-level data.

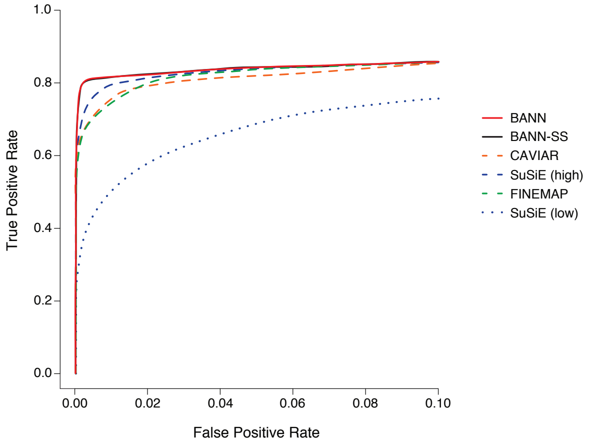

(a) SNP-Level Methods (Sparse Traits)

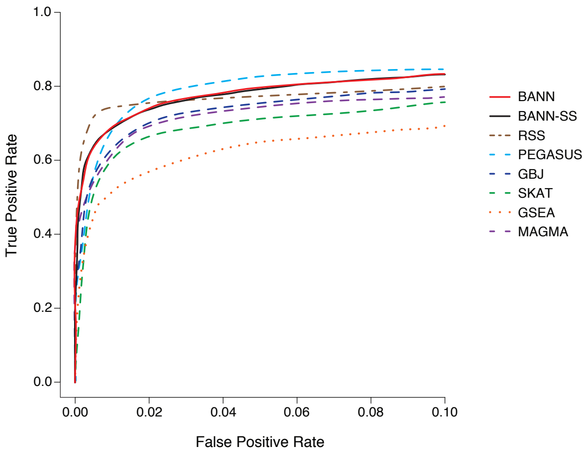

(b) SNP-Set Methods (Sparse Traits)

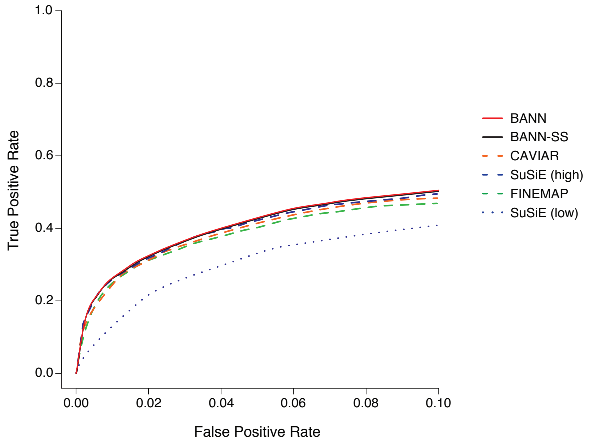

(c) SNP-Level Methods (Polygenic Traits)

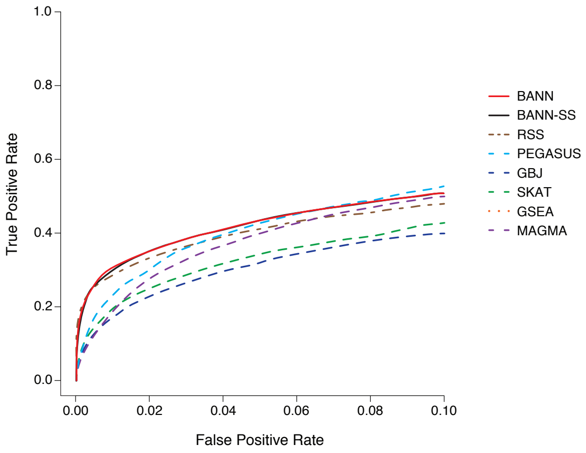

(d) SNP-Set Methods (Polygenic Traits)

**Figure S2. Receiver operating characteristic (ROC) curves comparing the performance of the BANNs (red) and BANN-SS (black) models with competing SNP and SNP-set mapping approaches in simulations (British cohort).** Here, quantitative traits are simulated to have broad-sense heritability of $H^2 = 0.2$ with only contributions from additive effects (i.e., $\rho = 1$). We show power versus false positive rate for two different trait architectures: **(a, b)** sparse where only 1% of SNP-sets are enriched for the trait; and **(c, d)** polygenic where 10% of SNP-sets are enriched. We then set the number of causal SNPs with non-zero effects to be 1% and 10% of all SNPs located within the selected enriched SNP-sets, respectively. To derive results, the full genotype matrix and phenotypic vector are given to the BANNs model and all competing methods that require individual-level data. For the BANN-SS model and other competing methods that take GWA summary statistics, we compute standard GWA SNP-level effect sizes and $P$-values (estimated using ordinary least squares). **(a, c)** Competing SNP-level mapping approaches include: CAVIAR [63], SuSiE [64], and FINEMAP [65]. The software for SuSiE requires an input $\ell$ which fixes the maximum number of causal SNPs in the model. We display results when this input number is high ($\ell = 3000$) and when this input number is low ($\ell = 10$). **(b, d)** Competing SNP-set mapping approaches include: RSS [7], PEGASUS [66], GBJ [67], SKAT [68], GSEA [69], and MAGMA [70]. Note that the upper limit of the x-axis has been truncated at 0.1. All results are based on 100 replicates (see Supporting Information, Section 9).

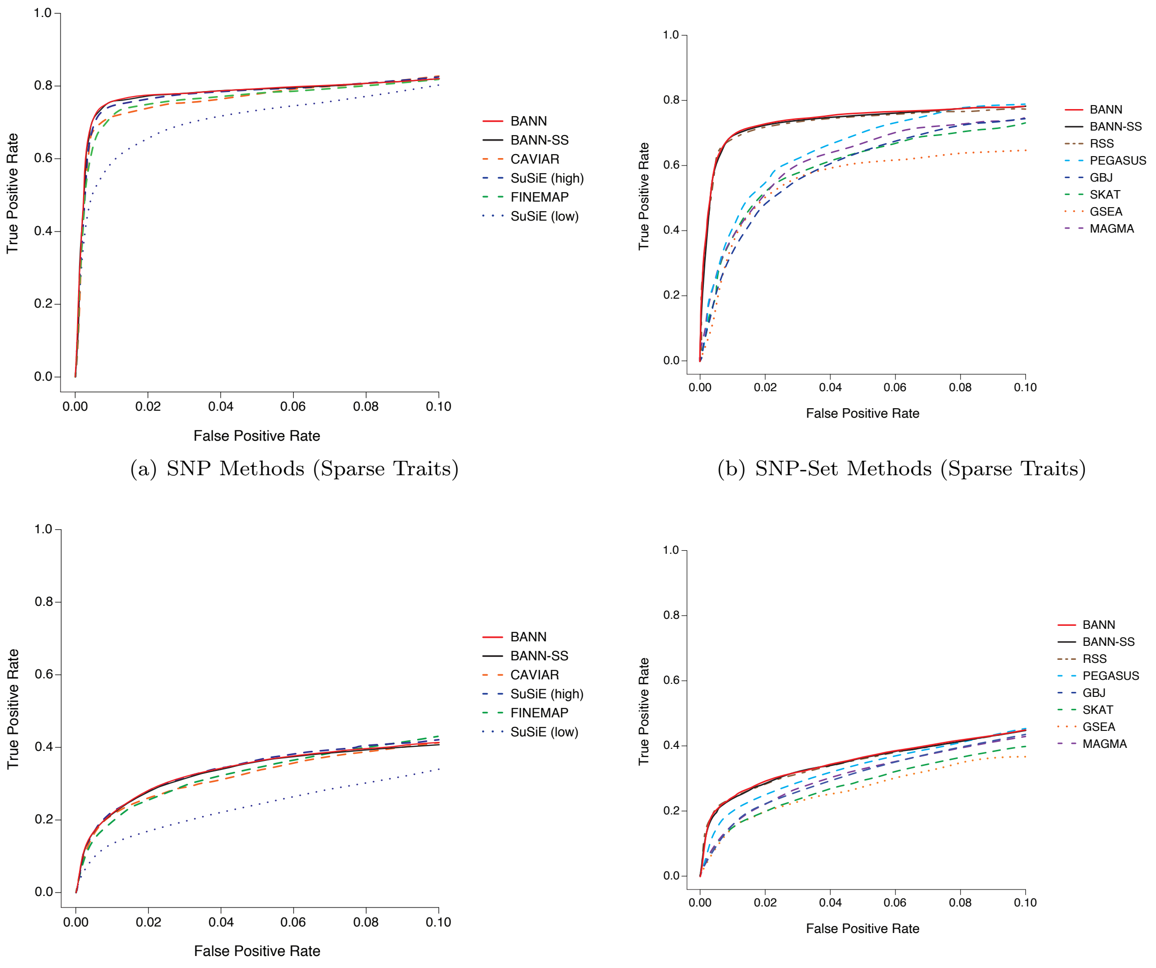

(a) SNP Methods (Sparse Traits)

(b) SNP-Set Methods (Sparse Traits)

(c) SNP Methods (Polygenic Traits)

(d) SNP-Set Methods (Polygenic Traits)

**Figure S3. Receiver operating characteristic (ROC) curves comparing the performance of the BANNs (red) and BANN-SS (black) models with competing SNP and SNP-set mapping approaches in simulations with population structure (European cohort).** Here, quantitative traits are simulated to have broad-sense heritability of $H^2 = 0.2$ with only contributions from additive effects (i.e., $\rho = 1$). In these simulations, traits were generated while using the top ten principal components (PCs) of the genotype matrix as covariates. We show power versus false positive rate for two different trait architectures: **(a, b)** sparse where only 1% of SNP-sets are enriched for the trait; and **(c, d)** polygenic where 10% of SNP-sets are enriched. We then set the number of causal SNPs with non-zero effects to be 1% and 10% of all SNPs located within the selected enriched SNP-sets, respectively. To derive results, the full genotype matrix and phenotypic vector are given to the BANNs model and all competing methods that require individual-level data. For the BANN-SS model and other competing methods that take GWA summary statistics, we compute standard GWA SNP-level effect sizes and $P$-values (estimated using ordinary least squares). **(a, c)** Competing SNP-level mapping approaches include: CAVIAR [63], SuSiE [64], and FINEMAP [65]. The software for SuSiE requires an input $\ell$ which fixes the maximum number of causal SNPs in the model. We display results when this input number is high ($\ell = 3000$) and when this input number is low ($\ell = 10$). **(b, d)** Competing SNP-set mapping approaches include: RSS [7], PEGASUS [66], GBJ [67], SKAT [68], GSEA [69], and MAGMA [70]. Note that the upper limit of the x-axis has been truncated at 0.1. All results are based on 100 replicates (see Supporting Information, Section 9).

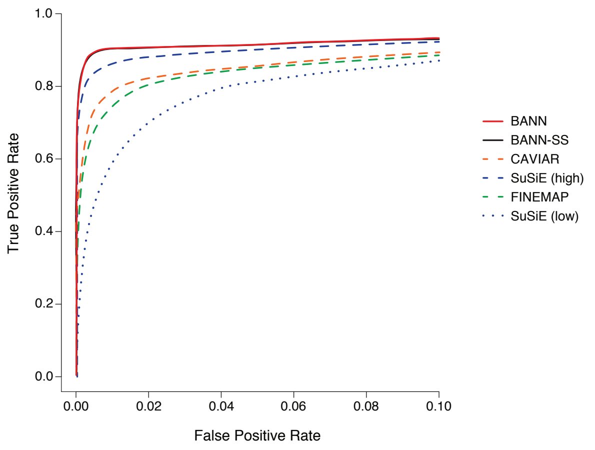

(a) SNP Methods (Sparse Traits)

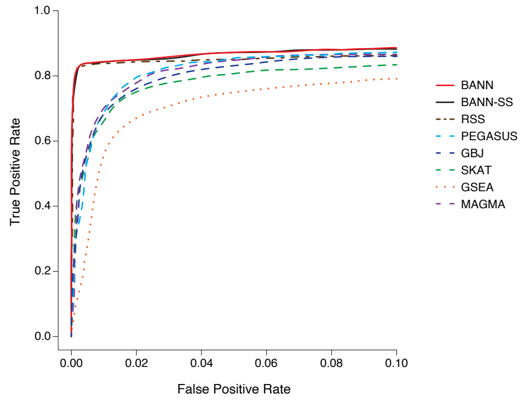

(b) SNP-Set Methods (Sparse Traits)

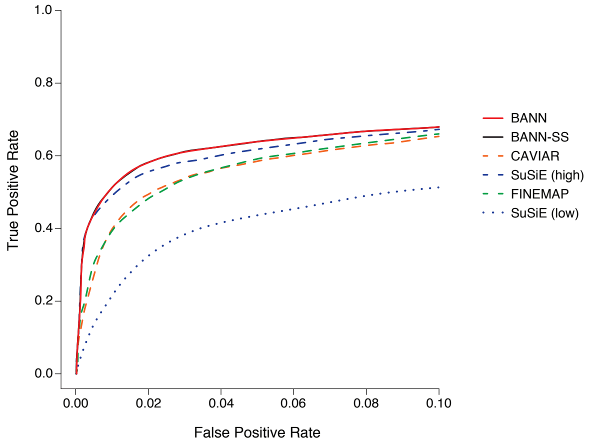

(c) SNP Methods (Polygenic Traits)

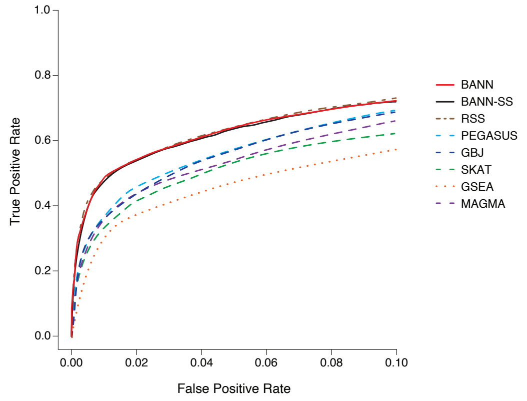

(d) SNP-Set Methods (Polygenic Traits)

**Figure S4. Receiver operating characteristic (ROC) curves comparing the performance of the BANNs (red) and BANN-SS (black) models with competing SNP and SNP-set mapping approaches in simulations with population structure (European cohort).** Here, quantitative traits are simulated to have broad-sense heritability of $H^2 = 0.6$ with only contributions from additive effects (i.e., $\rho = 1$). In these simulations, traits were generated while using the top ten principal components (PCs) of the genotype matrix as covariates. We show power versus false positive rate for two different trait architectures: **(a, b)** sparse where only 1% of SNP-sets are enriched for the trait; and **(c, d)** polygenic where 10% of SNP-sets are enriched. We then set the number of causal SNPs with non-zero effects to be 1% and 10% of all SNPs located within the selected enriched SNP-sets, respectively. To derive results, the full genotype matrix and phenotypic vector are given to the BANNs model and all competing methods that require individual-level data. For the BANN-SS model and other competing methods that take GWA summary statistics, we compute standard GWA SNP-level effect sizes and $P$-values (estimated using ordinary least squares). **(a, c)** Competing SNP-level mapping approaches include: CAVIAR [63], SuSiE [64], and FINEMAP [65]. The software for SuSiE requires an input $\ell$ which fixes the maximum number of causal SNPs in the model. We display results when this input number is high ($\ell = 3000$) and when this input number is low ($\ell = 10$). **(b, d)** Competing SNP-set mapping approaches include: RSS [7], PEGASUS [66], GBJ [67], SKAT [68], GSEA [69], and MAGMA [70]. Note that the upper limit of the x-axis has been truncated at 0.1. All results are based on 100 replicates (see Supporting Information, Section 9).

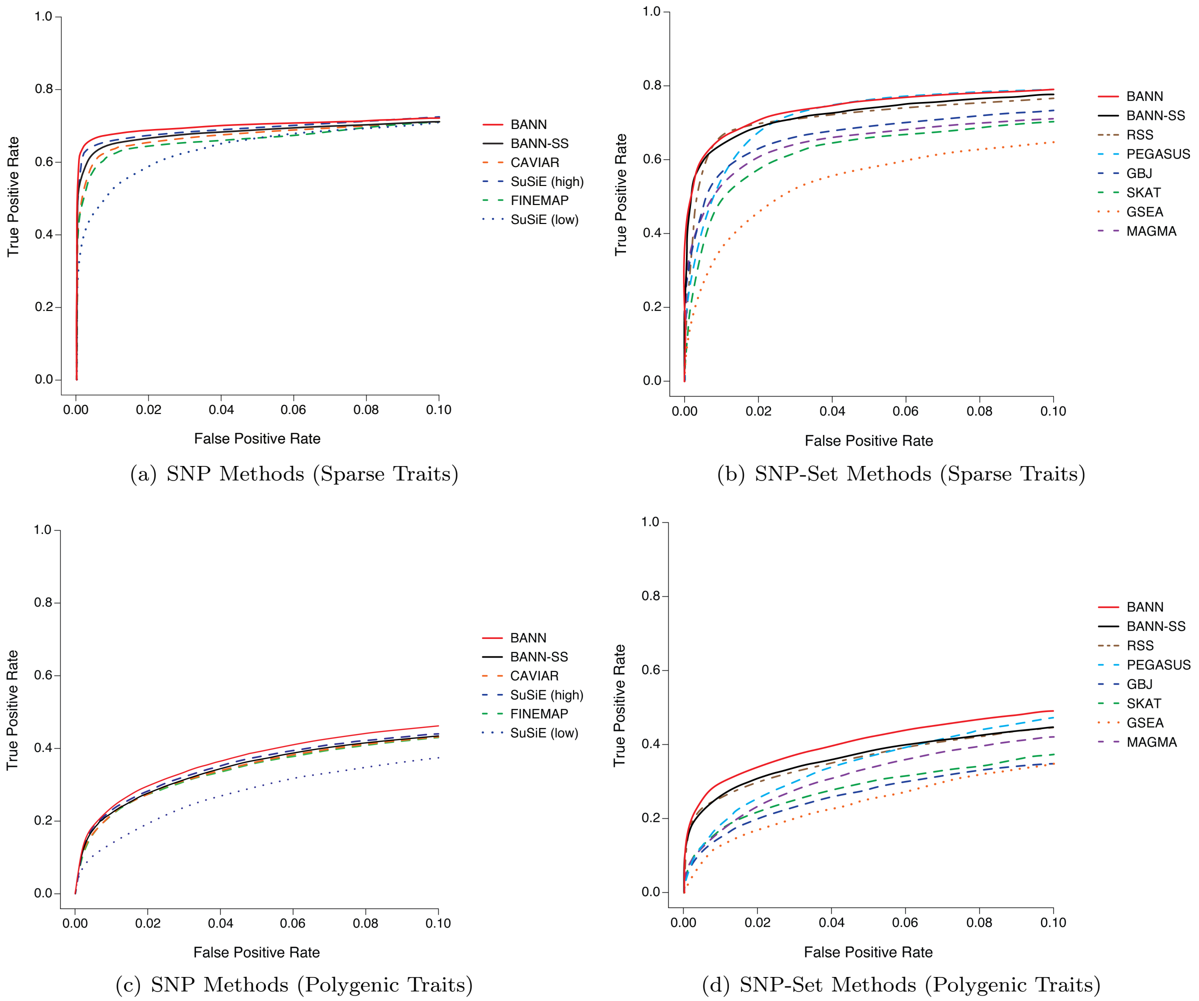

**Figure S5. Receiver operating characteristic (ROC) curves comparing the performance of the BANNs (red) and BANN-SS (black) models with competing SNP and SNP-set mapping approaches in simulations (British cohort).** Here, quantitative traits are simulated to have broad-sense heritability of $H^2 = 0.2$ with equal contributions from additive effects and epistatic interactions (i.e., $\rho = 0.5$). We show power versus false positive rate for two different trait architectures: **(a, b)** sparse where only 1% of SNP-sets are enriched for the trait; and **(c, d)** polygenic where 10% of SNP-sets are enriched. We then set the number of causal SNPs with non-zero effects to be 1% and 10% of all SNPs located within the enriched SNP-sets, respectively. To derive results, the full genotype matrix and phenotypic vector are given to the BANNs model and all competing methods that require individual-level data. For the BANN-SS model and other competing methods that take GWA summary statistics, we compute standard GWA SNP-level effect sizes and $P$-values (estimated using ordinary least squares). **(a, c)** Competing SNP-level mapping approaches include: CAVIAR [63], SuSiE [64], and FINEMAP [65]. The software for SuSiE requires an input $\ell$ which fixes the maximum number of causal SNPs in the model. We display results when this input number is high ($\ell = 3000$) and when this input number is low ($\ell = 10$). **(b, d)** Competing SNP-set mapping approaches include: RSS [7], PEGASUS [66], GBJ [67], SKAT [68], GSEA [69], and MAGMA [70]. Note that the upper limit of the x-axis has been truncated at 0.1. All results are based on 100 replicates (see Supporting Information, Section 9).

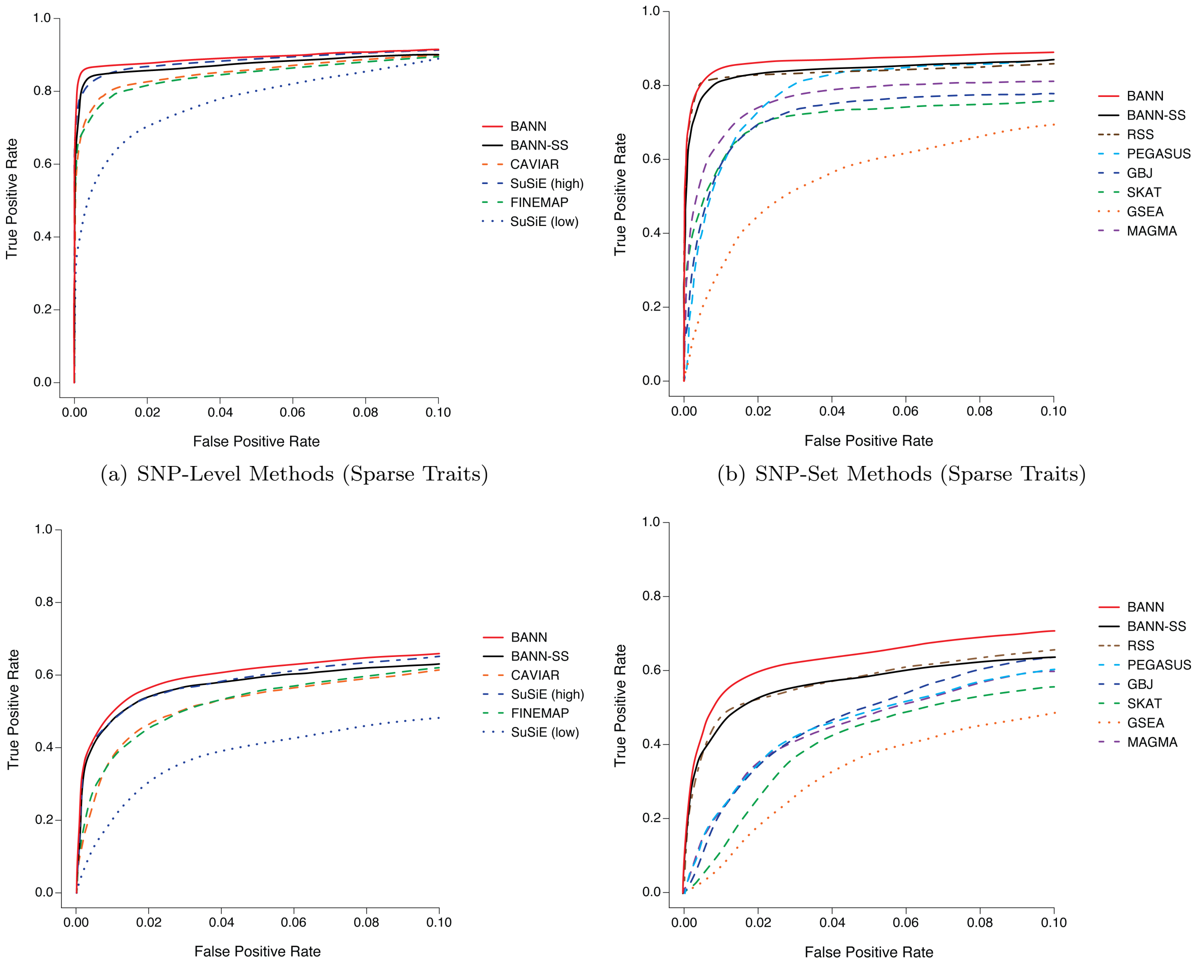

(a) SNP-Level Methods (Sparse Traits)

(b) SNP-Set Methods (Sparse Traits)

(c) SNP-Level Methods (Polygenic Traits)

(d) SNP-Set Methods (Polygenic Traits)

**Figure S6. Receiver operating characteristic (ROC) curves comparing the performance of the BANNs (red) and BANN-SS (black) models with competing SNP and SNP-set mapping approaches in simulations (British cohort).** Here, quantitative traits are simulated to have broad-sense heritability of $H^2 = 0.6$ with equal contributions from additive effects and epistatic interactions (i.e., $\rho = 0.5$). We show power versus false positive rate for two different trait architectures: **(a, b)** sparse where only 1% of SNP-sets are enriched for the trait; and **(c, d)** polygenic where 10% of SNP-sets are enriched. We then set the number of causal SNPs with non-zero effects to be 1% and 10% of all SNPs located within the enriched SNP-sets, respectively. To derive results, the full genotype matrix and phenotypic vector are given to the BANNs model and all competing methods that require individual-level data. For the BANN-SS model and other competing methods that take GWA summary statistics, we compute standard GWA SNP-level effect sizes and $P$-values (estimated using ordinary least squares). **(a, c)** Competing SNP-level mapping approaches include: CAVIAR [63], SuSiE [64], and FINEMAP [65]. The software for SuSiE requires an input $\ell$ which fixes the maximum number of causal SNPs in the model. We display results when this input number is high ($\ell = 3000$) and when this input number is low ($\ell = 10$). **(b, d)** Competing SNP-set mapping approaches include: RSS [7], PEGASUS [66], GBJ [67], SKAT [68], GSEA [69], and MAGMA [70]. Note that the upper limit of the x-axis has been truncated at 0.1. All results are based on 100 replicates (see Supporting Information, Section 9).

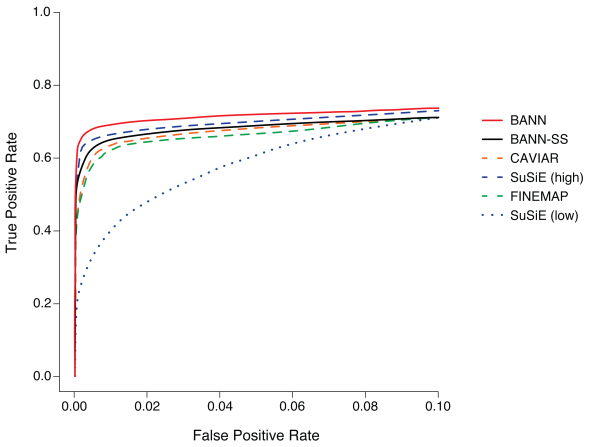

(a) SNP Methods (Sparse Traits)

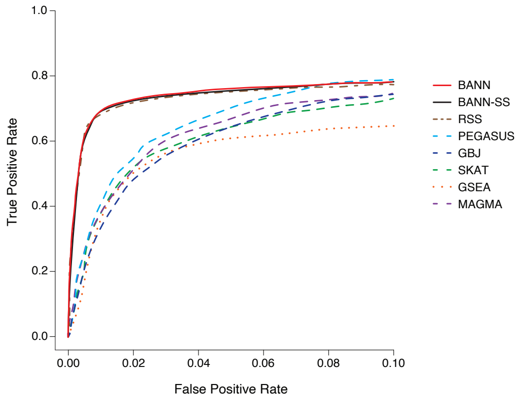

(b) SNP-Set Methods (Sparse Traits)

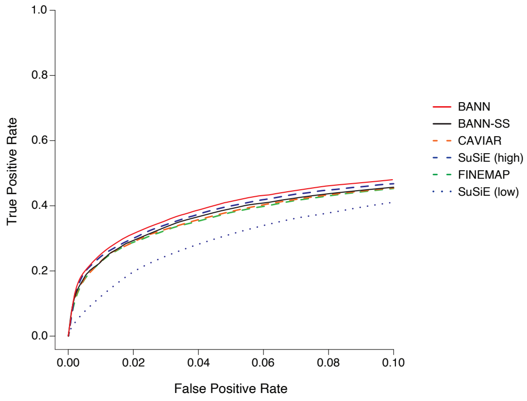

(c) SNP Methods (Polygenic Traits)

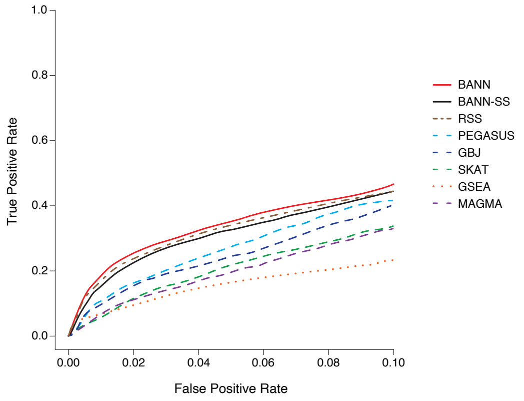

(d) SNP-Set Methods (Polygenic Traits)

**Figure S7. Receiver operating characteristic (ROC) curves comparing the performance of the BANNs (red) and BANN-SS (black) models with competing SNP and SNP-set mapping approaches in simulations with population structure (European cohort).** Here, quantitative traits are simulated to have broad-sense heritability of $H^2 = 0.2$ with equal contributions from additive effects and epistatic interactions (i.e., $\rho = 0.5$). In these simulations, traits were generated while using the top ten principal components (PCs) of the genotype matrix as covariates. We show power versus false positive rate for two different trait architectures: **(a, b)** sparse where only 1% of SNP-sets are enriched for the trait; and **(c, d)** polygenic where 10% of SNP-sets are enriched. We then set the number of causal SNPs with non-zero effects to be 1% and 10% of all SNPs located within the enriched SNP-sets, respectively. To derive results, the full genotype matrix and phenotypic vector are given to the BANNs model and all competing methods that require individual-level data. For the BANN-SS model and other competing methods that take GWA summary statistics, we compute standard GWA SNP-level effect sizes and $P$-values (estimated using ordinary least squares). **(a, c)** Competing SNP-level mapping approaches include: CAVIAR [63], SuSiE [64], and FINEMAP [65]. The software for SuSiE requires an input $\ell$ which fixes the maximum number of causal SNPs in the model. We display results when this input number is high ($\ell = 3000$) and when this input number is low ($\ell = 10$). **(b, d)** Competing SNP-set mapping approaches include: RSS [7], PEGASUS [66], GBJ [67], SKAT [68], GSEA [69], and MAGMA [70]. Note that the upper limit of the x-axis has been truncated at 0.1. All results are based on 100 replicates (see Supporting Information, Section 9).

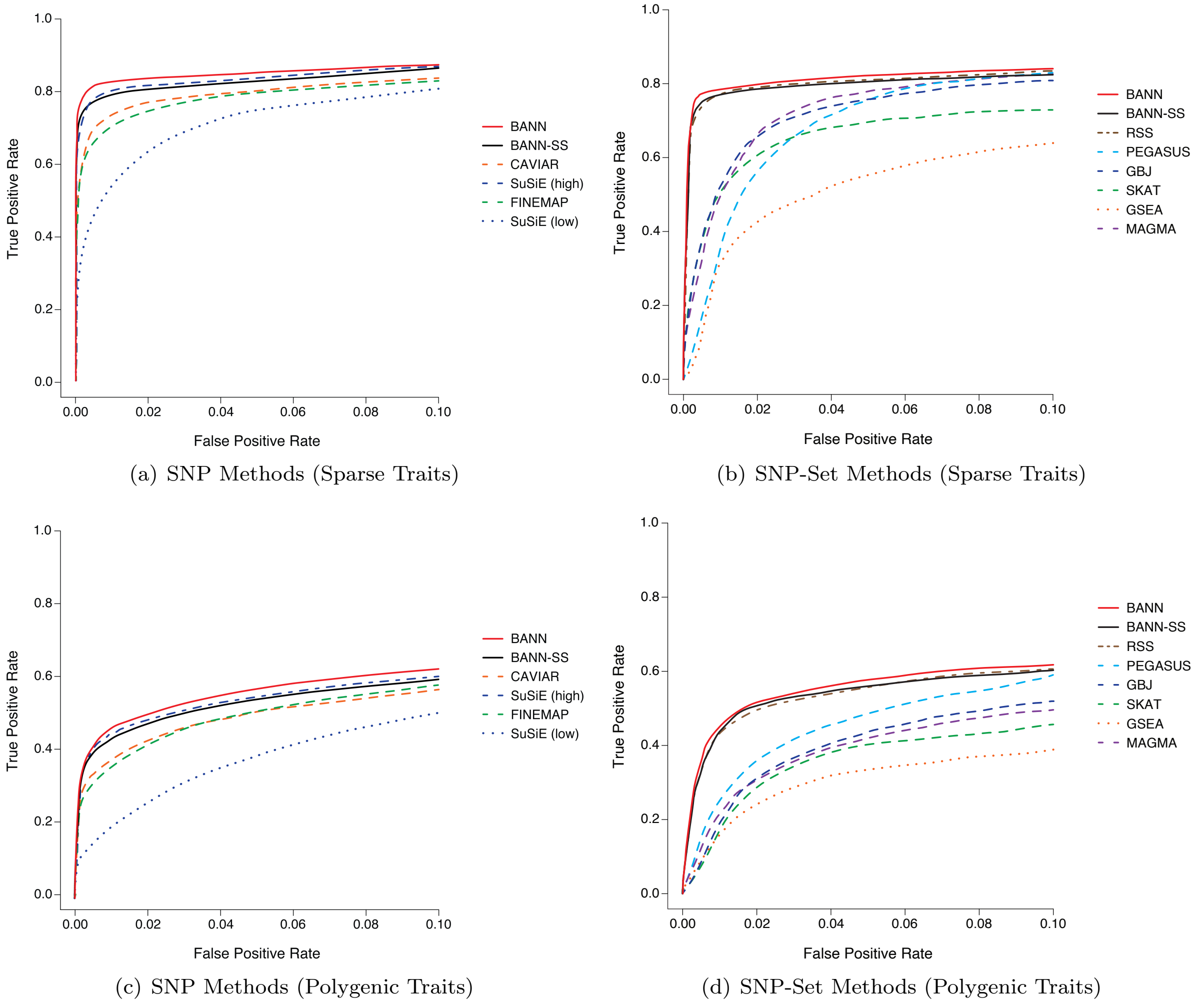

**Figure S8. Receiver operating characteristic (ROC) curves comparing the performance of the BANNs (red) and BANN-SS (black) models with competing SNP and SNP-set mapping approaches in simulations with population structure (European cohort).** Here, quantitative traits are simulated to have broad-sense heritability of $H^2 = 0.6$ with equal contributions from additive effects and epistatic interactions (i.e., $\rho = 0.5$). In these simulations, traits were generated while using the top ten principal components (PCs) of the genotype matrix as covariates. We show power versus false positive rate for two different trait architectures: **(a, b)** sparse where only 1% of SNP-sets are enriched for the trait; and **(c, d)** polygenic where 10% of SNP-sets are enriched. We then set the number of causal SNPs with non-zero effects to be 1% and 10% of all SNPs located within the enriched SNP-sets, respectively. To derive results, the full genotype matrix and phenotypic vector are given to the BANNs model and all competing methods that require individual-level data. For the BANN-SS model and other competing methods that take GWA summary statistics, we compute standard GWA SNP-level effect sizes and $P$-values (estimated using ordinary least squares). **(a, c)** Competing SNP-level mapping approaches include: CAVIAR [63], SuSiE [64], and FINEMAP [65]. The software for SuSiE requires an input $\ell$ which fixes the maximum number of causal SNPs in the model. We display results when this input number is high ($\ell = 3000$) and when this input number is low ($\ell = 10$). **(b, d)** Competing SNP-set mapping approaches include: RSS [7], PEGASUS [66], GBJ [67], SKAT [68], GSEA [69], and MAGMA [70]. Note that the upper limit of the x-axis has been truncated at 0.1. All results are based on 100 replicates (see Supporting Information, Section 9).

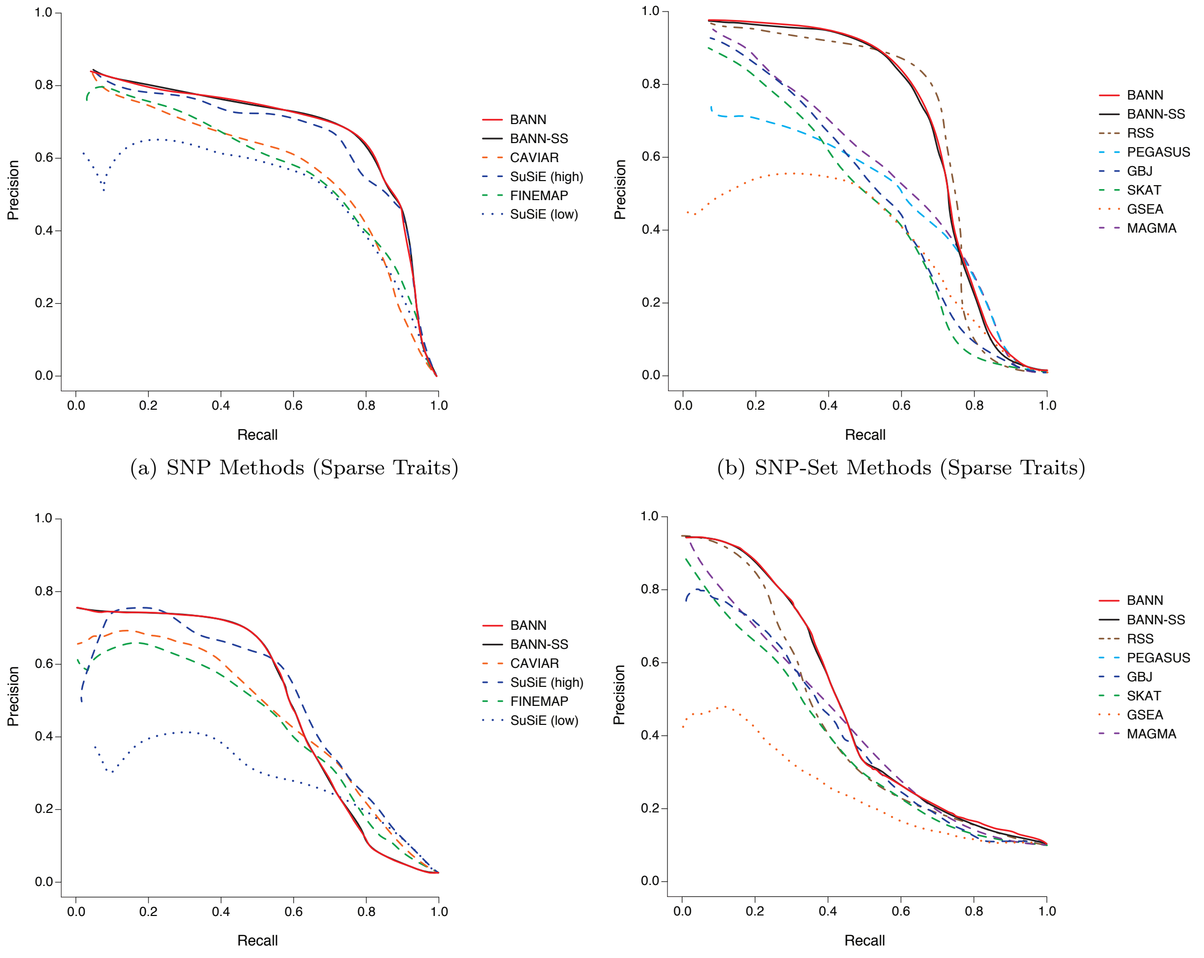

(a) SNP Methods (Sparse Traits)

(b) SNP-Set Methods (Sparse Traits)

(c) SNP Methods (Polygenic Traits)

(d) SNP-Set Methods (Polygenic Traits)

**Figure S9. Precision-recall curves comparing the performance of the BANNs (red) and BANN-SS (black) models with competing SNP and SNP-set mapping approaches in simulations (British cohort).** Here, quantitative traits are simulated to have broad-sense heritability of $H^2 = 0.2$ with only contributions from additive effects (i.e., $\rho = 1$). We show precision versus recall for two different trait architectures: **(a, b)** sparse where only 1% of SNP-sets are enriched for the trait; and **(c, d)** polygenic where 10% of SNP-sets are enriched. We then set the number of causal SNPs with non-zero effects to be 1% and 10% of all SNPs located within the selected enriched SNP-sets, respectively. To derive results, the full genotype matrix and phenotypic vector are given to the BANNs model and all competing methods that require individual-level data. For the BANN-SS model and other competing methods that take GWA summary statistics, we compute standard GWA SNP-level effect sizes and $P$-values (estimated using ordinary least squares). **(a, c)** Competing SNP-level mapping approaches include: CAVIAR [63], SuSiE [64], and FINEMAP [65]. The software for SuSiE requires an input $\ell$ which fixes the maximum number of causal SNPs in the model. We display results when this input number is high ($\ell = 3000$) and when this input number is low ($\ell = 10$). **(b, d)** Competing SNP-set mapping approaches include: RSS [7], PEGASUS [66], GBJ [67], SKAT [68], GSEA [69], and MAGMA [70]. Note that, for traits with sparse architectures, the top ranked SNPs and SNP-sets are always true positives, and therefore the minimal recall is not 0. All results are based on 100 replicates (see Supporting Information, Section 9).

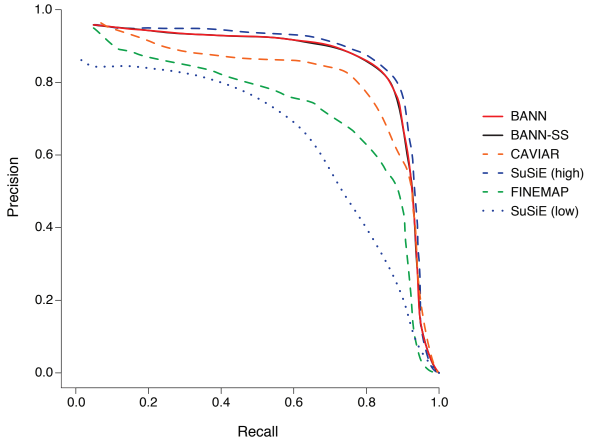

(a) SNP Methods (Sparse Traits)

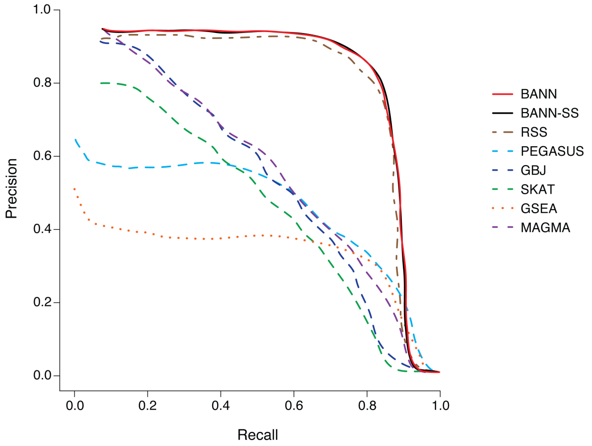

(b) SNP-Set Methods (Sparse Traits)

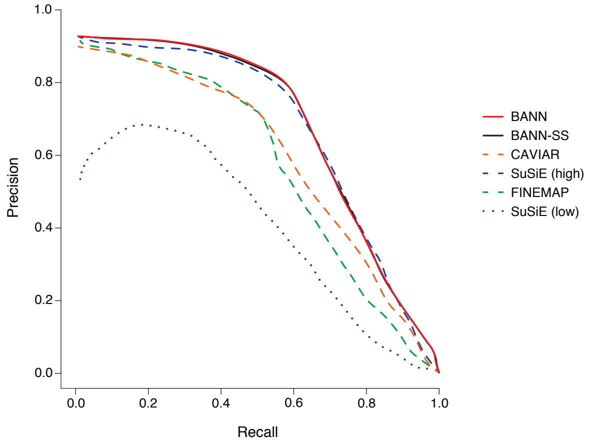

(c) SNP Methods (Polygenic Traits)

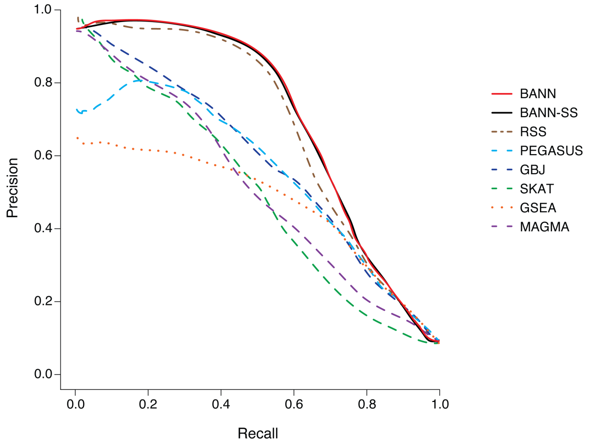

(d) SNP-Set Methods (Polygenic Traits)

**Figure S10. Precision-recall curves comparing the performance of the BANNs (red) and BANN-SS (black) models with competing SNP and SNP-set mapping approaches in simulations (British cohort).** Here, quantitative traits are simulated to have broad-sense heritability of $H^2 = 0.6$ with only contributions from additive effects (i.e., $\rho = 1$). We show precision versus recall for two different trait architectures: **(a, b)** sparse where only 1% of SNP-sets are enriched for the trait; and **(c, d)** polygenic where 10% of SNP-sets are enriched. We then set the number of causal SNPs with non-zero effects to be 1% and 10% of all SNPs located within the selected enriched SNP-sets, respectively. To derive results, the full genotype matrix and phenotypic vector are given to the BANNs model and all competing methods that require individual-level data. For the BANN-SS model and other competing methods that take GWA summary statistics, we compute standard GWA SNP-level effect sizes and $P$-values (estimated using ordinary least squares). **(a, c)** Competing SNP-level mapping approaches include: CAVIAR [63], SuSiE [64], and FINEMAP [65]. The software for SuSiE requires an input $\ell$ which fixes the maximum number of causal SNPs in the model. We display results when this input number is high ($\ell = 3000$) and when this input number is low ($\ell = 10$). **(b, d)** Competing SNP-set mapping approaches include: RSS [7], PEGASUS [66], GBJ [67], SKAT [68], GSEA [69], and MAGMA [70]. Note that, for traits with sparse architectures, the top ranked SNPs and SNP-sets are always true positives, and therefore the minimal recall is not 0. All results are based on 100 replicates (see Supporting Information, Section 9).

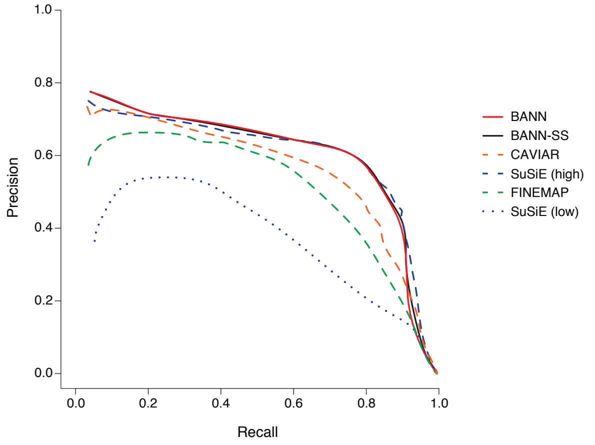

(a) SNP Methods (Sparse Traits)

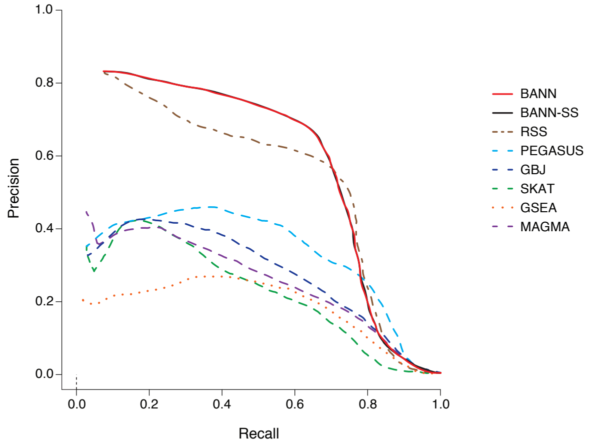

(b) SNP-Set Methods (Sparse Traits)

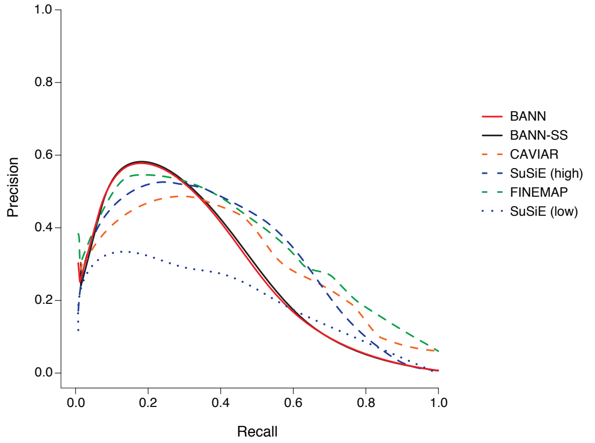

(c) SNP Methods (Polygenic Traits)

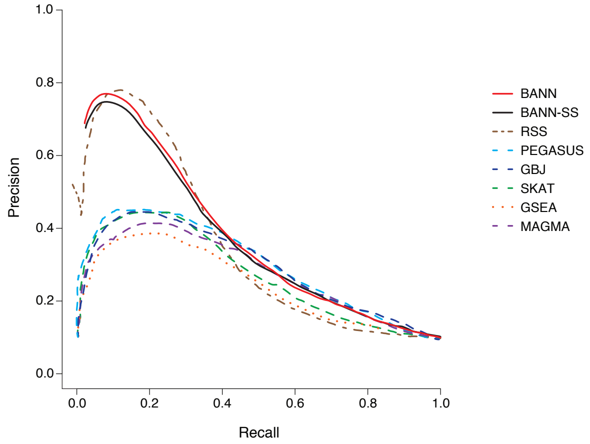

(d) SNP-Set Methods (Polygenic Traits)

**Figure S11. Precision-recall curves comparing the performance of the BANNs (red) and BANN-SS (black) models with competing SNP and SNP-set mapping approaches in simulations with population structure (European cohort).** Here, quantitative traits are simulated to have broad-sense heritability of $H^2 = 0.2$ with only contributions from additive effects (i.e., $\rho = 1$). In these simulations, traits were generated while using the top ten principal components (PCs) of the genotype matrix as covariates. We show precision versus recall for two different trait architectures: **(a, b)** sparse where only 1% of SNP-sets are enriched for the trait; and **(c, d)** polygenic where 10% of SNP-sets are enriched. We then set the number of causal SNPs with non-zero effects to be 1% and 10% of all SNPs located within the selected enriched SNP-sets, respectively. To derive results, the full genotype matrix and phenotypic vector are given to the BANNs model and all competing methods that require individual-level data. For the BANN-SS model and other competing methods that take GWA summary statistics, we compute standard GWA SNP-level effect sizes and $P$-values (estimated using ordinary least squares). **(a, c)** Competing SNP-level mapping approaches include: CAVIAR [63], SuSiE [64], and FINEMAP [65]. The software for SuSiE requires an input $\ell$ which fixes the maximum number of causal SNPs in the model. We display results when this input number is high ($\ell = 3000$) and when this input number is low ($\ell = 10$). **(b, d)** Competing SNP-set mapping approaches include: RSS [7], PEGASUS [66], GBJ [67], SKAT [68], GSEA [69], and MAGMA [70]. Note that, for traits with sparse architectures, the top ranked SNPs and SNP-sets are always true positives, and therefore the minimal recall is not 0. All results are based on 100 replicates (see Supporting Information, Section 9).

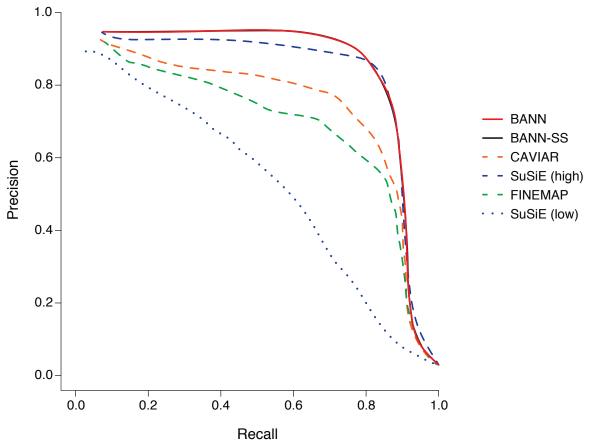

(a) SNP Methods (Sparse Traits)

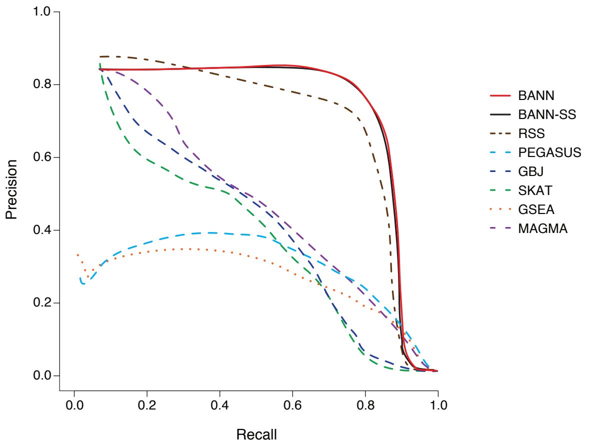

(b) SNP-Set Methods (Sparse Traits)

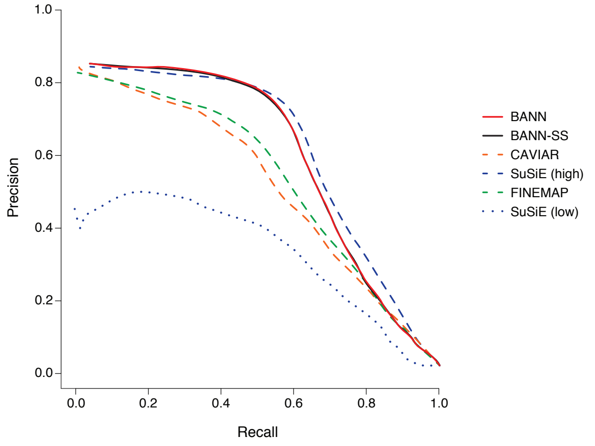

(c) SNP Methods (Polygenic Traits)

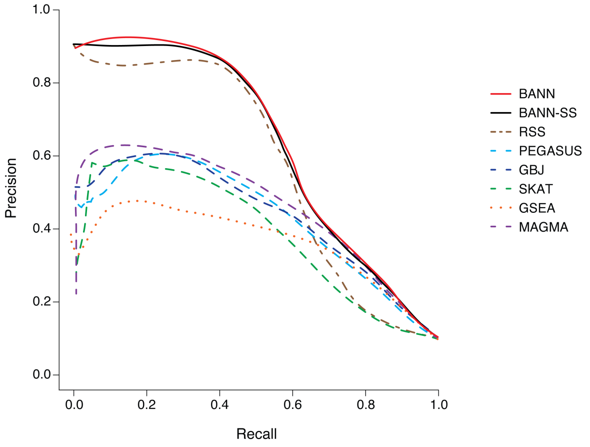

(d) SNP-Set Methods (Polygenic Traits)

**Figure S12. Precision-recall curves comparing the performance of the BANNs (red) and BANN-SS (black) models with competing SNP and SNP-set mapping approaches in simulations with population structure (European cohort).** Here, quantitative traits are simulated to have broad-sense heritability of $H^2 = 0.6$ with only contributions from additive effects (i.e., $\rho = 1$). In these simulations, traits were generated while using the top ten principal components (PCs) of the genotype matrix as covariates. We show precision versus recall for two different trait architectures: **(a, b)** sparse where only 1% of SNP-sets are enriched for the trait; and **(c, d)** polygenic where 10% of SNP-sets are enriched. We then set the number of causal SNPs with non-zero effects to be 1% and 10% of all SNPs located within the selected enriched SNP-sets, respectively. To derive results, the full genotype matrix and phenotypic vector are given to the BANNs model and all competing methods that require individual-level data. For the BANN-SS model and other competing methods that take GWA summary statistics, we compute standard GWA SNP-level effect sizes and $P$-values (estimated using ordinary least squares). **(a, c)** Competing SNP-level mapping approaches include: CAVIAR [63], SuSiE [64], and FINEMAP [65]. The software for SuSiE requires an input $\ell$ which fixes the maximum number of causal SNPs in the model. We display results when this input number is high ($\ell = 3000$) and when this input number is low ($\ell = 10$). **(b, d)** Competing SNP-set mapping approaches include: RSS [7], PEGASUS [66], GBJ [67], SKAT [68], GSEA [69], and MAGMA [70]. Note that, for traits with sparse architectures, the top ranked SNPs and SNP-sets are always true positives, and therefore the minimal recall is not 0. All results are based on 100 replicates (see Supporting Information, Section 9).

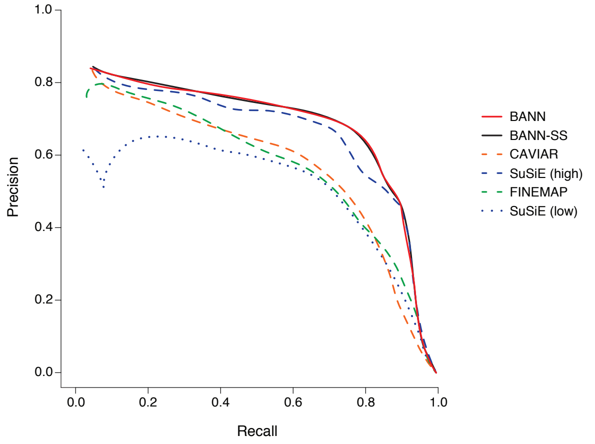

(a) SNP Methods (Sparse Traits)

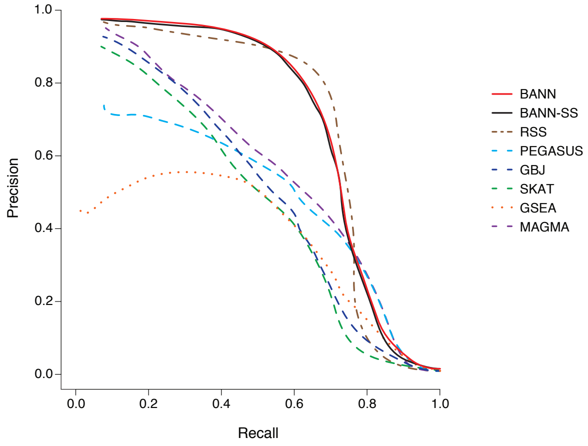

(b) SNP-Set Methods (Sparse Traits)

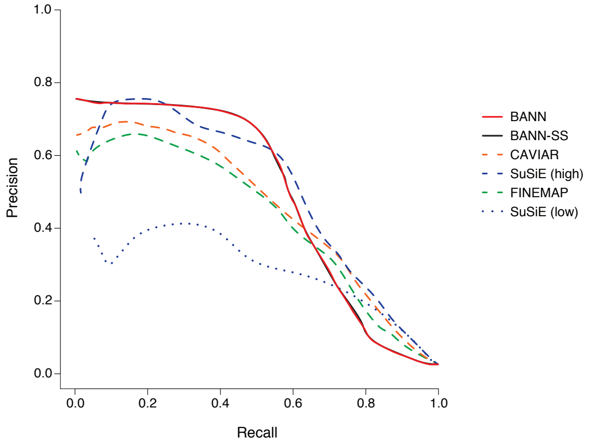

(c) SNP Methods (Polygenic Traits)

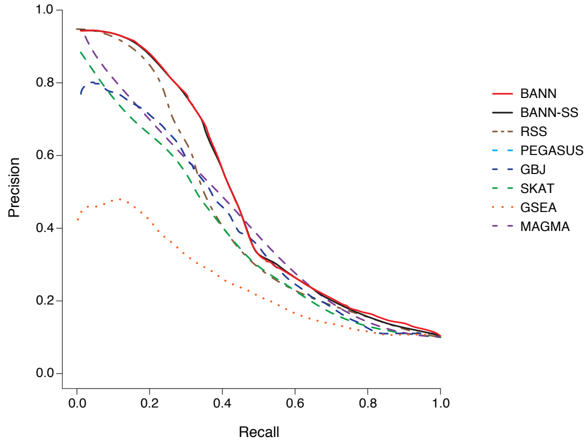

(d) SNP-Set Methods (Polygenic Traits)

**Figure S13. Precision-recall curves comparing the performance of the BANNs (red) and BANN-SS (black) models with competing SNP and SNP-set mapping approaches in simulations (British cohort).** Here, quantitative traits are simulated to have broad-sense heritability of $H^2 = 0.2$ with equal contributions from additive effects and epistatic interactions (i.e., $\rho = 0.5$). We show precision versus recall for two different trait architectures: **(a, b)** sparse where only 1% of SNP-sets are enriched for the trait; and **(c, d)** polygenic where 10% of SNP-sets are enriched. We then set the number of causal SNPs with non-zero effects to be 1% and 10% of all SNPs located within the selected enriched SNP-sets, respectively. To derive results, the full genotype matrix and phenotypic vector are given to the BANNs model and all competing methods that require individual-level data. For the BANN-SS model and other competing methods that take GWA summary statistics, we compute standard GWA SNP-level effect sizes and $P$-values (estimated using ordinary least squares). **(a, c)** Competing SNP-level mapping approaches include: CAVIAR [63], SuSiE [64], and FINEMAP [65]. The software for SuSiE requires an input $\ell$ which fixes the maximum number of causal SNPs in the model. We display results when this input number is high ($\ell = 3000$) and when this input number is low ($\ell = 10$). **(b, d)** Competing SNP-set mapping approaches include: RSS [7], PEGASUS [66], GBJ [67], SKAT [68], GSEA [69], and MAGMA [70]. Note that, for traits with sparse architectures, the top ranked SNPs and SNP-sets are always true positives, and therefore the minimal recall is not 0. All results are based on 100 replicates (see Supporting Information, Section 9).

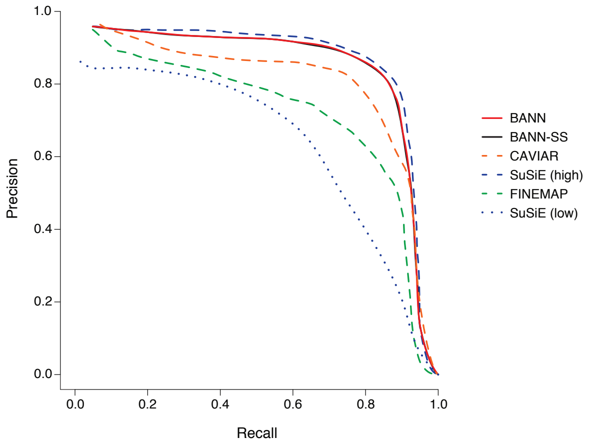

(a) SNP Methods (Sparse Traits)

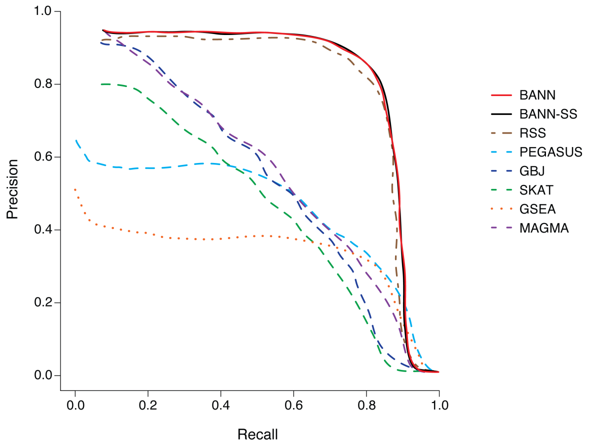

(b) SNP-Set Methods (Sparse Traits)

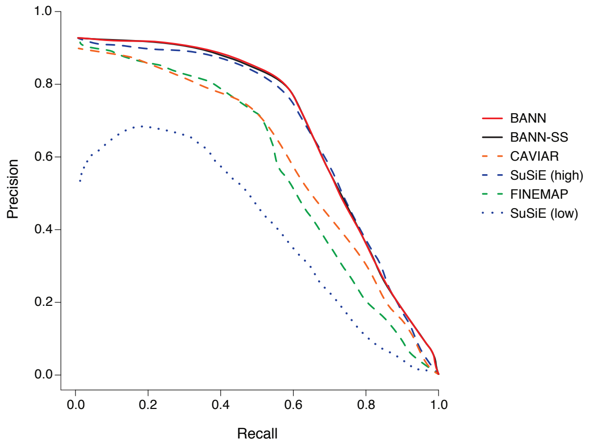

(c) SNP Methods (Polygenic Traits)

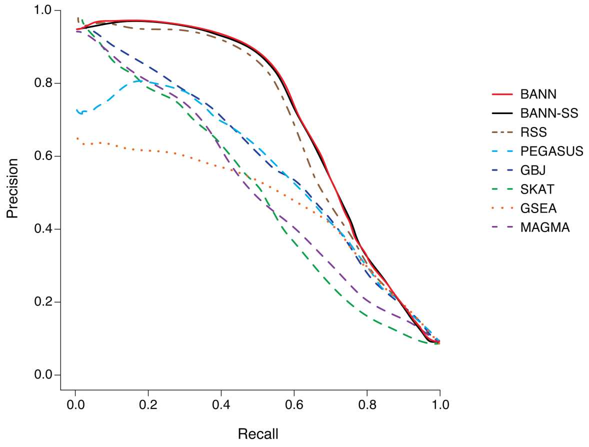

(d) SNP-Set Methods (Polygenic Traits)

**Figure S14. Precision-recall curves comparing the performance of the BANNs (red) and BANN-SS (black) models with competing SNP and SNP-set mapping approaches in simulations (British cohort).** Here, quantitative traits are simulated to have broad-sense heritability of $H^2 = 0.6$ with equal contributions from additive effects and epistatic interactions (i.e., $\rho = 0.5$). We show precision versus recall for two different trait architectures: **(a, b)** sparse where only 1% of SNP-sets are enriched for the trait; and **(c, d)** polygenic where 10% of SNP-sets are enriched. We then set the number of causal SNPs with non-zero effects to be 1% and 10% of all SNPs located within the selected enriched SNP-sets, respectively. To derive results, the full genotype matrix and phenotypic vector are given to the BANNs model and all competing methods that require individual-level data. For the BANN-SS model and other competing methods that take GWA summary statistics, we compute standard GWA SNP-level effect sizes and $P$-values (estimated using ordinary least squares). **(a, c)** Competing SNP-level mapping approaches include: CAVIAR [63], SuSiE [64], and FINEMAP [65]. The software for SuSiE requires an input $\ell$ which fixes the maximum number of causal SNPs in the model. We display results when this input number is high ($\ell = 3000$) and when this input number is low ($\ell = 10$). **(b, d)** Competing SNP-set mapping approaches include: RSS [7], PEGASUS [66], GBJ [67], SKAT [68], GSEA [69], and MAGMA [70]. Note that, for traits with sparse architectures, the top ranked SNPs and SNP-sets are always true positives, and therefore the minimal recall is not 0. All results are based on 100 replicates (see Supporting Information, Section 9).

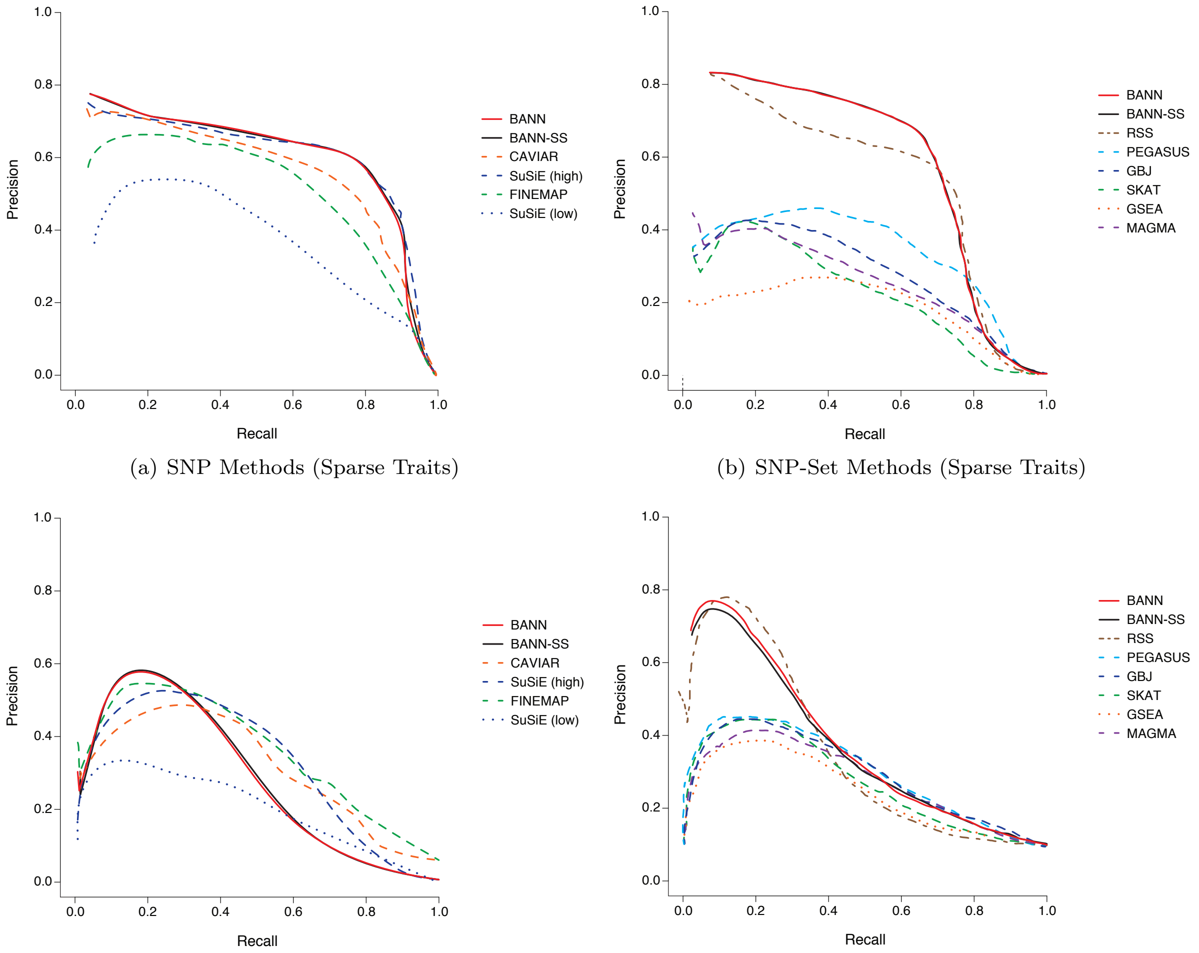

(a) SNP Methods (Sparse Traits)

(b) SNP-Set Methods (Sparse Traits)

(c) SNP Methods (Polygenic Traits)

(d) SNP-Set Methods (Polygenic Traits)

**Figure S15. Precision-recall curves comparing the performance of the BANNs (red) and BANN-SS (black) models with competing SNP and SNP-set mapping approaches in simulations with population structure (European cohort).** Here, quantitative traits are simulated to have broad-sense heritability of $H^2 = 0.2$ with equal contributions from additive effects and epistatic interactions (i.e., $\rho = 0.5$). In these simulations, traits were generated while using the top ten principal components (PCs) of the genotype matrix as covariates. We show precision versus recall for two different trait architectures: **(a, b)** sparse where only 1% of SNP-sets are enriched for the trait; and **(c, d)** polygenic where 10% of SNP-sets are enriched. We then set the number of causal SNPs with non-zero effects to be 1% and 10% of all SNPs located within the selected enriched SNP-sets, respectively. To derive results, the full genotype matrix and phenotypic vector are given to the BANNs model and all competing methods that require individual-level data. For the BANN-SS model and other competing methods that take GWA summary statistics, we compute standard GWA SNP-level effect sizes and $P$-values (estimated using ordinary least squares). **(a, c)** Competing SNP-level mapping approaches include: CAVIAR [63], SuSiE [64], and FINEMAP [65]. The software for SuSiE requires an input $\ell$ which fixes the maximum number of causal SNPs in the model. We display results when this input number is high ($\ell = 3000$) and when this input number is low ($\ell = 10$). **(b, d)** Competing SNP-set mapping approaches include: RSS [7], PEGASUS [66], GBJ [67], SKAT [68], GSEA [69], and MAGMA [70]. Note that, for traits with sparse architectures, the top ranked SNPs and SNP-sets are always true positives, and therefore the minimal recall is not 0. All results are based on 100 replicates (see Supporting Information, Section 9).

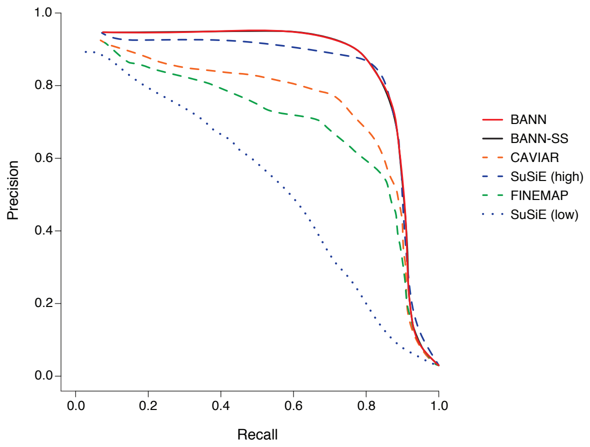

(a) SNP Methods (Sparse Traits)

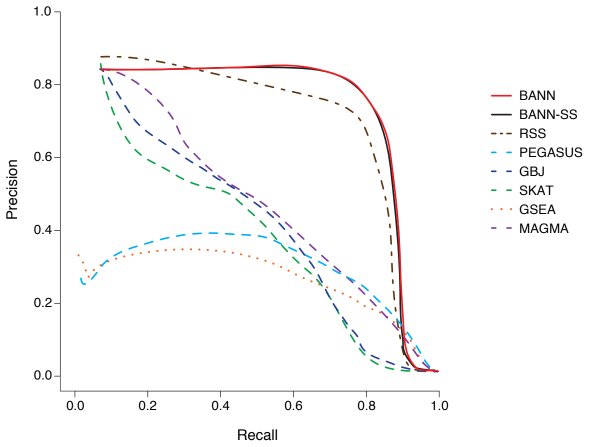

(b) SNP-Set Methods (Sparse Traits)

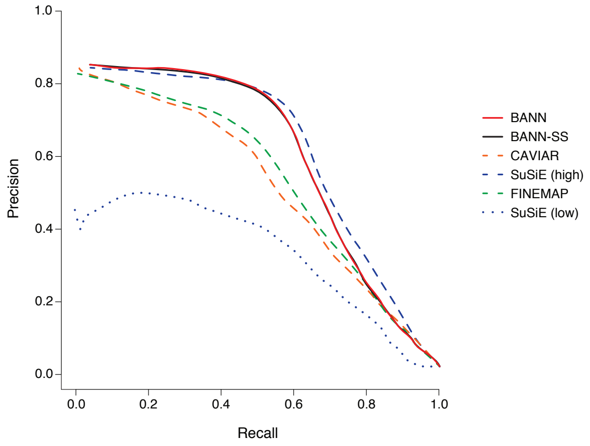

(c) SNP Methods (Polygenic Traits)

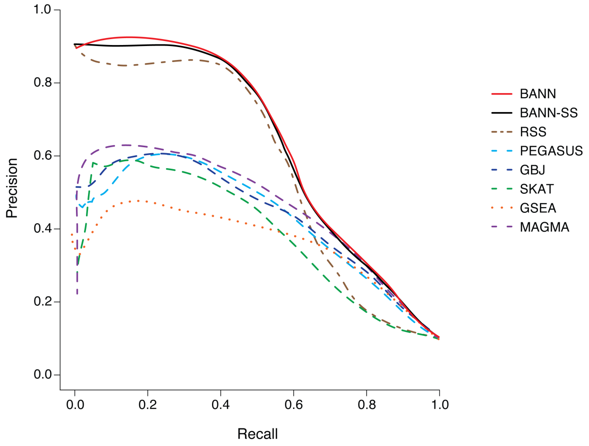

(d) SNP-Set Methods (Polygenic Traits)

**Figure S16. Precision-recall curves comparing the performance of the BANNs (red) and BANN-SS (black) models with competing SNP and SNP-set mapping approaches in simulations with population structure (European cohort).** Here, quantitative traits are simulated to have broad-sense heritability of $H^2 = 0.2$ with equal contributions from additive effects and epistatic interactions (i.e., $\rho = 0.5$). In these simulations, traits were generated while using the top ten principal components (PCs) of the genotype matrix as covariates. We show precision versus recall for two different trait architectures: **(a, b)** sparse where only 1% of SNP-sets are enriched for the trait; and **(c, d)** polygenic where 10% of SNP-sets are enriched. We then set the number of causal SNPs with non-zero effects to be 1% and 10% of all SNPs located within the selected enriched SNP-sets, respectively. To derive results, the full genotype matrix and phenotypic vector are given to the BANNs model and all competing methods that require individual-level data. For the BANN-SS model and other competing methods that take GWA summary statistics, we compute standard GWA SNP-level effect sizes and $P$-values (estimated using ordinary least squares). **(a, c)** Competing SNP-level mapping approaches include: CAVIAR [63], SuSiE [64], and FINEMAP [65]. The software for SuSiE requires an input $\ell$ which fixes the maximum number of causal SNPs in the model. We display results when this input number is high ($\ell = 3000$) and when this input number is low ($\ell = 10$). **(b, d)** Competing SNP-set mapping approaches include: RSS [7], PEGASUS [66], GBJ [67], SKAT [68], GSEA [69], and MAGMA [70]. Note that, for traits with sparse architectures, the top ranked SNPs and SNP-sets are always true positives, and therefore the minimal recall is not 0. All results are based on 100 replicates (see Supporting Information, Section 9).

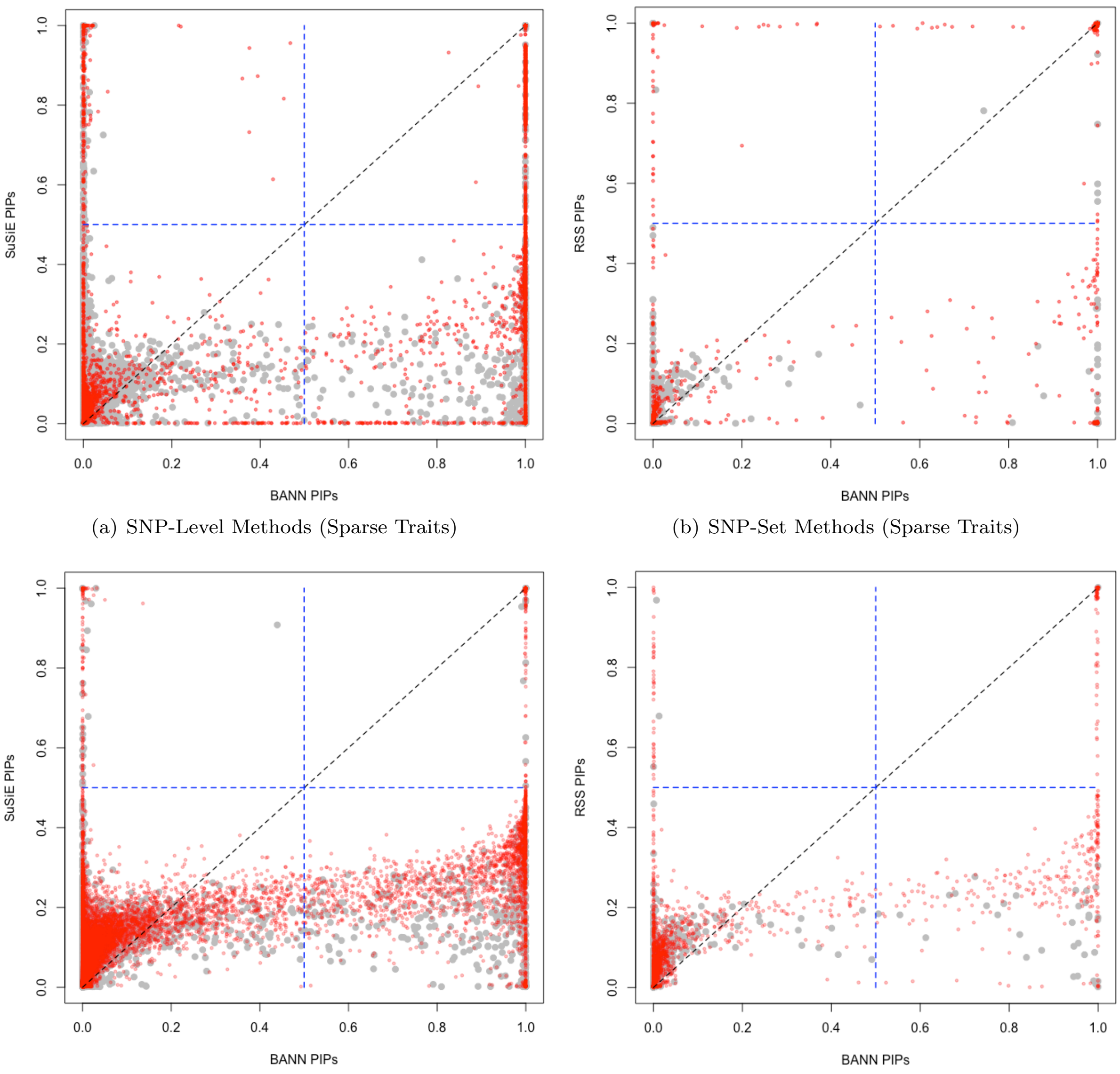

(a) SNP-Level Methods (Sparse Traits)

(b) SNP-Set Methods (Sparse Traits)

(c) SNP-Level Methods (Polygenic Traits)

(d) SNP-Set Methods (Polygenic Traits)

**Figure S17. Scatter plots comparing how the integrative neural network training procedure enables the ability to identify associated SNPs and enriched SNP-sets in simulations (British cohort).** Quantitative traits are simulated to have broad-sense heritability of $H^2 = 0.2$ with only contributions from additive effects set (i.e., $\rho = 1$). We consider two different trait architectures: **(a, b)** sparse where only 1% of SNP-sets are enriched for the trait; and **(c, d)** polygenic where 10% of SNP-sets are enriched. We set the number of causal SNPs with non-zero effects to be $\sim 1\%$ and $\sim 10\%$ of all SNPs located within the enriched SNP-sets, respectively. Results are shown comparing the posterior inclusion probabilities (PIPs) derived by the BANNs model fit with individual-level data on the x-axis and **(a, c)** SuSiE [64] and **(b, d)** RSS [7] on the y-axis, respectively. Here, SuSie is fit while assuming a high maximum number of causal SNPs ($\ell = 3000$). The blue horizontal and vertical dashed lines are marked at the "median probability criterion" (i.e., PIPs for SNPs and SNP-sets greater than 0.5) [51]. True positive causal variants used to generate the synthetic phenotypes are colored in red, while non-causal variants are given in grey. SNPs and SNP-sets in the top right quadrant are selected by both approaches; while, elements in the bottom right and top left quadrants are uniquely identified by BANNs and SuSie/RSS, respectively. Each plot combines results from 100 simulated replicates (see Section 9).

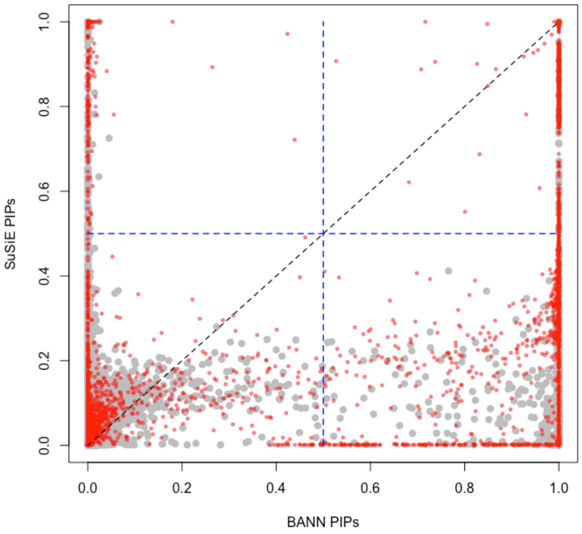

(a) SNP-Level Methods (Sparse Traits)

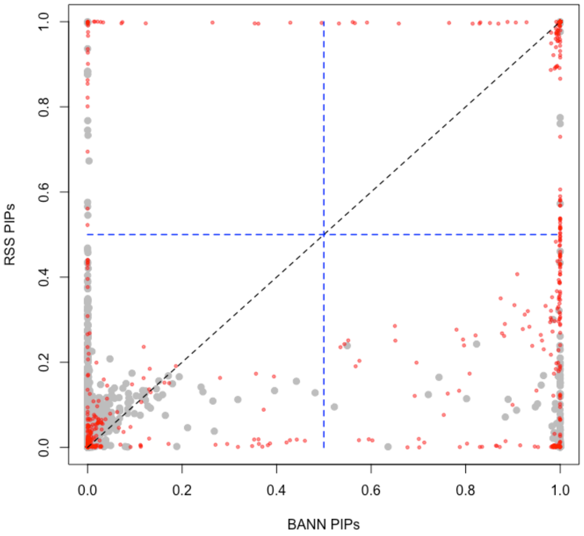

(b) SNP-Set Methods (Sparse Traits)

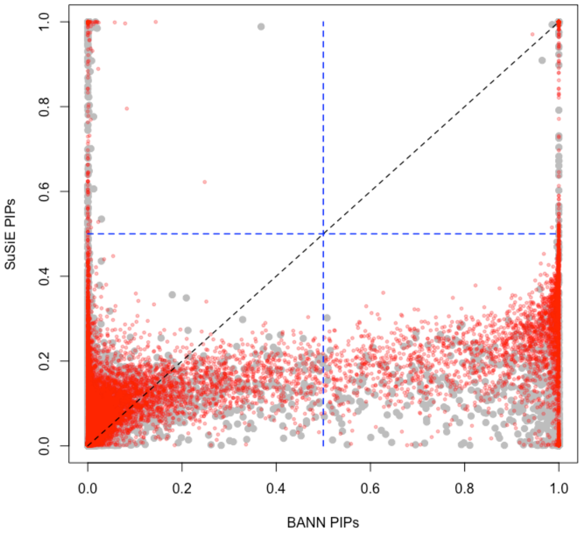

(c) SNP-Level Methods (Polygenic Traits)

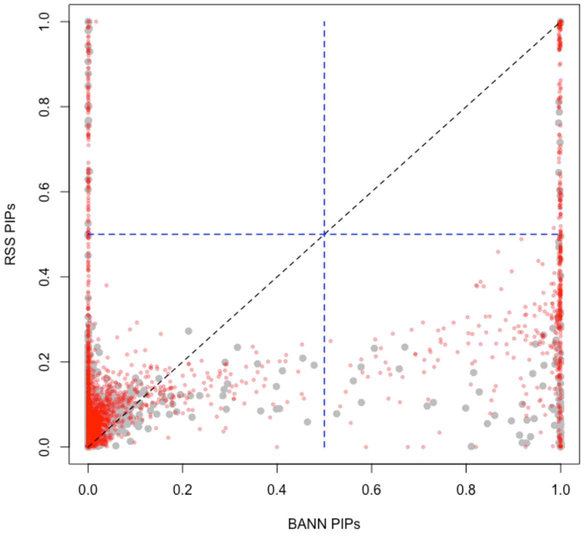

(d) SNP-Set Methods (Polygenic Traits)

**Figure S18. Scatter plots comparing how the integrative neural network training procedure enables the ability to identify associated SNPs and enriched SNP-sets in simulations with population structure (European cohort).** Quantitative traits are simulated to have broad-sense heritability of $H^2 = 0.2$ with only contributions from additive effects set (i.e., $\rho = 1$). We consider two different trait architectures: **(a, b)** sparse where only 1% of SNP-sets are enriched for the trait; and **(c, d)** polygenic where 10% of SNP-sets are enriched. We set the number of causal SNPs with non-zero effects to be 1% and 10% of all SNPs located within the enriched SNP-sets, respectively. In these simulations, traits were generated while also using the top ten principal components (PCs) of the genotype matrix as covariates. Results are shown comparing the posterior inclusion probabilities (PIPs) derived by the BANNs model fit with individual-level data on the x-axis and **(a, c)** SuSiE [64] and **(b, d)** RSS [7] on the y-axis, respectively. Here, SuSie is fit while assuming a high maximum number of causal SNPs ($\ell = 3000$). The blue horizontal and vertical dashed lines are marked at the "median probability criterion" (i.e., PIPs for SNPs and SNP-sets greater than 0.5) [51]. True positive causal variants used to generate the synthetic phenotypes are colored in red, while non-causal variants are given in grey. SNPs and SNP-sets in the top right quadrant are selected by both approaches; while, elements in the bottom right and top left quadrants are uniquely identified by BANNs and SuSie/RSS, respectively. Each plot combines results from 100 simulated replicates (see Section 9).

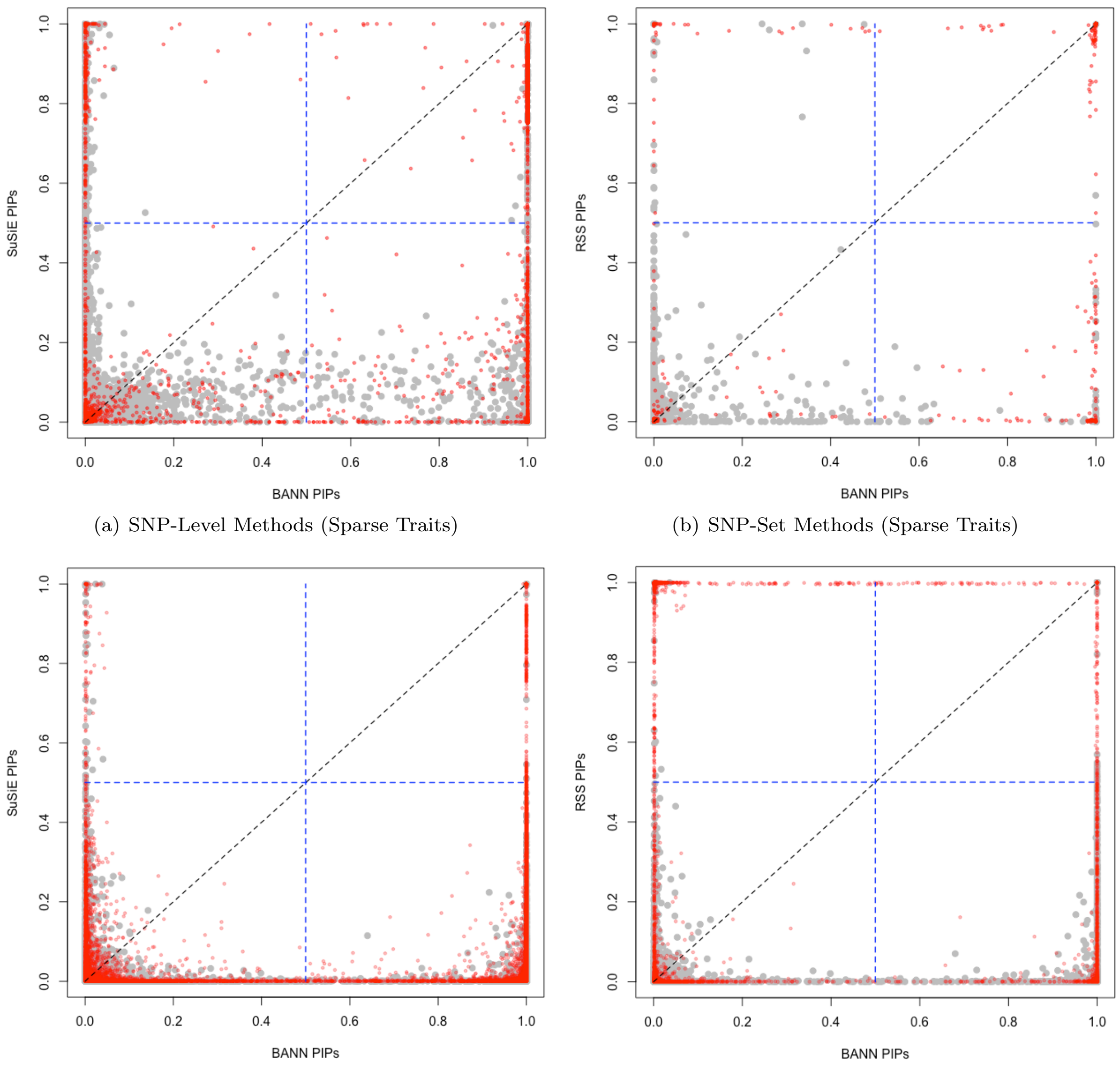

(a) SNP-Level Methods (Sparse Traits)

(b) SNP-Set Methods (Sparse Traits)

(c) SNP-Level Methods (Polygenic Traits)

(d) SNP-Set Methods (Polygenic Traits)

**Figure S19. Scatter plots comparing how the integrative neural network training procedure enables the ability to identify associated SNPs and enriched SNP-sets in simulations with population structure (European cohort).** Quantitative traits are simulated to have broad-sense heritability of $H^2 = 0.6$ with only contributions from additive effects set (i.e., $\rho = 1$). We consider two different trait architectures: **(a, b)** sparse where only 1% of SNP-sets are enriched for the trait; and **(c, d)** polygenic where 10% of SNP-sets are enriched. We set the number of causal SNPs with non-zero effects to be 1% and 10% of all SNPs located within the enriched SNP-sets, respectively. In these simulations, traits were generated while also using the top ten principal components (PCs) of the genotype matrix as covariates. Results are shown comparing the posterior inclusion probabilities (PIPs) derived by the BANNs model fit with individual-level data on the x-axis and **(a, c)** SuSiE [64] and **(b, d)** RSS [7] on the y-axis, respectively. Here, SuSie is fit while assuming a high maximum number of causal SNPs ($\ell = 3000$). The blue horizontal and vertical dashed lines are marked at the "median probability criterion" (i.e., PIPs for SNPs and SNP-sets greater than 0.5) [51]. True positive causal variants used to generate the synthetic phenotypes are colored in red, while non-causal variants are given in grey. SNPs and SNP-sets in the top right quadrant are selected by both approaches; while, elements in the bottom right and top left quadrants are uniquely identified by BANNs and SuSie/RSS, respectively. Each plot combines results from 100 simulated replicates (see Section 9).

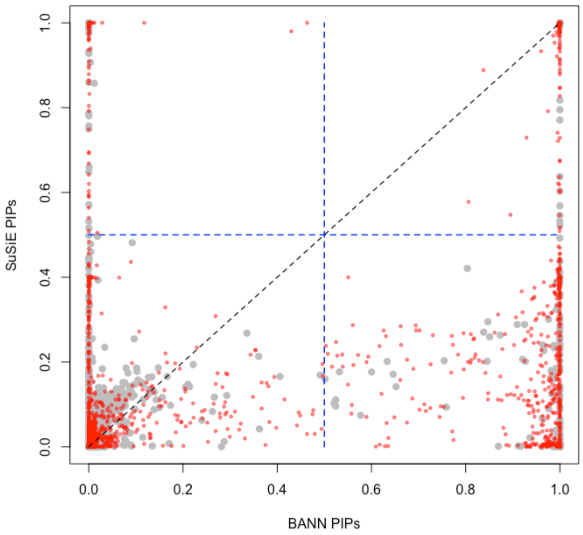

(a) SNP-Level Methods (Sparse Traits)

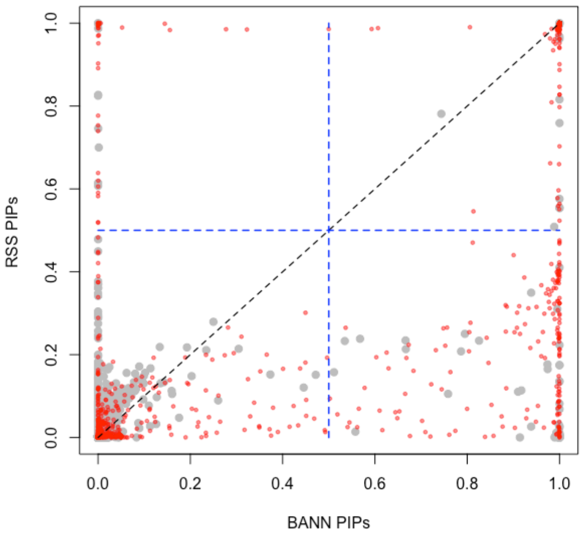

(b) SNP-Set Methods (Sparse Traits)

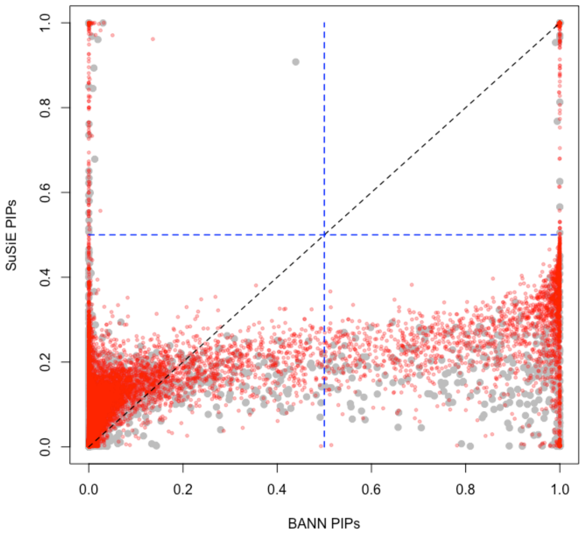

(c) SNP-Level Methods (Polygenic Traits)

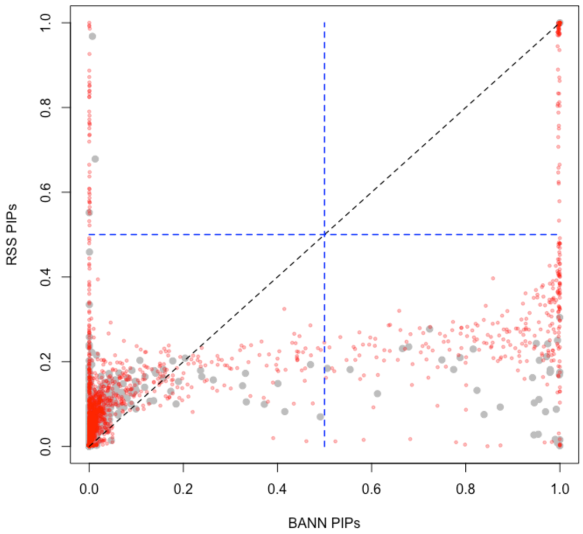

(d) SNP-Set Methods (Polygenic Traits)

**Figure S20. Scatter plots comparing how the integrative neural network training procedure enables the ability to identify associated SNPs and enriched SNP-sets in simulations (British cohort).** Quantitative traits are simulated to have broad-sense heritability of $H^2 = 0.2$ with equal contributions from additive effects and epistatic interactions (i.e., $\rho = 0.5$). We consider two different trait architectures: **(a, b)** sparse where only 1% of SNP-sets are enriched for the trait; and **(c, d)** polygenic where 10% of SNP-sets are enriched. We set the number of causal SNPs with non-zero effects to be 1% and 10% of all SNPs located within the enriched SNP-sets, respectively. Results are shown comparing the posterior inclusion probabilities (PIPs) derived by the BANNs model fit with individual-level data on the x-axis and **(a, c)** SuSiE [64] and **(b, d)** RSS [7] on the y-axis, respectively. Here, SuSie is fit while assuming a high maximum number of causal SNPs ($\ell = 3000$). The blue horizontal and vertical dashed lines are marked at the "median probability criterion" (i.e., PIPs for SNPs and SNP-sets greater than 0.5) [51]. True positive causal variants used to generate the synthetic phenotypes are colored in red, while non-causal variants are given in grey. SNPs and SNP-sets in the top right quadrant are selected by both approaches; while, elements in the bottom right and top left quadrants are uniquely identified by BANNs and SuSie/RSS, respectively. Each plot combines results from 100 simulated replicates (see Section 9).

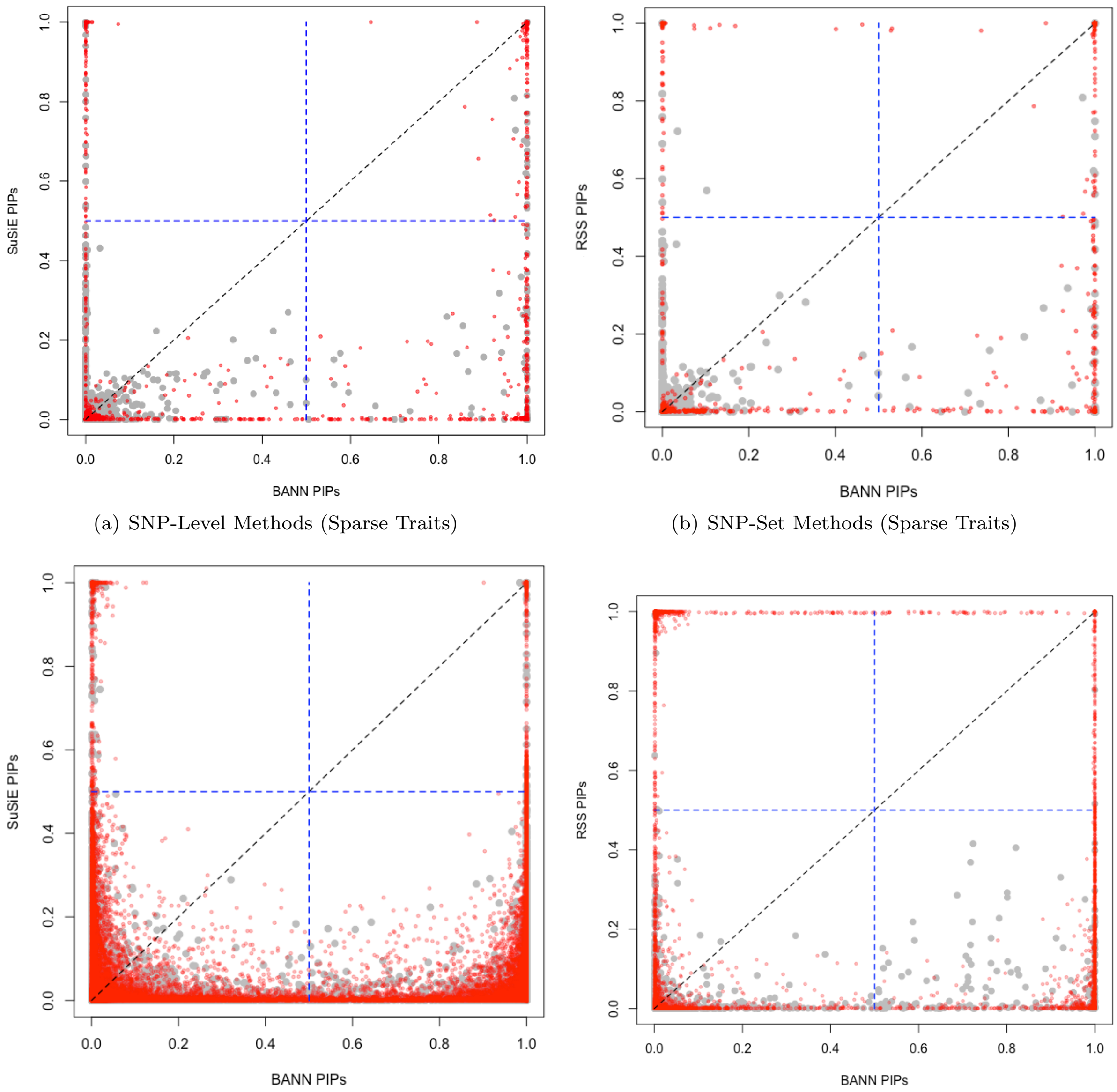

(a) SNP-Level Methods (Sparse Traits)

(b) SNP-Set Methods (Sparse Traits)

(c) SNP-Level Methods (Polygenic Traits)

(d) SNP-Set Methods (Polygenic Traits)

**Figure S21. Scatter plots comparing how the integrative neural network training procedure enables the ability to identify associated SNPs and enriched SNP-sets in simulations (British cohort).** Quantitative traits are simulated to have broad-sense heritability of $H^2 = 0.6$ with equal contributions from additive effects and epistatic interactions (i.e., $\rho = 0.5$). We consider two different trait architectures: **(a, b)** sparse where only 1% of SNP-sets are enriched for the trait; and **(c, d)** polygenic where 10% of SNP-sets are enriched. We set the number of causal SNPs with non-zero effects to be 1% and 10% of all SNPs located within the enriched SNP-sets, respectively. Results are shown comparing the posterior inclusion probabilities (PIPs) derived by the BANNs model fit with individual-level data on the x-axis and **(a, c)** SuSiE [64] and **(b, d)** RSS [7] on the y-axis, respectively. Here, SuSie is fit while assuming a high maximum number of causal SNPs ($\ell = 3000$). The blue horizontal and vertical dashed lines are marked at the "median probability criterion" (i.e., PIPs for SNPs and SNP-sets greater than 0.5) [51]. True positive causal variants used to generate the synthetic phenotypes are colored in red, while non-causal variants are given in grey. SNPs and SNP-sets in the top right quadrant are selected by both approaches; while, elements in the bottom right and top left quadrants are uniquely identified by BANNs and SuSie/RSS, respectively. Each plot combines results from 100 simulated replicates (see Section 9).

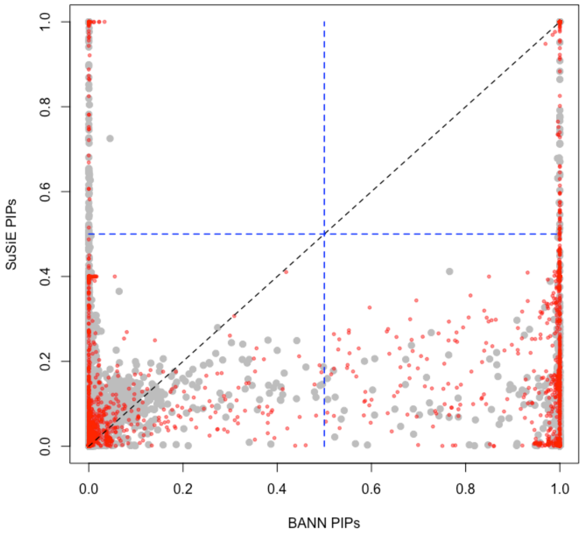

(a) SNP-Level Methods (Sparse Traits)

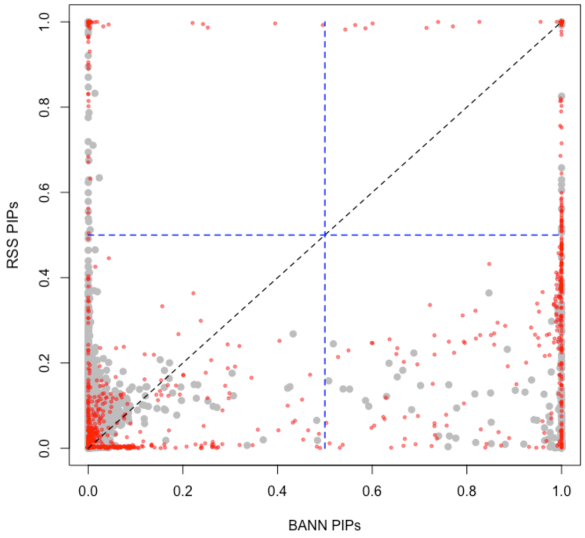

(b) SNP-Set Methods (Sparse Traits)

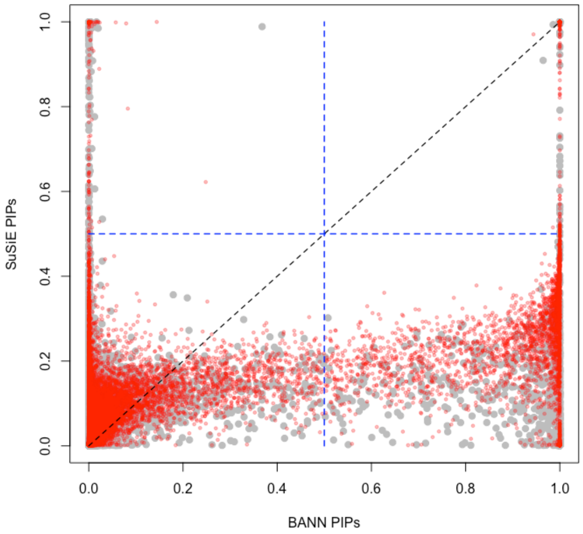

(c) SNP-Level Methods (Polygenic Traits)

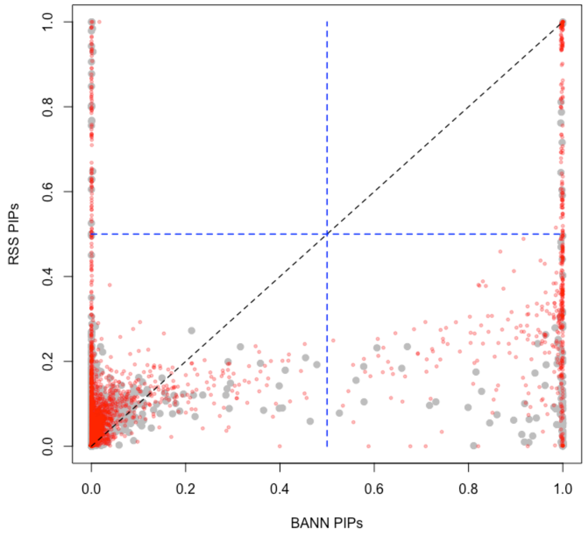

(d) SNP-Set Methods (Polygenic Traits)

**Figure S22. Scatter plots comparing how the integrative neural network training procedure enables the ability to identify associated SNPs and enriched SNP-sets in simulations with population structure (European cohort).** Quantitative traits are simulated to have broad-sense heritability of $H^2 = 0.2$ with equal contributions from additive effects and epistatic interactions (i.e., $\rho = 0.5$). We consider two different trait architectures: **(a, b)** sparse where only 1% of SNP-sets are enriched for the trait; and **(c, d)** polygenic where 10% of SNP-sets are enriched. We set the number of causal SNPs with non-zero effects to be 1% and 10% of all SNPs located within the enriched SNP-sets, respectively. In these simulations, traits were generated while also using the top ten principal components (PCs) of the genotype matrix as covariates. Results are shown comparing the posterior inclusion probabilities (PIPs) derived by the BANNs model fit with individual-level data on the x-axis and **(a, c)** SuSiE [64] and **(b, d)** RSS [7] on the y-axis, respectively. Here, SuSie is fit while assuming a high maximum number of causal SNPs ($\ell = 3000$). The blue horizontal and vertical dashed lines are marked at the "median probability criterion" (i.e., PIPs for SNPs and SNP-sets greater than 0.5) [51]. True positive causal variants used to generate the synthetic phenotypes are colored in red, while non-causal variants are given in grey. SNPs and SNP-sets in the top right quadrant are selected by both approaches; while, elements in the bottom right and top left quadrants are uniquely identified by BANNs and SuSie/RSS, respectively. Each plot combines results from 100 simulated replicates (see Section 9).

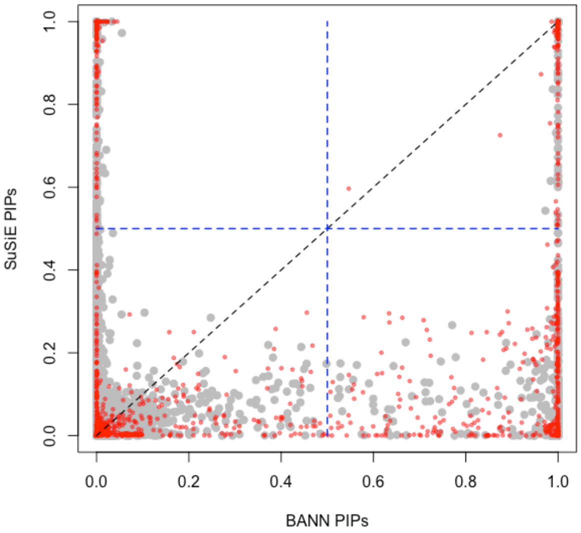

(a) SNP-Level Methods (Sparse Traits)

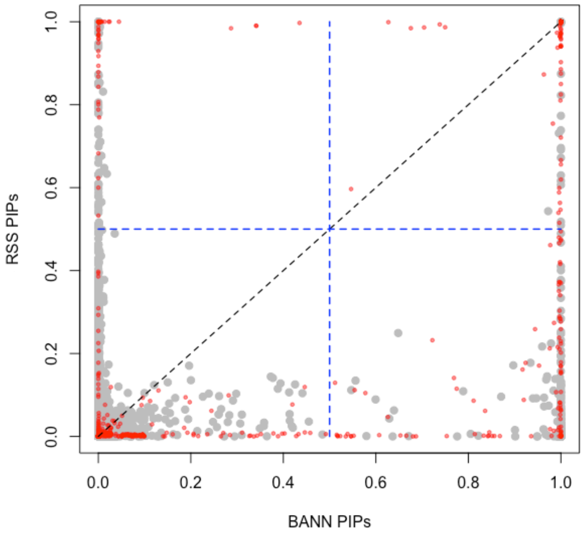

(b) SNP-Set Methods (Sparse Traits)

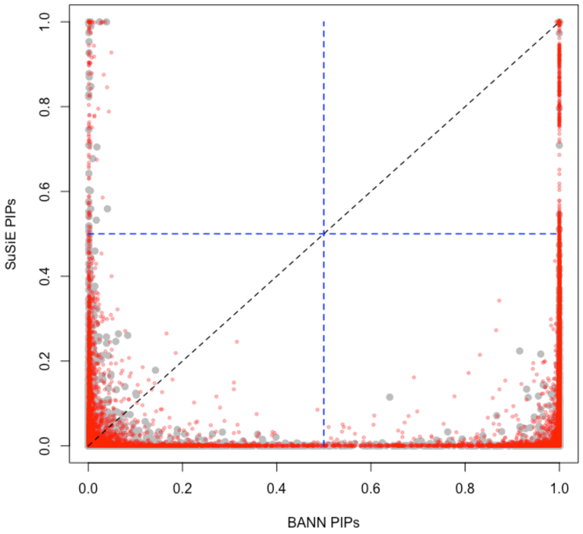

(c) SNP-Level Methods (Polygenic Traits)

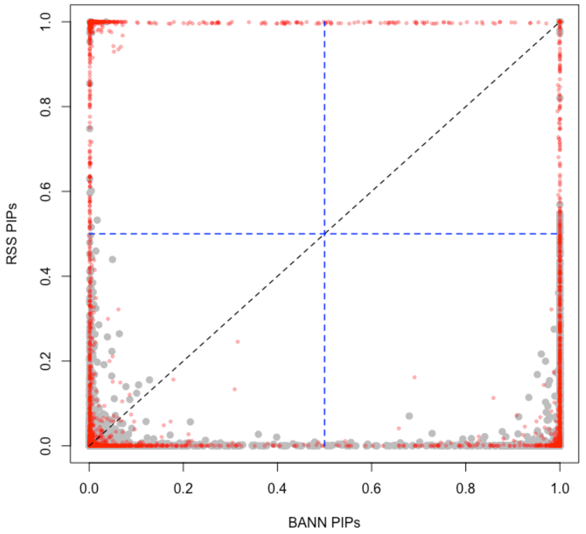

(d) SNP-Set Methods (Polygenic Traits)

**Figure S23. Scatter plots comparing how the integrative neural network training procedure enables the ability to identify associated SNPs and enriched SNP-sets in simulations with population structure (European cohort).** Quantitative traits are simulated to have broad-sense heritability of $H^2 = 0.6$ with equal contributions from additive effects and epistatic interactions (i.e., $\rho = 0.5$). We consider two different trait architectures: **(a, b)** sparse where only 1% of SNP-sets are enriched for the trait; and **(c, d)** polygenic where 10% of SNP-sets are enriched. We set the number of causal SNPs with non-zero effects to be 1% and 10% of all SNPs located within the enriched SNP-sets, respectively. In these simulations, traits were generated while also using the top ten principal components (PCs) of the genotype matrix as covariates. Results are shown comparing the posterior inclusion probabilities (PIPs) derived by the BANNs model fit with individual-level data on the x-axis and **(a, c)** SuSiE [64] and **(b, d)** RSS [7] on the y-axis, respectively. Here, SuSie is fit while assuming a high maximum number of causal SNPs ($\ell = 3000$). The blue horizontal and vertical dashed lines are marked at the "median probability criterion" (i.e., PIPs for SNPs and SNP-sets greater than 0.5) [51]. True positive causal variants used to generate the synthetic phenotypes are colored in red, while non-causal variants are given in grey. SNPs and SNP-sets in the top right quadrant are selected by both approaches; while, elements in the bottom right and top left quadrants are uniquely identified by BANNs and SuSie/RSS, respectively. Each plot combines results from 100 simulated replicates (see Section 9).

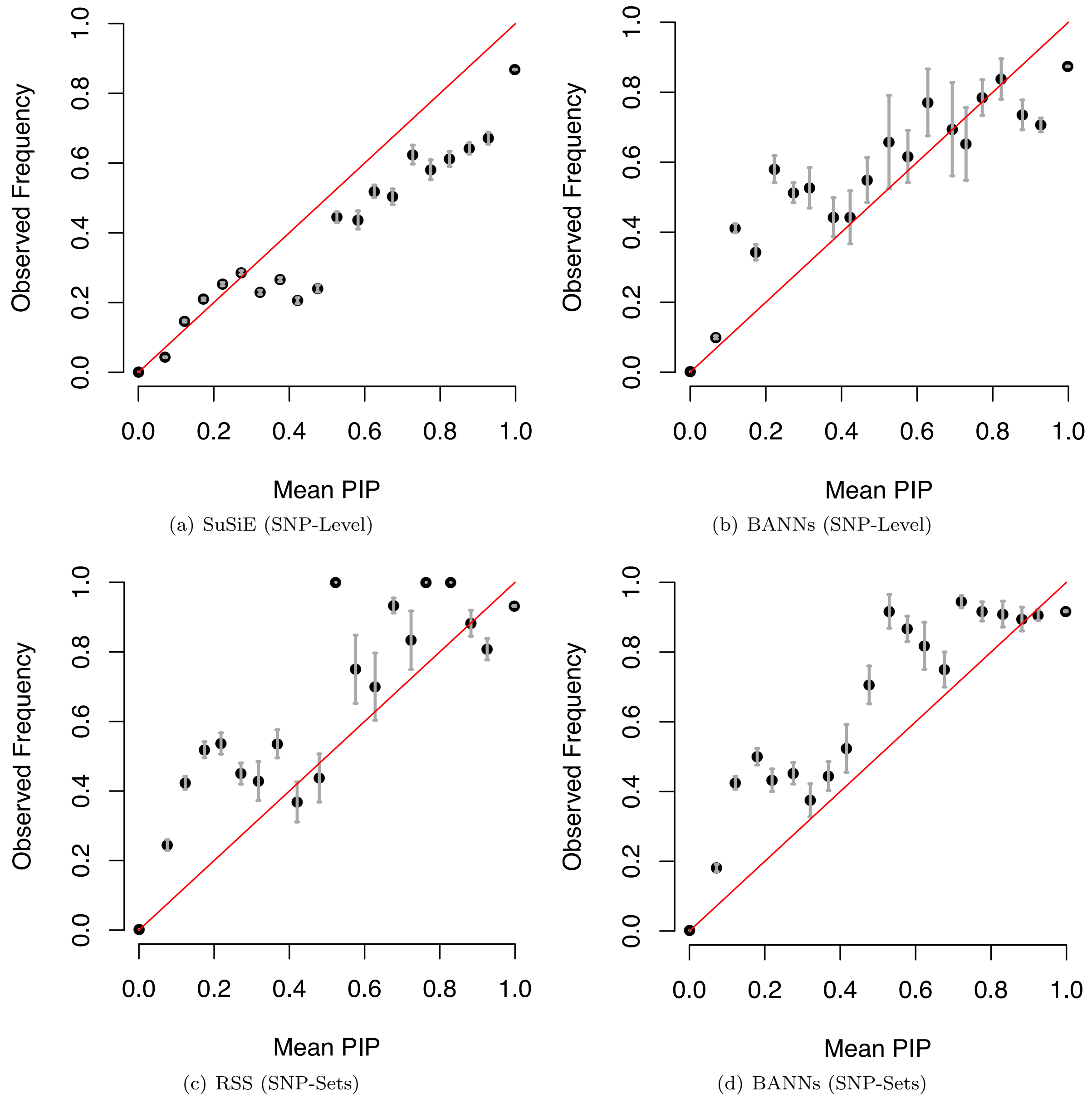

(a) SuSiE (SNP-Level)

(b) BANNs (SNP-Level)

(c) RSS (SNP-Sets)

(d) BANNs (SNP-Sets)

**Figure S24. Assessments of posterior inclusion probability (PIP) calibration for both SNP-level associations and enrichment of SNP-sets.** This experiment follows largely from previous work [14, 64]. Here, SNPs and SNP-sets across simulations are grouped into bins according to their reported PIPs (using 20 equally spaced bins, from 0 to 1).The plots show the average PIP for each bin against the proportion of causal SNPs or SNP-sets in that bin. A well calibrated method should produce points near the x-axis = y-axis line (i.e., the diagonal red lines). Gray error bars show ±2 standard errors. Panel **(a, b)** shows the comparison of BANNs SNP layer with SuSiE [64], and **(c, d)** shows the comparison of BANNs SNP-set layer with RSS [7]. While the inclusion probabilities are not perfectly calibrated for any of the methods, the empirical power and false discovery rate (FDR) above the "median probability criterion" (i.e., PIPs for SNPs and SNP-sets greater than 0.5) [51] are still reasonably well controlled (see Tables S1-S8). We hypothesize that these calibration results are due both to consequences of both variational inference and the level of polygenicity with which we simulated synthetic phenotypes.

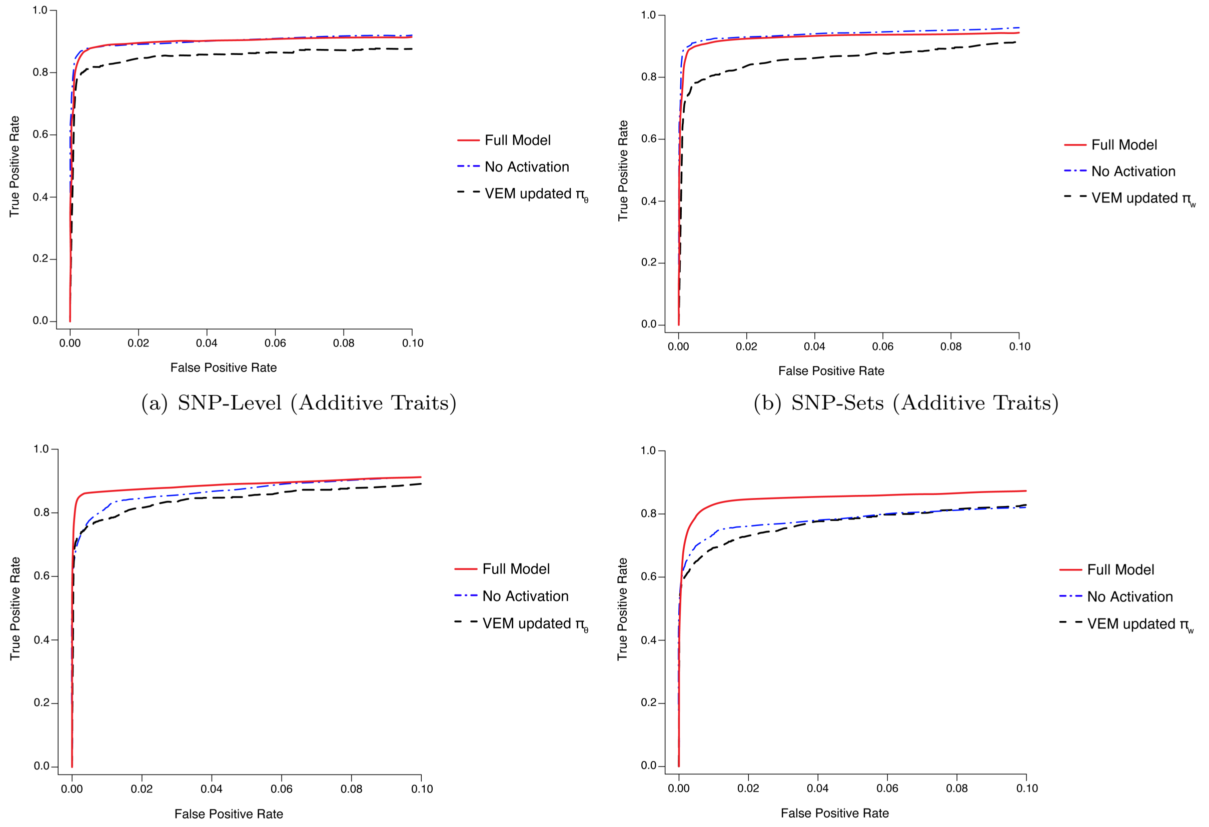

(a) SNP-Level (Additive Traits)

(b) SNP-Sets (Additive Traits)

(c) SNP-Level (Additive & Epistatic Traits)

(d) SNP-Sets (Additive & Epistatic Traits)

**Figure S25. Receiver operating characteristic (ROC) curves comparing the performance of the BANNs models with different modifications via an ablation test.** To investigate how choices in the model setup contribute to variable selection, we performed an "ablation analysis" where we modified parts of the BANNs framework independently and observed their direct effect on model performance (see Section 5). We considered two different modifications to our model: (1) removing the activation function and training a fully linear hierarchical model, and (2) removing the approximate Bayesian model averaging approach and updating the probabilities $\pi_\theta$ and $\pi_w$ as additional parameters in the variational EM algorithm. In the normal BANNs setup, we initialize $L$ different models with varying priors for inclusion probabilities specified over a grid $\{\pi_\theta^{(1)}, \ldots, \pi_\theta^{(L)}\} \in [1/J, 1]$ and $\{\pi_w^{(1)}, \ldots, \pi_w^{(L)}\} \in [1/G, 1]$, respectively. However, in the case of the latter ablation modification, we initialize $\pi_\theta = 1/J$ and $\pi_w = 1/G$ as an analogy to the "single causal variant" assumption frequently used in fine mapping [64]. Next, we update their values in the M-step of the algorithm according to the following analytic expressions: **(a, c)** $\pi_\theta/1-\pi_\theta = \sum_j \sum_k \alpha_{jk}/\sum_j \sum_k(1-\alpha_{jk})$, and **(b, d)** $\pi_w/1-\pi_w = \sum_g \alpha_g/\sum_g(1-\alpha_g)$. Results here are shown using simulations with the self-identified "white British" ancestry cohort from the UK Biobank on synthetic traits that have broad-sense heritability $H^2 = 0.6$ with sparse genetic architecture. Each plot combines results from 100 simulated replicates (see Section 9).

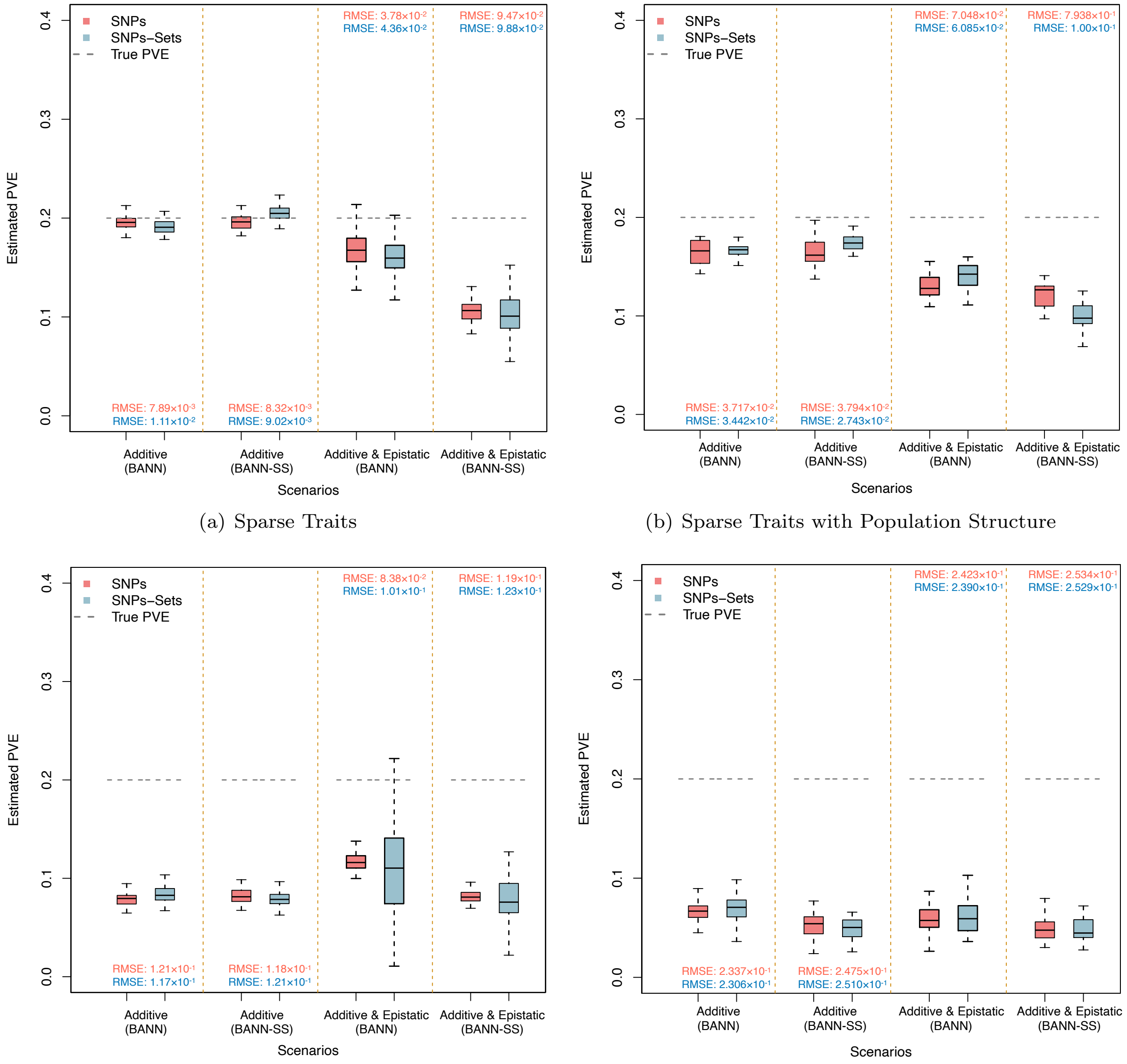

(a) Sparse Traits

(b) Sparse Traits with Population Structure

(c) Polygenic Traits

(d) Polygenic Traits with Population Structure

**Figure S26. Boxplots depicting the ability of the BANNs and BANN-SS models to estimate the phenotypic variation explained (PVE) by SNPs (pink) and SNP-sets (blue) for complex traits in simulations.** In this work, we define PVE as the total proportion of phenotypic variance that is explained by sparse genetic effects (both additive and non-additive) [15]. Here, quantitative traits are simulated to have broad-sense heritability of $H^2 = 0.2$ with different levels of contributions from additive effects and epistatic interactions. We consider two different trait architectures: **(a, b)** sparse where only 1% of SNP-sets are enriched for the trait; and **(c, d)** polygenic where 10% of SNP-sets are enriched. Panels **(a, c)** show heritability estimates on simulations with genetic data from individuals who self-identify as being of "white British" ancestry in the UK Biobank; while, panels **(b, d)** show heritability estimates on simulations with genetic data from individuals who more broadly identify as being of European ancestry. True heritability values are shown as the dashed grey horizontal lines. The root mean square error (RMSE) between the BANNs model estimates of the PVE and the true values are also provided.

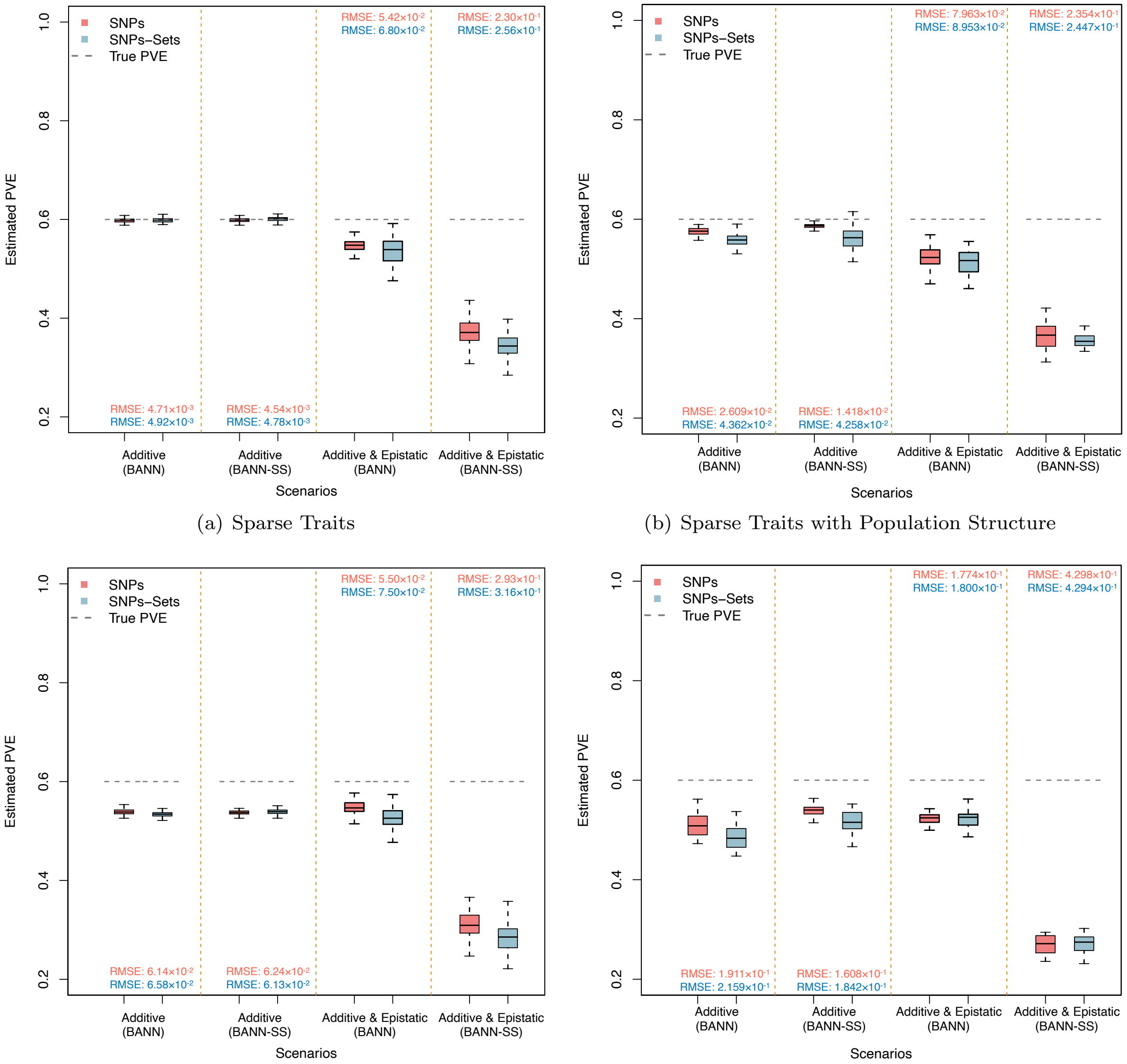

(a) Sparse Traits

(b) Sparse Traits with Population Structure

(c) Polygenic Traits

(d) Polygenic Traits with Population Structure

**Figure S27. Boxplots depicting the ability of the BANNs and BANN-SS models to estimate the phenotypic variation explained (PVE) by SNPs (pink) and SNP-sets (blue) for complex traits in simulations.** In this work, we define PVE as the total proportion of phenotypic variance that is explained by sparse genetic effects (both additive and non-additive) [15]. Here, quantitative traits are simulated to have broad-sense heritability of $H^2 = 0.6$ with different levels of contributions from additive effects and epistatic interactions. We consider two different trait architectures: **(a, b)** sparse where only 1% of SNP-sets are enriched for the trait; and **(c, d)** polygenic where 10% of SNP-sets are enriched. Panels **(a, c)** show heritability estimates on simulations with genetic data from individuals who self-identify as being of "white British" ancestry in the UK Biobank; while, panels **(b, d)** show heritability estimates on simulations with genetic data from individuals who more broadly identify as being of European ancestry. True heritability values are shown as the dashed grey horizontal lines. The root mean square error (RMSE) between the BANNs model estimates of the PVE and the true values are also provided.

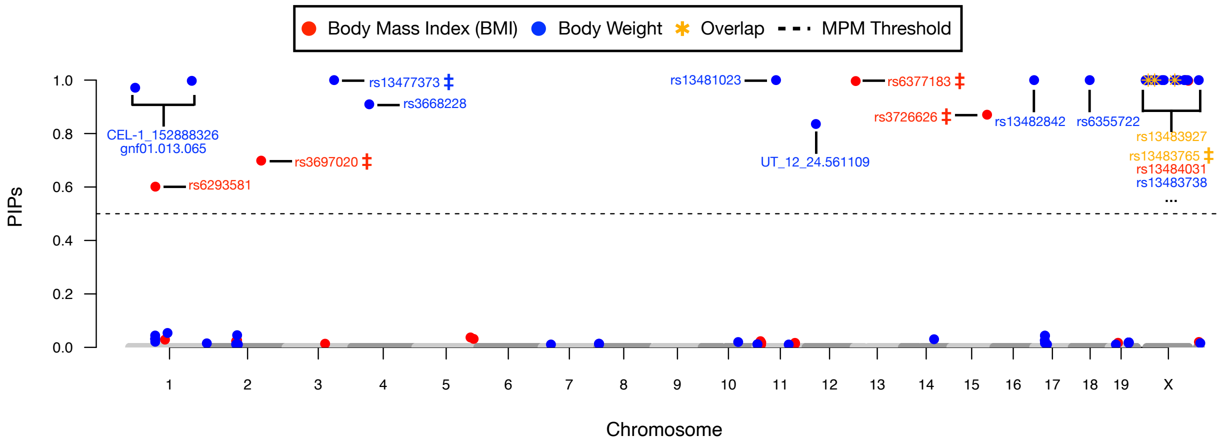

(a) Body Mass Index and Body Weight

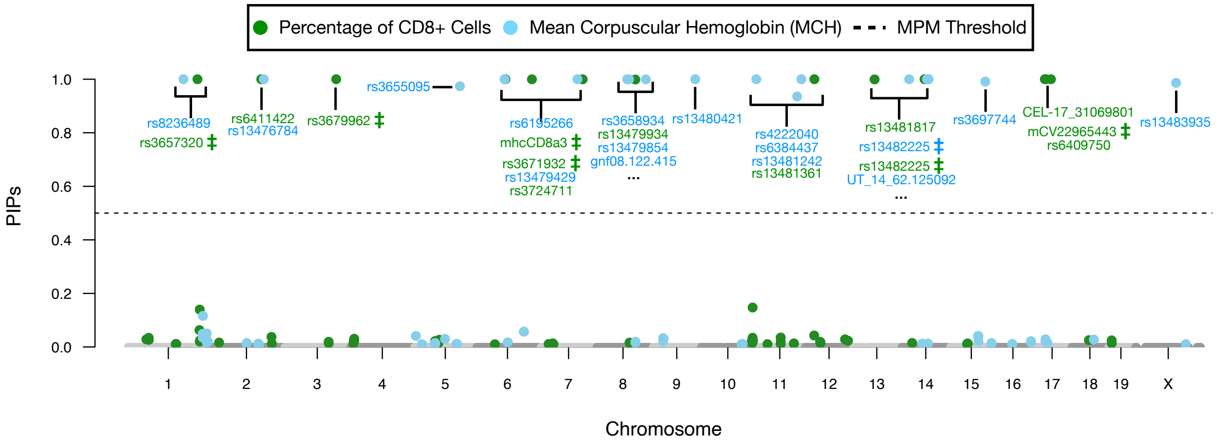

(b) % CD8+ cells and Mean Corpuscular Hemoglobin

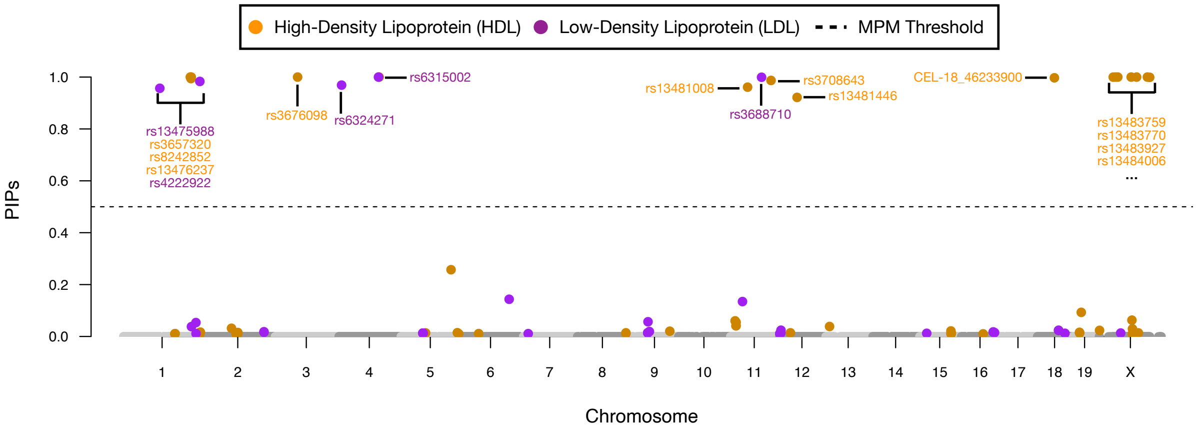

(c) High-Density and Low-Density Lipoprotein

**Figure S28. Manhattan plots of variant-level fine mapping results for six traits in heterogeneous stock of mice from the Wellcome Trust Centre for Human Genetics.** Traits are grouped based on their category and include: **(a)** body mass index (BMI) and body weight, **(b)** percentage of CD8+ cells and mean corpuscular hemoglobin (MCH), and **(c)** high-density and low-density lipoprotein (HDL and LDL, respectively) cholesterol. Posterior inclusion probabilities (PIP) for the input layer weights are derived from the BANNs model fit on individual-level data and are plotted for each SNP against their genomic positions. Chromosomes are shown in alternating colors for clarity. The black dashed line is marked at 0.5 and represents the "median probability model (MPM)" threshold [51]. Here, we only color code SNPs that had a PIP greater than 1% in either trait. SNPs with PIPs exceeding 1% in both traits are marked by a star and denoted as falling in the "overlap" category. BANNs estimated the following PVEs on the SNP and SNP-set levels for these traits, respectively: (*i*) 0.09 and 0.08 for BMI, (*ii*) 0.39 and 0.40 for body weight, (*iii*) 0.51 and 0.48 for percentage of CD8+ cells, (*iv*) 0.34 and 0.32 for MCH, (*v*) 0.34 and 0.28 for HDL, and (*vi*) 0.15 and 0.15 for LDL.

(a)

| | *p* value | *q* value | Odds.ratio | Combined score | # of sig. genes in dbGaP |
|---|---|---|---|---|---|
| Metabolic Syndrome X | 1.316e-04 | 4.541e-02 | 115.94 | 1036.01 | 2 |
| Lipoproteins, HDL | 9.365e-04 | 1.616e-01 | 43.72 | 304.85 | 2 |
| Cholesterol, HDL | 2.188e-03 | 2.516e-01 | 11.20 | 68.63 | 3 |
| Apolipoprotein A-I | 8.221e-03 | 7.091e-01 | 121.21 | 581.95 | 1 |
| Natriuretic Peptide, Brain | 1.342e-02 | 9.26e-01 | 74.07 | 319.33 | 1 |
| Triglycerides | 1.402e-02 | 8.06e-01 | 10.93 | 46.64 | 2 |
| Lipids | 6.332e-02 | 1.000 | 15.33 | 42.29 | 1 |
| Arteries | 6.473e-02 | 1.000 | 14.98 | 41.01 | 1 |
| Alcoholism | 6.755e-02 | 1.000 | 14.34 | 38.64 | 1 |
| Iron | 1.095e-01 | 1.000 | 8.66 | 19.15 | 1 |

(b)

| | *p* value | *q* value | Odds.ratio | Combined score | # of sig. genes in dbGaP |
|---|---|---|---|---|---|
| 1-Alkyl-2-acetylglycerophosphocholine Esterase | 2.098e-03 | 7.239e-01 | 476.19 | 2936.47 | 1 |
| Lipoproteins, LDL | 1.075e-02 | 1.000 | 92.59 | 419.68 | 1 |
| Alzheimer Disease | 2.376e-02 | 1.000 | 41.67 | 155.82 | 1 |
| Coronary Disease | 2.435e-02 | 1.000 | 40.65 | 151.02 | 1 |
| Coronary Artery Disease | 5.995e-02 | 1.000 | 16.26 | 45.76 | 1 |
| Myocardial Infarction | 6.677e-02 | 1.000 | 14.56 | 39.40 | 1 |
| Triglycerides | 7.101e-02 | 1.000 | 13.66 | 36.13 | 1 |
| Cholesterol | 7.776e-02 | 1.000 | 12.44 | 31.77 | 1 |
| Cholesterol, LDL | 8.781e-02 | 1.000 | 10.96 | 26.67 | 1 |
| Cholesterol, HDL | 1.024e-01 | 1.000 | 9.34 | 21.27 | 1 |

(c)

| | *p* value | *q* value | Odds.ratio | Combined score | # of sig. genes in GWAS Catalog |
|---|---|---|---|---|---|
| Apolipoprotein A1 levels | 3.409e-09 | 5.922e-06 | 800 | 15597.40 | 3 |
| High density lipoprotein cholesterol levels | 1.932e-07 | 1.678e-04 | 75.12 | 1161.28 | 4 |
| HDL cholesterol levels | 5.749e-07 | 3.329e-04 | 57.35 | 824.03 | 4 |
| HDL cholesterol | 6.234e-07 | 2.707e-04 | 28.01 | 400.23 | 5 |
| Triglyceride levels | 9.011e-07 | 3.130e-04 | 51.28 | 713.83 | 4 |
| Metabolite levels (lipoprotein measures) | 9.875e-07 | 2.859e-04 | 148.15 | 2048.61 | 3 |
| Lipid metabolism phenotypes | 2.611e-06 | 6.480e-04 | 108.11 | 1389.80 | 3 |
| Metabolic syndrome | 4.752e-06 | 1.032e-03 | 88.89 | 1089.51 | 3 |
| Mean diameter of HDL particles | 7.861e-06 | 1.517e-03 | 444.44 | 5223.81 | 2 |
| Total cholesterol levels | 9.924e-06 | 1.724e-03 | 28.07 | 323.38 | 4 |

(d)

| | *p* value | *q* value | Odds.ratio | Combined score | # of sig. genes in GWAS Catalog |
|---|---|---|---|---|---|
| Waist-to-hip circumference ratio (smoking years interaction) | 7.497e-07 | 1.302e-03 | 1333.33 | 18804.85 | 2 |
| Cerebrospinal fluid t-tau levels in mild cognitive impairment | 1.018e-05 | 8.841e-03 | 392.16 | 4507.90 | 2 |
| Cerebrospinal AB1-42 levels in mild cognitive impairment | 1.280e-05 | 7.409e-03 | 350.88 | 3953.12 | 2 |
| Cerebrospinal AB1-42 levels in Alzheimer's disease dementia | 1.571e-05 | 6.822e-03 | 317.46 | 3511.50 | 2 |
| Logical memory (immediate recall) | 1.892e-05 | 6.573e-03 | 289.86 | 3152.23 | 2 |
| Logical memory (delayed recall) | 2.064e-05 | 5.121e-03 | 277.78 | 2996.75 | 2 |
| Cerebrospinal fluid p-tau levels | 2.064e-05 | 5.975e-03 | 277.78 | 2996.75 | 2 |
| Cerebrospinal fluid p-tau levels in mild cognitive impairment | 2.43e-05 | 5.275e-03 | 256.41 | 2724.40 | 2 |
| Cerebrospinal fluid t-tau levels | 2.624e-05 | 5.064e-03 | 246.91 | 2604.53 | 2 |
| Bladder cancer | 2.825e-05 | 4.907e-03 | 238.10 | 2493.89 | 2 |

**Figure S29. Gene set enrichment analyses using the significant SNP-sets identified by BANNs for high-density and low-density lipoprotein (HDL and LDL, respectively) traits in the Framingham Heart Study [47].** Here, SNP-set annotations are based on gene boundaries defined by the NCBI's RefSeq database in the UCSC Genome Browser [49]. Unannotated SNPs located within the same genomic region were labeled as being within the "intergenic region" between two genes. Posterior inclusion probabilities (PIP) for the input and hidden layer weights are derived by fitting the BANNs model on individual-level data. A SNP-set is considered significant if it has a PIP$(g) \geq 0.5$ (i.e., the "median probability model" threshold [51]). We take these significant SNP-sets and conduct "gene set enrichment analysis" using Enrichr [71, 72] to identify the categories they overrepresent in **(a, b)** the database of Genotypes and Phenotypes (dbGaP) and **(c, d)** the GWAS Catalog (2019). Nearly all enriched categories are related with **(a, c)** HDL and **(b, d)** LDL, respectively. Note that in LDL, the BANNs framework identified the gene *APOB* as having a high PIP = 0.976. There have been hypotheses connecting LDL to cognitive traits [73,74], and *APOB* has been shown to be related to cerebrospinal fluid and memory [75–77]. Therefore, we argue that results in panel **(d)** are also relevant.

(a)

| | *p* value | *q* value | Odds.ratio | Combined score | # of sig. genes in GWAS Catalog |
|---|---|---|---|---|---|
| Blood Pressure Determination | 2.035e-03 | 4.681e-02 | 33.8 | 210 | 2 |
| Natriuretic Peptide, Brain | 1.874e-02 | 1.678e-01 | 58.7 | 234 | 1 |
| Brain | 3.512e-02 | 1.678e-01 | 30.2 | 101 | 1 |
| Body Weight Changes | 3.512e-02 | 1.678e-01 | 30.2 | 101 | 1 |
| Glomerular Filtration Rate | 4.622e-02 | 1.678e-01 | 22.7 | 69.6 | 1 |
| Erythrocytes | 5.323e-02 | 1.678e-01 | 19.5 | 57.3 | 1 |
| Hypertrophy, Left Ventricular | 5.721e-02 | 1.678e-01 | 18.1 | 51.8 | 1 |
| Exercise Test | 6.906e-02 | 1.678e-01 | 14.9 | 39.7 | 1 |
| Elbow | 7.297e-02 | 1.678e-01 | 14 | 36.7 | 1 |
| Amyotrophic Lateral Sclerosis | 7.688e-02 | 1.678e-01 | 13.3 | 34 | 1 |

(b)

| | *p* value | *q* value | Odds.ratio | Combined score | # of sig. genes in dbGaP |
|---|---|---|---|---|---|
| Cholesterol, HDL | 8.286e-03 | 4.142e-02 | 18.4 | 88.4 | 2 |
| Blood Flow Velocity | 1.035e-02 | 4.142e-02 | 114 | 521 | 1 |
| Insulin Resistance | 2.297e-02 | 4.571e-02 | 50 | 189 | 1 |
| Lipoproteins, HDL | 2.415e-02 | 4.571e-02 | 47.5 | 177 | 1 |
| Amyotrophic Lateral Sclerosis | 3e-02 | 4.571e-02 | 37.9 | 133 | 1 |
| Lipids | 3.428e-02 | 4.571e-02 | 33.1 | 112 | 1 |
| Macular Degeneration | 4.624e-02 | 5.285e-02 | 24.3 | 74.6 | 1 |
| Iron | 5.998e-02 | 5.998e-02 | 18.5 | 52.1 | 1 |

(c)

| | *p* value | *q* value | Odds.ratio | Combined score | # of sig. genes in GWAS Catalog |
|---|---|---|---|---|---|
| Seborrheic dermatitis | 3.495e-03 | 3.146e-02 | 370 | 2093 | 1 |
| QRS duration | 1.909e-02 | 6.966e-02 | 61.5 | 244 | 1 |
| PR interval | 2.322e-02 | 6.966e-02 | 50.3 | 189 | 1 |
| Creatinine levels | 3.346e-02 | 7.244e-02 | 34.5 | 117 | 1 |
| QT interval | 4.024e-02 | 7.244e-02 | 28.6 | 91.8 | 1 |
| Triglycerides | 6.266e-02 | 8.311e-02 | 18 | 50 | 1 |
| Total cholesterol levels | 6.464e-02 | 8.311e-02 | 17.5 | 47.8 | 1 |
| Diastolic blood pressure | 2.054e-01 | 2.085e-01 | 5 | 7.91 | 1 |
| Systolic blood pressure | 2.085e-01 | 2.085e-01 | 4.91 | 7.7 | 1 |

(d)

| | *p* value | *q* value | Odds.ratio | Combined score | # of sig. genes in GWAS Catalog |
|---|---|---|---|---|---|
| Low density lipoprotein cholesterol levels | 1.313e-04 | 5.199e-03 | 158 | 1415 | 2 |
| Ischemic stroke | 3.055e-04 | 5.199e-03 | 102 | 827 | 2 |
| Stroke | 3.627e-04 | 5.199e-03 | 93.5 | 741 | 2 |
| Metabolite levels | 8.363e-04 | 8.99e-03 | 60.8 | 431 | 2 |
| HDL cholesterol change in response to fenofibrate | 1.999e-03 | 1.719e-02 | 714 | 4437 | 1 |
| Cholesterol efflux capacity | 2.797e-03 | 2.005e-02 | 476 | 2798 | 1 |
| Metabolic syndrome (bivariate traits) | 3.595e-03 | 2.208e-02 | 357 | 2008 | 1 |
| Non-small cell lung cancer | 6.383e-03 | 3.429e-02 | 190 | 962 | 1 |
| C-reactive protein levels or HDL-cholesterol levels (pleiotropy) | 7.973e-03 | 3.429e-02 | 150 | 726 | 1 |
| Cardiovascular disease risk factors | 7.973e-03 | 3.429e-02 | 150 | 726 | 1 |

**Figure S30. Gene set enrichment analyses using the significant SNP-sets identified by BANNs for high-density and low-density lipoprotein (HDL and LDL, respectively) traits in the Framingham Heart Study [47].** Here, SNP-set annotations are based on gene boundaries defined by the NCBI's RefSeq database in the UCSC Genome Browser [49]. Unannotated SNPs located within the same genomic region were labeled as being within the "intergenic region" between two genes. In this analysis, each gene boundary annotation is modfied by adding SNPs within a ±500 kilobase (kb) buffer to account for possible regulatory elements. Posterior inclusion probabilities (PIP) for the input and hidden layer weights are derived by fitting the BANNs model on individual-level data. A SNP-set is considered significant if it has a PIP($g$) $\geq$ 0.5 (i.e., the "median probability model" threshold [51]). We take these significant SNP-sets and conduct "gene set enrichment analysis" using Enrichr [71, 72] to identify the categories they overrepresent in **(a, b)** the database of Genotypes and Phenotypes (dbGaP) and **(c, d)** the GWAS Catalog (2019). Nearly all enriched categories are related with **(a, c)** HDL and **(b, d)** LDL, respectively.

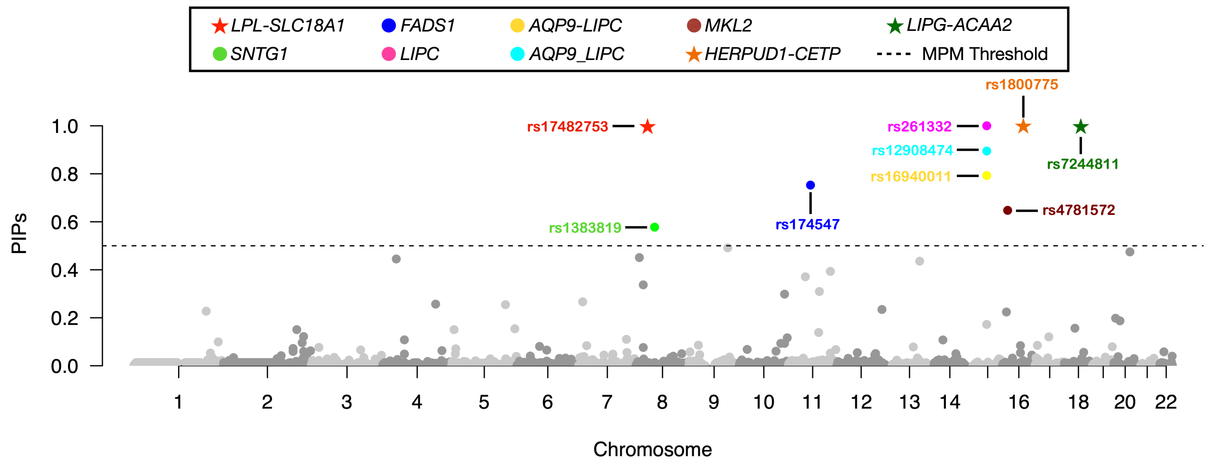

(a) High-Density Lipoprotein (HDL)

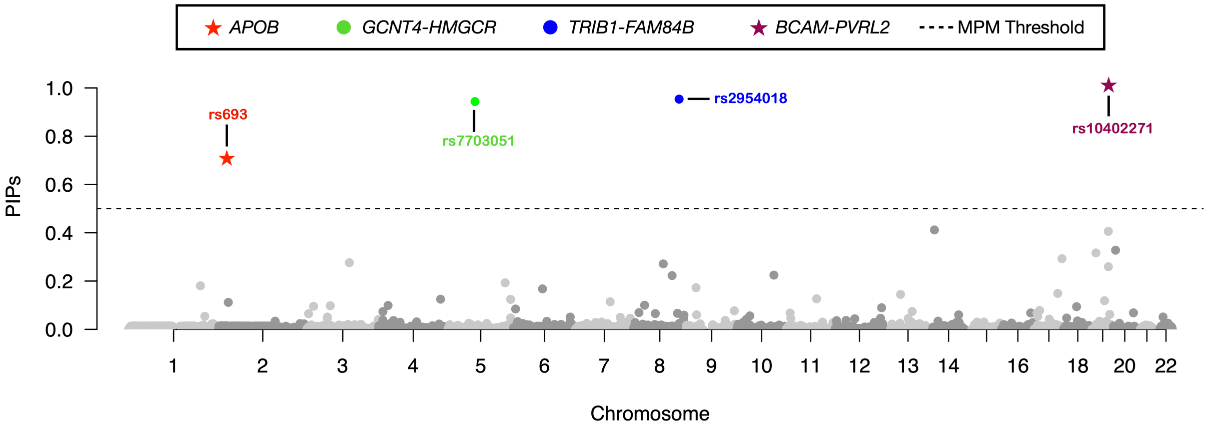

(b) Low-Density Lipoprotein (LDL)

**Figure S31. Manhattan plot of variant-level association mapping results for high-density and low-density lipoprotein (HDL and LDL, respectively) traits in ten thousand randomly sampled individuals of European ancestry from the UK Biobank [50].** Posterior inclusion probabilities (PIP) for the neural network weights are derived from the BANNs model fit on individual-level data and are plotted for each SNP against their genomic positions. Chromosomes are shown in alternating colors for clarity. The black dashed line is marked at 0.5 and represents the "median probability model" threshold [51]. SNPs with PIPs above that threshold are color coded based on their SNP-set annotation. Here, SNP-set annotations are based on gene boundaries defined by the NCBI's RefSeq database in the UCSC Genome Browser [49]. Unannotated SNPs located within the same genomic region were labeled as being within the "intergenic region" between two genes. These regions are labeled as *Gene1-Gene2* in the legend. Gene set enrichment analyses for these SNP-sets can be found in Fig. S31. Stars (★) denote SNPs and SNP-sets that replicate findings from our analyses of HDL and LDL in the Framingham Heart Study (See Fig. 4 in the main text).

(a)

| | *p* value | *q* value | Odds.ratio | Combined score | # of sig. genes in dbGaP |
|---|---|---|---|---|---|
| Lipoproteins, HDL | 2.812e-10 | 9.701e-08 | 67.84 | 1491.82 | 6 |
| Cholesterol, HDL | 1.216e-09 | 2.098e-07 | 17.39 | 356.90 | 9 |
| Metabolic Syndrome X | 4.760e-06 | 5.474e-04 | 89.96 | 1102.42 | 3 |
| Lipid Metabolism | 7.262e-05 | 6.264e-03 | 153.27 | 1460.57 | 2 |
| Electrocardiography | 2.677e-04 | 1.847e-02 | 23.78 | 195.61 | 3 |
| Triglycerides | 4.038e-04 | 2.322e-02 | 11.31 | 88.35 | 4 |
| Uric Acid | 1.310e-03 | 6.457e-02 | 37.28 | 247.44 | 2 |
| Iron | 1.412e-03 | 6.091e-02 | 13.43 | 88.16 | 3 |
| Phosphatidylcholines | 4.343e-03 | 1.665e-01 | 229.89 | 1250.34 | 1 |
| Epilepsies, Partial | 5.788e-03 | 1.997e-01 | 172.41 | 888.28 | 1 |

(b)

| | *p* value | *q* value | Odds.ratio | Combined score | # of sig. genes in dbGaP |
|---|---|---|---|---|---|
| Lipoproteins, LDL | 3.579e-06 | 1.235e-03 | 98.04 | 1229.45 | 3 |
| Cholesterol, HDL | 1.976e-04 | 3.409e-02 | 13.18 | 112.43 | 4 |
| Metabolic Syndrome X | 1.938e-07 | 3.329e-04 | 51.15 | 201.72 | 1 |
| Cholesterol | 2.131e-02 | 1.000 | 8.78 | 33.79 | 2 |
| Menopause | 2.604e-02 | 1.000 | 37.95 | 138.45 | 1 |
| Cholesterol, LDL | 2.694e-02 | 1.000 | 7.34 | 27.97 | 2 |
| Eosinophils | 3.840e-02 | 1.000 | 25.58 | 83.37 | 1 |
| Lipoproteins, HDL | 5.062e-02 | 1.000 | 19.29 | 57.54 | 1 |
| Alzheimer Disease | 6.589e-02 | 1.000 | 14.71 | 40.00 | 1 |
| Coronary Disease | 6.749e-02 | 1.000 | 14.35 | 38.68 | 1 |

(c)

| | *p* value | *q* value | Odds.ratio | Combined score | # of sig. genes in GWAS Catalog |
|---|---|---|---|---|---|
| High density lipoprotein cholesterol levels | 6.787e-014 | 1.179e-10 | 77.70 | 2356.18 | 8 |
| Metabolic syndrome | 2.690e-13 | 2.337e-10 | 107.28 | 3105.10 | 7 |
| HDL cholesterol levels | 6.635e-13 | 3.685e-10 | 59.33 | 1666.02 | 8 |
| HDL cholesterol | 7.740e-13 | 3.361e-10 | 28.98 | 808.09 | 10 |
| Metabolite levels (lipoprotein measures) | 1.550e-12 | 5.384e-10 | 153.26 | 4167.47 | 6 |
| Triglyceride levels | 1.593e-12 | 4.612e-10 | 53.05 | 1441.12 | 8 |
| Metabolite levels | 2.713e-12 | 6.733e-10 | 49.70 | 1323.78 | 8 |
| Lipid metabolism phenotypes | 1.205e-11 | 2.617e-09 | 111.84 | 2811.73 | 6 |
| C-reactive protein levels or HDL-cholesterol levels (pleiotropy) | 6.803e-11 | 1.313e-08 | 172.41 | 4036.39 | 5 |
| Cholesterol, total | 3.479e-10 | 6.043e-08 | 19.90 | 868.93 | 7 |

(d)

| | *p* value | *q* value | Odds.ratio | Combined score | # of sig. genes in GWAS Catalog |
|---|---|---|---|---|---|
| LDL cholesterol | 5.756e-09 | 1.000e-05 | 39.00 | 739.90 | 6 |
| Cerebrospinal AB1-42 levels in Alzheimer's disease dementia | 6.72e-07 | 5.836e-04 | 168.07 | 2388.75 | 3 |
| Metabolite levels (lipoprotein measures) | 1.473e-06 | 8.530e-04 | 130.72 | 1755.31 | 3 |
| Body mass index x age interaction | 4.942e-06 | 2.146e-03 | 88.24 | 1078.03 | 3 |
| Waist-to-hip circumference ratio (smoking years interaction) | 6.790e-06 | 2.359e-03 | 470.59 | 5600.05 | 2 |
| Total cholesterol levels | 1.705e-05 | 4.935e-03 | 24.77 | 271.94 | 4 |
| Response to ziprazidone in schizophrenia | 1.898e-05 | 4.710e-03 | 294.12 | 3197.64 | 2 |
| LDL cholesterol levels | 2.058e-05 | 4.469e-03 | 55.15 | 595.10 | 3 |
| Body mass index (age>50) | 2.581e-05 | 4.982e-03 | 51.15 | 540.39 | 3 |
| HDL cholesterol | 4.120e-05 | 7.156e-03 | 19.77 | 199.65 | 4 |

**Figure S32. Gene set enrichment analyses using the significant SNP-sets identified by BANNs for high-density and low-density lipoprotein (HDL and LDL, respectively) traits in ten thousand randomly sampled individuals of European ancestry from the UK Biobank [50].** Here, SNP-set annotations are based on gene boundaries defined by the NCBI's RefSeq database in the UCSC Genome Browser [49]. Unannotated SNPs located within the same genomic region were labeled as being within the "intergenic region" between two genes. Posterior inclusion probabilities (PIP) for the input and hidden layer weights are derived by fitting the BANNs model on individual-level data. A SNP-set is considered significant if it has a PIP$(g) \geq 0.5$ (i.e., the "median probability model" threshold [51]). We take these significant SNP-sets and conduct "gene set enrichment analysis" using Enrichr [71, 72] to identify the categories they overrepresent in **(a, b)** the database of Genotypes and Phenotypes (dbGaP) and **(c, d)** the GWAS Catalog (2019). Nearly all enriched categories are related with **(a, c)** HDL and **(b, d)** LDL, respectively. Note that in LDL, the BANNs framework again identifies the gene *APOB* as having a high PIP (replicating the finding in the Framingham Heart Study). There have been hypotheses connecting LDL to cognitive traits [73,74], and *APOB* has been shown to be related to cerebrospinal fluid and memory [75–77]. Therefore, we argue that results in panel **(b)** are also relevant (a similar argument can be made for Fig. S33).

(a)

| | *p* value | *q* value | Odds.ratio | Combined score | # of sig. genes in dbGaP |
|---|---|---|---|---|---|
| Neurotic Disorders | 3.474e-05 | 1.806e-02 | 344.22 | 3534.359 | 2 |
| Conduct Disorder | 2.753e-04 | 7.157e-03 | 98.30 | 805.849 | 2 |
| Respiratory Function Tests | 1.624e-03 | 2.790e-02 | 14.35 | 92.163 | 3 |
| Cholesterol, HDL | 2.146e-02 | 2.790e-02 | 8.23 | 50.581 | 4 |
| Erythrocytes | 2.938e-03 | 3.055e-02 | 27.5 | 160 | 2 |
| Myocardial Infarction | 5.257e-03 | 3.958e-02 | 9.36 | 49.1 | 3 |
| Epilepsies, Partial | 6.186e-03 | 3.958e-02 | 222 | 1128 | 1 |
| Triglycerides | 6.266e-03 | 3.958e-02 | 8.77 | 44.5 | 3 |
| Lipids | 8.014e-03 | 3.958e-02 | 16.1 | 77.9 | 2 |
| C-Reactive Protein | 8.192e-03 | 3.958e-02 | 15.9 | 76.6 | 2 |

(b)

| | *p* value | *q* value | Odds.ratio | Combined score | # of sig. genes in dbGaP |
|---|---|---|---|---|---|
| Leukocyte Count | 2.374e-02 | 1.197e-01 | 46.5 | 174 | 1 |
| Mental Competency | 2.492e-02 | 1.197e-01 | 44.2 | 163 | 1 |
| Body Weights and Measures | 5.098e-02 | 1.197e-01 | 21 | 62.6 | 1 |
| Arteries | 5.213e-02 | 1.197e-01 | 20.6 | 60.7 | 1 |
| Waist-Hip Ratio | 5.441e-02 | 1.197e-01 | 19.7 | 57.2 | 1 |
| Body Weight | 1.222e-01 | 2.24e-01 | 8.36 | 17.6 | 1 |
| Cholesterol | 1.495e-01 | 2.289e-01 | 6.71 | 12.8 | 1 |
| Cholesterol, LDL | 1.679e-01 | 2.289e-01 | 5.91 | 10.5 | 1 |
| Cholesterol, HDL | 1.944e-01 | 2.289e-01 | 5.01 | 8.21 | 1 |
| Body Height | 2.081e-01 | 2.289e-01 | 4.64 | 7.29 | 1 |

(c)

| | *p* value | *q* value | Odds.ratio | Combined score | # of sig. genes in GWAS Catalog |
|---|---|---|---|---|---|
| Immune response to measles-mumps-rubella vaccine | 2.710e-06 | 5.308e-04 | 143 | 1827 | 3 |
| Motion sickness | 1.622e-05 | 1.59e-03 | 73.7 | 813 | 3 |
| Cardiovascular risk factors (age interaction) | 8.314e-05 | 5.087e-03 | 197 | 1848 | 2 |
| Brain imaging | 1.038e-04 | 5.087e-03 | 172 | 1578 | 2 |
| Cutaneous squamous cell carcinoma | 2.091e-04 | 8.198e-03 | 115 | 972 | 2 |
| Waist-to-hip ratio adjusted for body mass index | 6.678e-04 | 2.181e-02 | 19.7 | 144 | 3 |
| HIV-1 susceptibility | 2.611e-06 | 6.480e-04 | 108.11 | 1389.80 | 3 |
| Rosacea symptom severity | 7.966e-04 | 2.231e-02 | 55 | 393 | 2 |
| Mercury levels | 1.334e-03 | 2.778e-02 | 15.4 | 102 | 3 |
| Emphysema imaging phenotypes | 1.34e-03 | 2.778e-02 | 41.7 | 276 | 2 |

(d)

| | *p* value | *q* value | Odds.ratio | Combined score | # of sig. genes in GWAS Catalog |
|---|---|---|---|---|---|
| Response to ziprazidone in schizophrenia | 4.791e-03 | 5.936e-02 | 259 | 1386 | 1 |
| Nonalcoholic fatty liver disease | 7.774e-03 | 5.936e-02 | 151 | 735 | 1 |
| Exploratory eye movement dysfunction in schizophrenia | 8.965e-03 | 5.936e-02 | 130 | 611 | 1 |
| Irritable bowel syndrome | 1.431e-02 | 5.936e-02 | 78.9 | 335 | 1 |
| Coronary artery calcified atherosclerotic plaque score in type2d | 1.49e-02 | 5.936e-02 | 75.6 | 318 | 1 |
| Colorectal or endometrial cancer | 1.727e-02 | 5.936e-02 | 64.8 | 263 | 1 |
| Response to amphetamines | 1.845e-02 | 5.936e-02 | 60.5 | 241 | 1 |
| Coronary artery calcified atherosclerotic plaque | 1.904e-02 | 5.936e-02 | 58.5 | 232 | 1 |
| Feeling miserable | 2.022e-02 | 5.936e-02 | 55 | 214 | 1 |
| Emphysema imaging phenotypes | 2.139e-02 | 5.936e-02 | 51.8 | 199 | 1 |

**Figure S33. Gene set enrichment analyses using the significant SNP-sets identified by BANNs for high-density and low-density lipoprotein (HDL and LDL, respectively) traits in ten thousand randomly sampled individuals of European ancestry from the UK Biobank [50].** Here, SNP-set annotations are based on gene boundaries defined by the NCBI's RefSeq database in the UCSC Genome Browser [49]. Unannotated SNPs located within the same genomic region were labeled as being within the "intergenic region" between two genes. In this analysis, each gene boundary annotation is modfied by adding SNPs within a $\pm 500$ kilobase (kb) buffer to account for possible regulatory elements. Posterior inclusion probabilities (PIP) for the input and hidden layer weights are derived by fitting the BANNs model on individual-level data. A SNP-set is considered significant if it has a PIP$(g) \geq 0.5$ (i.e., the "median probability model" threshold [51]). We take these significant SNP-sets and conduct "gene set enrichment analysis" using Enrichr [71,72] to identify the categories they overrepresent in **(a, b)** the database of Genotypes and Phenotypes (dbGaP) and **(c, d)** the GWAS Catalog (2019). Note that for panel **(a)**, BANNs did not find many enriched SNP-sets with PIPs meeting the "median probability model" threshold and so we used a lower SNP-set threshold (PIP $\geq 0.1$) to enable Enrichr to find associated dbGaP categories.

### 13 Supplementary Tables

| | | SNP-Level Approaches | | | | | |
|---|---|---|---|---|---|---|---|
| **Trait Type** | **Metric** | **BANN** | **BANN-SS** | **SuSiE (High)** | **SuSiE (Low)** | **CAVIAR** | **FINEMAP** |
| Sparse | **Power** | **0.614 (0.106)** | 0.609 (0.106) | 0.608 (0.087) | 0.424 (0.103) | 0.597 (0.167) | 0.529 (0.156) |
| | **FDR** | 0.211(0.092) | 0.210 (0.104) | **0.207 (0.052)** | 0.512 (0.088) | 0.331 (0.155) | 0.346 (0.048) |
| Polygenic | **Power** | 0.119 (0.039) | **0.120 (0.053)** | 0.103 (0.021) | 0.072 (0.037) | 0.098 (0.032) | 0.101 (0.044) |
| | **FDR** | 0.224 (0.098) | 0.227 (0.107) | **0.217 (0.034)** | 0.491 (0.051) | 0.244 (0.086) | 0.257 (0.103) |

| | | SNP-Set Level Approaches | | | | | |
|---|---|---|---|---|---|---|---|
| **Trait Type** | **Metric** | **BANN** | **BANN-SS** | **RSS** | **PEGASUS** | **SKAT** | **MAGMA** | **GSEA** |
| Sparse | **Power** | 0.608 (0.132) | 0.602 (0.161) | 0.609 (0.127) | 0.672 (0.118) | 0.543 (0.142) | **0.737 (0.123)** | 0.431 (0.138) |
| | **FDR** | **0.052 (0.107)** | 0.067 (0.098) | 0.088 (0.103) | 0.514 (0.137) | 0.466 (0.156) | 0.571 (0.126) | 0.579 (0.149) |
| Polygenic | **Power** | 0.078 (0.027) | 0.081 (0.041) | 0.073 (0.024) | 0.148 (0.032) | 0.112 (0.042) | **0.152 (0.039)** | 0.081 (0.031) |
| | **FDR** | 0.166 (0.151) | 0.163 (0.126) | **0.065 (0.112)** | 0.179 (0.098) | 0.181 (0.101) | 0.191 (0.018) | 0.221 (0.141) |

**Table S1. Comparing the empirical power and false discovery rates (FDR) of the BANNs framework against competing SNP and SNP-set mapping approaches in simulations.** Here, quantitative traits are simulated to have broad-sense heritability of $H^2 = 0.2$ with only contributions from additive effects set (i.e., $\rho = 1$). We consider two different trait architectures: sparse where only 1% of SNP-sets are enriched for the trait; and polygenic where 10% of SNP-sets are enriched. We set the number of causal SNPs with non-zero effects to be 1% and 10% of all SNPs located within the enriched SNP-sets, respectively. **(Top)** Competing SNP-level mapping approaches include: CAVIAR [63], SuSiE [64], and FINEMAP [65]. The software for SuSiE requires an input $\ell$ which fixes the maximum number of causal SNPs in the model. We display results when this input number is high ($\ell = 3000$) and when this input number is low ($\ell = 10$). **(Bottom)** Competing SNP-set mapping approaches include: RSS [7], PEGASUS [66], GBJ [67], SKAT [68], GSEA [69], and MAGMA [70]. Results for the BANN, BANN-SS, and other Bayesian methods are evaluated based on the "median probability criterion" (i.e., PIPs for SNPs and SNP-sets greater than 0.5) [51]. Results for the frequentist approaches are based on Bonferroni-corrected thresholds for multiple hypothesis testing ($P = 0.05/36518 = 1.37 \times 10^{-6}$ at the SNP-level and $P = 0.05/2816 = 1.78 \times 10^{-5}$ at the SNP-set level, respectively). All results are based on 100 replicates and standard deviations of the estimates across runs are given in the parentheses. Approaches with the greatest power are bolded in purple, while methods with the lowest FDR is bolded in blue.

| | | SNP-Level Approaches | | | | | |
|---|---|---|---|---|---|---|---|
| **Trait Type** | **Metric** | **BANN** | **BANN-SS** | **SuSiE (High)** | **SuSiE (Low)** | **CAVIAR** | **FINEMAP** |
| **Sparse** | **Power** | **0.851 (0.096)** | 0.812 (0.081) | 0.803 (0.034) | 0.631 (0.098) | 0.774 (0.159) | 0.722 (0.132) |
| | **FDR** | 0.201 (0.027) | 0.196 (0.029) | **0.185 (0.063)** | 0.522 (0.106) | 0.196 (0.093) | 0.248 (0.083) |
| **Polygenic** | **Power** | **0.374 (0.071)** | 0.369 (0.067) | 0.296 (0.074) | 0.198 (0.061) | 0.319 (0.106) | 0.332 (0.044) |
| | **FDR** | 0.212 (0.018) | **0.198 (0.026)** | 0.208 (0.022) | 0.414 (0.042) | 0.205 (0.031) | 0.307 (0.109) |

| | | SNP-Set Level Approaches | | | | | | |
|---|---|---|---|---|---|---|---|---|
| **Trait Type** | **Metric** | **BANN** | **BANN-SS** | **RSS** | **PEGASUS** | **SKAT** | **MAGMA** | **GSEA** |
| **Sparse** | **Power** | **0.823 (0.108)** | 0.821 (0.112) | 0.783 (0.105) | 0.815 (0.102) | 0.757 (0.114) | 0.821 (0.097) | 0.627 (0.123) |
| | **FDR** | 0.121 (0.087) | 0.127 (0.091 | **0.099 (0.096)** | 0.713 (0.081) | 0.692 (0.089) | 0.742 (0.061) | 0.581 (0.051) |
| **Polygenic** | **Power** | 0.276 (0.083) | 0.279 (0.097) | 0.272 (0.029) | 0.416 (0.076) | 0.331 (0.038) | **0.451 (0.049)** | 0.241 (0.022) |
| | **FDR** | 0.171 (0.034) | 0.166 (0.041) | **0.069 (0.040)** | 0.309 (0.087) | 0.282 (0.079) | 0.383 (0.083) | 0.322 (0.018) |

**Table S2. Comparing the empirical power and false discovery rates (FDR) of the BANNs framework against competing SNP and SNP-set mapping approaches in simulations.** Here, quantitative traits are simulated to have broad-sense heritability of $H^2 = 0.6$ with only contributions from additive effects set (i.e., $\rho = 1$). We consider two different trait architectures: sparse where only 1% of SNP-sets are enriched for the trait; and polygenic where 10% of SNP-sets are enriched. We set the number of causal SNPs with non-zero effects to be 1% and 10% of all SNPs located within the enriched SNP-sets, respectively. **(Top)** Competing SNP-level mapping approaches include: CAVIAR [63], SuSiE [64], and FINEMAP [65]. The software for SuSiE requires an input $\ell$ which fixes the maximum number of causal SNPs in the model. We display results when this input number is high ($\ell = 3000$) and when this input number is low ($\ell = 10$). **(Bottom)** Competing SNP-set mapping approaches include: RSS [7], PEGASUS [66], GBJ [67], SKAT [68], GSEA [69], and MAGMA [70]. Results for the BANN, BANN-SS, and other Bayesian methods are evaluated based on the "median probability criterion" (i.e., PIPs for SNPs and SNP-sets greater than 0.5) [51]. Results for the frequentist approaches are based on Bonferroni-corrected thresholds for multiple hypothesis testing ($P = 0.05/36518 = 1.37 \times 10^{-6}$ at the SNP-level and $P = 0.05/2816 = 1.78 \times 10^{-5}$ at the SNP-set level, respectively). All results are based on 100 replicates and standard deviations of the estimates across runs are given in the parentheses. Approaches with the greatest power are bolded in purple, while methods with the lowest FDR is bolded in blue.

| | | SNP-Level Approaches | | | | | |
|---|---|---|---|---|---|---|---|
| **Trait Type** | **Metric** | **BANN** | **BANN-SS** | **SuSiE (High)** | **SuSiE (Low)** | **CAVIAR** | **FINEMAP** |
| Sparse | **Power** | **0.621 (0.132)** | 0.613 (0.100) | 0.602 (0.084) | 0.591 (0.122) | 0.585 (0.094) | 0.569 (0.117) |
| | **FDR** | 0.219 (0.094) | 0.214 (0.104) | **0.173 (0.099)** | 0.191 (0.056) | 0.184 (0.080) | 0.221 (0.103) |
| Polygenic | **Power** | 0.120 (0.103) | 0.117 (0.059) | **0.127 (0.088)** | 0.081 (0.070) | 0.109 (0.041) | 0.091 (0.022) |
| | **FDR** | **0.194 (0.090)** | 0.223 (0.103) | 0.206 (0.061) | 0.274 (0.092) | 0.256 (0.106) | 0.321 (0.092) |

| | | SNP-Set Level Approaches | | | | | | |
|---|---|---|---|---|---|---|---|---|
| **Trait Type** | **Metric** | **BANN** | **BANN-SS** | **RSS** | **PEGASUS** | **SKAT** | **MAGMA** | **GSEA** |
| Sparse | **Power** | **0.616 (0.112)** | 0.602 (0.126) | 0.610 (0.101) | 0.573 (0.156) | 0.536 (0.124) | 0.549 (0.119) | 0.498 (0.090) |
| | **FDR** | 0.129 (0.093) | 0.182 (0.081) | **0.097 (0.107)** | 0.521 (0.133) | 0.512 (0.231) | 0.617 (0.216) | 0.633 (0.105) |
| Polygenic | **Power** | **0.102 (0.108)** | 0.096 (0.104) | **0.102 (0.117)** | 0.099 (0.018) | 0.063 (0.102) | 0.078 (0.053) | 0.057 (0.022) |
| | **FDR** | 0.228 (0.084) | **0.186 (0.092)** | 0.231 (0.097) | 0.443 (0.201) | 0.502 (0.310) | 0.491 (0.199) | 0.388 (0.178) |

**Table S3. Comparing the empirical power and false discovery rates (FDR) of the BANNs framework against competing SNP and SNP-set mapping approaches in simulations with population structure (European cohort).** Here, quantitative traits are simulated to have broad-sense heritability of $H^2 = 0.2$ with only contributions from additive effects set (i.e., $\rho = 1$). We consider two different trait architectures: sparse where only 1% of SNP-sets are enriched for the trait; and polygenic where 10% of SNP-sets are enriched. We set the number of causal SNPs with non-zero effects to be 1% and 10% of all SNPs located within the enriched SNP-sets, respectively. **(Top)** Competing SNP-level mapping approaches include: CAVIAR [63], SuSiE [64], and FINEMAP [65]. The software for SuSiE requires an input $\ell$ which fixes the maximum number of causal SNPs in the model. We display results when this input number is high ($\ell = 3000$) and when this input number is low ($\ell = 10$). **(Bottom)** Competing SNP-set mapping approaches include: RSS [7], PEGASUS [66], GBJ [67], SKAT [68], GSEA [69], and MAGMA [70]. Results for the BANN, BANN-SS, and other Bayesian methods are evaluated based on the "median probability criterion" (i.e., PIPs for SNPs and SNP-sets greater than 0.5) [51]. Results for the frequentist approaches are based on Bonferroni-corrected thresholds for multiple hypothesis testing ($P = 0.05/36518 = 1.37 \times 10^{-6}$ at the SNP-level and $P = 0.05/2816 = 1.78 \times 10^{-5}$ at the SNP-set level, respectively). All results are based on 100 replicates and standard deviations of the estimates across runs are given in the parentheses. Approaches with the greatest power are bolded in purple, while methods with the lowest FDR is bolded in blue.

| | | SNP-Level Approaches | | | | | |
|---|---|---|---|---|---|---|---|
| **Trait Type** | **Metric** | **BANN** | **BANN-SS** | **SuSiE (High)** | **SuSiE (Low)** | **CAVIAR** | **FINEMAP** |
| **Sparse** | **Power** | **0.822 (0.113)** | 0.818 (0.092) | 0.811 (0.071) | 0.623 (0.103) | 0.787(0.109) | 0.762 (0.121) |
| | **FDR** | **0.118 (0.074)** | 0.201 (0.104) | 0.123 (0.099) | 0.461 (0.056) | 0.227 (0.080) | 0.203 (0.103) |
| **Polygenic** | **Power** | **0.335 (0.042)** | 0.317 (0.059) | 0.326 (0.088) | 0.122 (0.070) | 0.301 (0.041) | 0.298 (0.031) |
| | **FDR** | 0.262 (0.090) | 0.341 (0.103) | 0.228(0.061) | 0.388 (0.092) | **0.211 (0.106)** | 0.294 (0.092) |

| | | SNP-Set Level Approaches | | | | | | |
|---|---|---|---|---|---|---|---|---|
| **Trait Type** | **Metric** | **BANN** | **BANN-SS** | **RSS** | **PEGASUS** | **SKAT** | **MAGMA** | **GSEA** |
| **Sparse** | **Power** | 0.781 (0.109) | 0.749 (0.117) | 0.743 (0.105) | **0.814 (0.112)** | 0.773 (0.127) | 0.802 (0.091) | 0.699 (0.118) |
| | **FDR** | **0.121 (0.104)** | 0.124 (0.098) | 0.312 (0.099) | 0.827 (0.056) | 0.805 (0.065) | 0.833 (0.051) | 0.841 (0.077) |
| **Polygenic** | **Power** | 0.294 (0.042) | 0.281 (0.053) | 0.301 (0.034) | 0.419 (0.047) | 0.341 (0.038) | **0.465 (0.038)** | 0.318 (0.078) |
| | **FDR** | 0.166 (0.054) | **0.159 (0.071)** | 0.178 (0.062) | 0.452 (0.089) | 0.418 (0.095) | 0.471 (0.079) | 0.516 (0.214) |

**Table S4. Comparing the empirical power and false discovery rates (FDR) of the BANNs framework against competing SNP and SNP-set mapping approaches in simulations with population structure (European cohort).** Here, quantitative traits are simulated to have broad-sense heritability of $H^2 = 0.6$ with only contributions from additive effects set (i.e., $\rho = 1$). We consider two different trait architectures: sparse where only 1% of SNP-sets are enriched for the trait; and polygenic where 10% of SNP-sets are enriched. We set the number of causal SNPs with non-zero effects to be 1% and 10% of all SNPs located within the enriched SNP-sets, respectively. **(Top)** Competing SNP-level mapping approaches include: CAVIAR [63], SuSiE [64], and FINEMAP [65]. The software for SuSiE requires an input $\ell$ which fixes the maximum number of causal SNPs in the model. We display results when this input number is high ($\ell = 3000$) and when this input number is low ($\ell = 10$). **(Bottom)** Competing SNP-set mapping approaches include: RSS [7], PEGASUS [66], GBJ [67], SKAT [68], GSEA [69], and MAGMA [70]. Results for the BANN, BANN-SS, and other Bayesian methods are evaluated based on the "median probability criterion" (i.e., PIPs for SNPs and SNP-sets greater than 0.5) [51]. Results for the frequentist approaches are based on Bonferroni-corrected thresholds for multiple hypothesis testing ($P = 0.05/36518 = 1.37 \times 10^{-6}$ at the SNP-level and $P = 0.05/2816 = 1.78 \times 10^{-5}$ at the SNP-set level, respectively). All results are based on 100 replicates and standard deviations of the estimates across runs are given in the parentheses. Approaches with the greatest power are bolded in purple, while methods with the lowest FDR is bolded in blue.

| | | SNP-Level Approaches | | | | | |
|---|---|---|---|---|---|---|---|
| **Trait Type** | **Metric** | **BANN** | **BANN-SS** | **SuSiE (High)** | **SuSiE (Low)** | **CAVIAR** | **FINEMAP** |
| **Sparse** | **Power** | **0.522 (0.122)** | 0.406 (0.094) | 0.402 (0.082) | 0.312 (0.077) | 0.309 (0.093) | 0.307 (0.087) |
| | **FDR** | 0.296 (0.113) | 0.311 (0.102) | **0.187 (0.061)** | 0.421 (0.089) | 0.207 (0.099) | 0.214 (0.068) |
| **Polygenic** | **Power** | **0.104 (0.058)** | 0.088 (0.033) | 0.094 (0.042) | 0.053 (0.032) | 0.081 (0.027) | 0.092 (0.031) |
| | **FDR** | 0.217 (0.061) | **0.186 (0.042)** | 0.193 (0.063) | 0.398 (0.092) | 0.194 (0.054) | 0.203 (0.059) |

| | | SNP-Set Level Approaches | | | | | | |
|---|---|---|---|---|---|---|---|---|
| **Trait Type** | **Metric** | **BANN** | **BANN-SS** | **RSS** | **PEGASUS** | **SKAT** | **MAGMA** | **GSEA** |
| **Sparse** | **Power** | **0.528 (0.113)** | 0.421 (0.098) | 0.476 (0.092) | 0.504 (0.109) | 0.426 (0.113) | 0.443 (0.128) | 0.378 (0.099) |
| | **FDR** | **0.073 (0.024)** | 0.095 (0.032) | 0.079 (0.023) | 0.201 (0.098) | 0.231 (0.106) | 0.312 (0.129) | 0.347 (0.127) |
| **Polygenic** | **Power** | 0.069 (0.029) | 0.051 (0.041) | 0.057 (0.024) | 0.056 (0.032) | 0.112 (0.042) | **0.152 (0.039)** | 0.081 (0.031) |
| | **FDR** | 0.166 (0.151) | 0.163 (0.126) | **0.065 (0.112)** | 0.179 (0.098) | 0.181 (0.101) | 0.191 (0.018) | 0.221 (0.141) |

**Table S5. Comparing the empirical power and false discovery rates (FDR) of the BANNs framework against competing SNP and SNP-set mapping approaches in simulations.** Here, quantitative traits are simulated to have broad-sense heritability of $H^2 = 0.2$ with contributions from both additive and epistatic effects set (i.e., $\rho = 0.5$). We consider two different trait architectures: sparse where only 1% of SNP-sets are enriched for the trait; and polygenic where 10% of SNP-sets are enriched. We set the number of causal SNPs with non-zero effects to be 1% and 10% of all SNPs located within the enriched SNP-sets, respectively. **(Top)** Competing SNP-level mapping approaches include: CAVIAR [63], SuSiE [64], and FINEMAP [65]. The software for SuSiE requires an input $\ell$ which fixes the maximum number of causal SNPs in the model. We display results when this input number is high ($\ell = 3000$) and when this input number is low ($\ell = 10$). **(Bottom)** Competing SNP-set mapping approaches include: RSS [7], PEGASUS [66], GBJ [67], SKAT [68], GSEA [69], and MAGMA [70]. Results for the BANN, BANN-SS, and other Bayesian methods are evaluated based on the "median probability criterion" (i.e., PIPs for SNPs and SNP-sets greater than 0.5) [51]. Results for the frequentist approaches are based on Bonferroni-corrected thresholds for multiple hypothesis testing ($P = 0.05/36518 = 1.37 \times 10^{-6}$ at the SNP-level and $P = 0.05/2816 = 1.78 \times 10^{-5}$ at the SNP-set level, respectively). All results are based on 100 replicates and standard deviations of the estimates across runs are given in the parentheses. Approaches with the greatest power are bolded in purple, while methods with the lowest FDR is bolded in blue.

| | | SNP-Level Approaches | | | | | |
|---|---|---|---|---|---|---|---|
| **Trait Type** | **Metric** | **BANN** | **BANN-SS** | **SuSiE (High)** | **SuSiE (Low)** | **CAVIAR** | **FINEMAP** |
| **Sparse** | **Power** | **0.851 (0.096)** | 0.812 (0.081) | 0.803 (0.034) | 0.631 (0.098) | 0.774 (0.159) | 0.722 (0.132) |
| | **FDR** | 0.201 (0.027) | 0.196 (0.029) | **0.185 (0.063)** | 0.522 (0.106) | 0.196 (0.093) | 0.248 (0.083) |
| **Polygenic** | **Power** | **0.374 (0.071)** | 0.369 (0.067) | 0.296 (0.074) | 0.198 (0.061) | 0.319 (0.106) | 0.332 (0.044) |
| | **FDR** | 0.212 (0.018) | **0.198 (0.026)** | 0.208 (0.022) | 0.414 (0.042) | 0.205 (0.031) | 0.307 (0.109) |

| | | SNP-Set Level Approaches | | | | | | |
|---|---|---|---|---|---|---|---|---|
| **Trait Type** | **Metric** | **BANN** | **BANN-SS** | **RSS** | **PEGASUS** | **SKAT** | **MAGMA** | **GSEA** |
| **Sparse** | **Power** | **0.823 (0.108)** | 0.821 (0.112) | 0.783 (0.105) | 0.815 (0.102) | 0.757 (0.114) | 0.821 (0.097) | 0.627 (0.123) |
| | **FDR** | 0.121 (0.087) | 0.127 (0.091) | **0.099 (0.096)** | 0.713 (0.081) | 0.692 (0.089) | 0.742 (0.061) | 0.581 (0.051) |
| **Polygenic** | **Power** | 0.276 (0.083) | 0.279 (0.097) | 0.272 (0.029) | 0.416 (0.076) | 0.331 (0.038) | **0.451 (0.049)** | 0.241 (0.022) |
| | **FDR** | 0.171 (0.034) | 0.166 (0.041) | **0.069 (0.040)** | 0.309 (0.087) | 0.282 (0.079) | 0.383 (0.083) | 0.322 (0.018) |

**Table S6. Comparing the empirical power and false discovery rates (FDR) of the BANNs framework against competing SNP and SNP-set mapping approaches in simulations.** Here, quantitative traits are simulated to have broad-sense heritability of $H^2 = 0.6$ with contributions from both additive and epistatic effects set (i.e., $\rho = 0.5$). We consider two different trait architectures: sparse where only 1% of SNP-sets are enriched for the trait; and polygenic where 10% of SNP-sets are enriched. We set the number of causal SNPs with non-zero effects to be 1% and 10% of all SNPs located within the enriched SNP-sets, respectively. **(Top)** Competing SNP-level mapping approaches include: CAVIAR [63], SuSiE [64], and FINEMAP [65]. The software for SuSiE requires an input $\ell$ which fixes the maximum number of causal SNPs in the model. We display results when this input number is high ($\ell = 3000$) and when this input number is low ($\ell = 10$). **(Bottom)** Competing SNP-set mapping approaches include: RSS [7], PEGASUS [66], GBJ [67], SKAT [68], GSEA [69], and MAGMA [70]. Results for the BANN, BANN-SS, and other Bayesian methods are evaluated based on the "median probability criterion" (i.e., PIPs for SNPs and SNP-sets greater than 0.5) [51]. Results for the frequentist approaches are based on Bonferroni-corrected thresholds for multiple hypothesis testing ($P = 0.05/36518 = 1.37 \times 10^{-6}$ at the SNP-level and $P = 0.05/2816 = 1.78 \times 10^{-5}$ at the SNP-set level, respectively). All results are based on 100 replicates and standard deviations of the estimates across runs are given in the parentheses. Approaches with the greatest power are bolded in purple, while methods with the lowest FDR is bolded in blue.

| | | SNP-Level Approaches | | | | | |
|---|---|---|---|---|---|---|---|
| **Trait Type** | **Metric** | **BANN** | **BANN-SS** | **SuSiE (High)** | **SuSiE (Low)** | **CAVIAR** | **FINEMAP** |
| **Sparse** | **Power** | **0.582 (0.114)** | 0.491 (0.092) | 0.526 (0.071) | 0.453 (0.103) | 0.474(0.109) | 0.491 (0.121) |
| | **FDR** | **0.118 (0.094)** | 0.201 (0.104) | 0.123 (0.099) | 0.461 (0.056) | 0.227 (0.080) | 0.203 (0.103) |
| **Polygenic** | **Power** | **0.112 (0.103)** | 0.087 (0.059) | 0.094 (0.088) | 0.042 (0.070) | 0.072 (0.041) | 0.064 (0.022) |
| | **FDR** | 0.262 (0.090) | 0.341 (0.103) | 0.228(0.061) | 0.388 (0.092) | **0.211 (0.106)** | 0.294 (0.092) |

| | | SNP-Set Level Approaches | | | | | | |
|---|---|---|---|---|---|---|---|---|
| **Trait Type** | **Metric** | **BANN** | **BANN-SS** | **RSS** | **PEGASUS** | **SKAT** | **MAGMA** | **GSEA** |
| **Sparse** | **Power** | **0.544 (0.076)** | 0.506 (0.053) | 0.509 (0.101) | 0.498 (0.122) | 0.388 (0.124) | 0.431 (0.119) | 0.320 (0.090) |
| | **FDR** | 0.132 (0.111) | 0.196 (0.097) | **0.130 (0.104)** | 0.564 (0.204) | 0.551 (0.120) | 0.427 (0.118) | 0.547 (0.221) |
| **Polygenic** | **Power** | **0.076 (0.123)** | 0.047 (0.104) | 0.051( 0.087) | 0.048 (0.018) | 0.036 (0.102) | 0.043 (0.053) | 0.022 (0.022) |
| | **FDR** | 0.241 (0.052) | 0.238 (0.092) | **0.212 (0.097)** | 0.397 (0.201) | 0.484 (0.310) | 0.563 (0.199) | 0.488 (0.178) |

**Table S7. Comparing the empirical power and false discovery rates (FDR) of the BANNs framework against competing SNP and SNP-set mapping approaches in simulations with population structure (European cohort).** Here, quantitative traits are simulated to have broad-sense heritability of $H^2 = 0.2$ with contributions from both additive and epistatic effects set (i.e., $\rho = 0.5$). We consider two different trait architectures: sparse where only 1% of SNP-sets are enriched for the trait; and polygenic where 10% of SNP-sets are enriched. We set the number of causal SNPs with non-zero effects to be 1% and 10% of all SNPs located within the enriched SNP-sets, respectively. **(Top)** Competing SNP-level mapping approaches include: CAVIAR [63], SuSiE [64], and FINEMAP [65]. The software for SuSiE requires an input $\ell$ which fixes the maximum number of causal SNPs in the model. We display results when this input number is high ($\ell = 3000$) and when this input number is low ($\ell = 10$). **(Bottom)** Competing SNP-set mapping approaches include: RSS [7], PEGASUS [66], GBJ [67], SKAT [68], GSEA [69], and MAGMA [70]. Results for the BANN, BANN-SS, and other Bayesian methods are evaluated based on the "median probability criterion" (i.e., PIPs for SNPs and SNP-sets greater than 0.5) [51]. Results for the frequentist approaches are based on Bonferroni-corrected thresholds for multiple hypothesis testing ($P = 0.05/36518 = 1.37 \times 10^{-6}$ at the SNP-level and $P = 0.05/2816 = 1.78 \times 10^{-5}$ at the SNP-set level, respectively). All results are based on 100 replicates and standard deviations of the estimates across runs are given in the parentheses. Approaches with the greatest power are bolded in purple, while methods with the lowest FDR is bolded in blue.

| | | SNP-Level Approaches | | | | | |
|---|---|---|---|---|---|---|---|
| **Trait Type** | **Metric** | **BANN** | **BANN-SS** | **SuSiE (High)** | **SuSiE (Low)** | **CAVIAR** | **FINEMAP** |
| **Sparse** | **Power** | **0.817 (0.126)** | 0.798 (0.117) | 0.792 (0.092) | 0.563 (0.104) | 0.752 (0.134) | 0.726 (0.128) |
| | **FDR** | **0.182 (0.038)** | 0.191 (0.045) | 0.346 (0.057) | 0.467 (0.075) | 0.237 (0.076) | 0.282 (0.084) |
| **Polygenic** | **Power** | **0.348 (0.109)** | 0.319 (0.094) | 0.305 (0.081) | 0.211 (0.039) | 0.327 (0.093) | 0.338 (0.053) |
| | **FDR** | 0.239 (0.047) | **0.221 (0.038)** | 0.224 (0.042) | 0.385 (0.035) | 0.309 (0.041) | 0.325 (0.091) |

| | | SNP-Set Level Approaches | | | | | | |
|---|---|---|---|---|---|---|---|---|
| **Trait Type** | **Metric** | **BANN** | **BANN-SS** | **RSS** | **PEGASUS** | **SKAT** | **MAGMA** | **GSEA** |
| **Sparse** | **Power** | **0.745 (0.093)** | 0.698 (0.102) | 0.702 (0.089) | 0.634 (0.104) | 0.521 (0.063) | 0.561 (0.097) | 0.481 (0.128) |
| | **FDR** | 0.146 (0.099) | **0.118 (0.097)** | 0.152 (0.107) | 0.311(0.132) | 0.275(0.126) | 0.262 (0.088) | 0.288 (0.101) |
| **Polygenic** | **Power** | 0.326 (0.123) | 0.273 (0.104) | 0.282 ( 0.094) | **0.382 (0.077)** | 0.216 (0.122) | 0.262(0.085) | 0.117 (0.035) |
| | **FDR** | **0.254 (0.052)** | 0.266 (0.092) | 0.261 (0.097) | 0.461 (0.201) | 0.392 (0.310) | 0.280 (0.199) | 0.457 (0.178) |

**Table S8. Comparing the empirical power and false discovery rates (FDR) of the BANNs framework against competing SNP and SNP-set mapping approaches in simulations with population structure (European cohort).** Here, quantitative traits are simulated to have broad-sense heritability of $H^2 = 0.6$ with contributions from both additive and epistatic effects set (i.e., $\rho = 0.5$). We consider two different trait architectures: sparse where only 1% of SNP-sets are enriched for the trait; and polygenic where 10% of SNP-sets are enriched. We set the number of causal SNPs with non-zero effects to be 1% and 10% of all SNPs located within the enriched SNP-sets, respectively. **(Top)** Competing SNP-level mapping approaches include: CAVIAR [63], SuSiE [64], and FINEMAP [65]. The software for SuSiE requires an input $\ell$ which fixes the maximum number of causal SNPs in the model. We display results when this input number is high ($\ell = 3000$) and when this input number is low ($\ell = 10$). **(Bottom)** Competing SNP-set mapping approaches include: RSS [7], PEGASUS [66], GBJ [67], SKAT [68], GSEA [69], and MAGMA [70]. Results for the BANN, BANN-SS, and other Bayesian methods are evaluated based on the "median probability criterion" (i.e., PIPs for SNPs and SNP-sets greater than 0.5) [51]. Results for the frequentist approaches are based on Bonferroni-corrected thresholds for multiple hypothesis testing ($P = 0.05/36518 = 1.37 \times 10^{-6}$ at the SNP-level and $P = 0.05/2816 = 1.78 \times 10^{-5}$ at the SNP-set level, respectively). All results are based on 100 replicates and standard deviations of the estimates across runs are given in the parentheses. Approaches with the greatest power are bolded in purple, while methods with the lowest FDR is bolded in blue.

| Simulation Parameters | | Average Run Time (seconds) | | | | |
|---|---|---|---|---|---|---|
| **SNPs** | **Samples Sizes** | **BANN** | **SuSiE (low)** | **SuSiE (high)** | **CAVIAR** | **FINEMAP** |
| 2500 | 1000 | 3.34 | 1.89 | 4.22 | 8.21 | 56.99 |
| | 2000 | 6.71 | 2.87 | 8.72 | 8.21 | 56.99 |
| | 4000 | 10.82 | 8.42 | 13.63 | 8.21 | 56.99 |
| 5000 | 1000 | 7.42 | 2.49 | 7.12 | 31.48 | 102.58 |
| | 2000 | 13.21 | 5.04 | 21.84 | 31.48 | 102.58 |
| | 4000 | 21.34 | 9.45 | 32.81 | 31.48 | 102.58 |
| 10000 | 1000 | 31.39 | 3.52 | 52.24 | 118.98 | 145.51 |
| | 2000 | 127.18 | 10.22 | 159.97 | 118.98 | 145.51 |
| | 4000 | 318.81 | 22.62 | 754.63 | 118.98 | 145.51 |

**Table S9. Computational time for running Bayesian annotated neural networks (BANNs) and other SNP-level association mapping approaches, as a function of the total number SNPs analyzed and the number of samples in the data.** Methods compared include: BANNs, CAVIAR [63], SuSiE [64], and FINEMAP [65]. Each table entry represents the average computation time (in seconds) it takes each approach to analyze a dataset of the size indicated. Run times were measured on an Intel i5-8259U CPU with base frequency of 2.30GHz, turbo frequency of 3.80GHz, and memory 16GB 2133 MHz LPDDR3. Here, we used 4 cores for parallelization when applicable. The software for SuSiE requires an input $\ell$ which fixes the maximum number of causal SNPs in the model. We display results when this input parameter is high ($\ell = 3000$) and when this input parameter is low ($\ell = 10$). Note that we implemented BANNs using the `Python 3` version of the software, and the timing for its variational algorithm includes inference on both SNPs and SNP-sets. CAVIAR and FINEMAP are set up to work with GWA summary statistics, so their inputs (and timing) are the same irrespective of the sample size.

| Simulation Parameters | | Average Run Time (seconds) | | | | | | |
|---|---|---|---|---|---|---|---|---|
| SNP-Sets | SNPs per SNP-set | BANN | RSS | PEGASUS | GBJ | SKAT | MAGMA | GSEA |
| 250 | 10 | 12.58 | 13.12 | 2.41 | 2.68 | 2.13 | 0.03 | 2.48 |
| | 20 | 44.32 | 58.21 | 2.13 | 5.18 | 3.82 | 0.08 | 4.68 |
| | 40 | 189.44 | 224.62 | 2.22 | 9.64 | 6.47 | 0.18 | 8.51 |
| 500 | 10 | 48.92 | 59.31 | 5.11 | 5.37 | 5.23 | 0.09 | 5.31 |
| | 20 | 223.14 | 244.07 | 5.02 | 11.26 | 9.22 | 0.21 | 11.12 |
| | 40 | 965.48 | 1026.12 | 5.72 | 27.91 | 14.84 | 0.24 | 20.36 |
| 1000 | 10 | 194.62 | 249.57 | 8.67 | 12.27 | 11.31 | 0.72 | 11.41 |
| | 20 | 1213.19 | 2176.33 | 8.93 | 27.62 | 18.16 | 1.48 | 24.93 |
| | 40 | 6823.31 | 14495.72 | 10.21 | 61.37 | 30.83 | 4.26 | 60.82 |

**Table S10. Computational time for running Bayesian annotated neural networks (BANNs) and other SNP-set level enrichment approaches, as a function of the total number SNP-sets analyzed and the number of SNPs within each SNP-set.** Methods compared include: BANNs, RSS [7], PEGASUS [66], GBJ [67], SKAT [68], GSEA [69], and MAGMA [70]. Here, we simulated 10 datasets for each pair of parameter values (number of SNP-sets analyzed and number of SNPs within each SNP-set). Sample size was held constant at $n = 10{,}000$ individuals. Each table entry represents the average computation time (in seconds) it takes each approach to analyze a dataset of the size indicated. Run times were measured on an Intel i5-8259U CPU with base frequency of 2.30GHz, turbo frequency of 3.80GHz, and memory 16GB 2133 MHz LPDDR3. Here, we used 4 cores for parallelization when applicable. Note that PEGASUS, GBJ, SKAT, and MAGMA are score-based methods and, thus, are expected to take the least amount of time to run. Both the BANNs framework and RSS are regression-based methods. The increased computational burden of these approaches results from its need to do (approximate) Bayesian posterior inference; however, the sparse and partially connected architecture of the BANNs model allows it to scale more favorably for larger dimensional datasets. Note that we implemented BANNs using the `Python 3` version of the software, and the timing for its variational algorithm includes inference on both SNPs and SNP-sets.

**Table S11. SNP and SNP-set results for body mass index (BMI) in the heterogenous stock of mice from the Wellcome Trust Centre for Human Genetics.** We analyze $J \approx$ 10,000 SNPs and $G = 1,925$ SNP-sets from $N =$ 1,814 mice—with specific numbers varying slightly depending on the quality control procedure for each phenotype (Supporting Information, Section 6). Here, SNP-set annotations are based on gene boundaries defined by the Mouse Genome Informatics database (see URLs listed in the main text). Unannotated SNPs located within the same genomic region were labeled as being within the "intergenic region" between two genes. This file gives the posterior inclusion probabilities (PIPs) for the input and hidden layer neural network weights after fitting the BANNs model on the individual-level data. We assess significance for both SNPs and SNP-sets according to the "median probability model" threshold [51] (i.e., PIP $\geq$ 0.5). Page #1 provides the variant-level association mapping results with columns corresponding to: (1) chromosome; (2) SNP ID; (3) chromosomal position in base-pair (bp) coordinates; (4) SNP PIP; and (5) SuSiE PIP, which corresponds to SNP-level posterior inclusion probabilities computed by SuSiE [64]. Page #2 provides the SNP-set level enrichment results with columns corresponding to: (1) chromosome; (2) SNP-set ID; (3-4) the starting and ending position of the SNP-set chromosomal boundaries; (5) SNP-set PIP; (6) RSS PIP, which corresponds to the posterior inclusion probabilities computed by RSS [7]; (7) the number of SNPs that have been annotated within each SNP-set; (8) the "top" associated SNP within each SNP-set; (9) the PIP of each top SNP.

**Table S12. SNP and SNP-set results for body weight in the heterogenous stock of mice from the Wellcome Trust Centre for Human Genetics.** We analyze $J \approx$ 10,000 SNPs and $G = 1,925$ SNP-sets from $N =$ 1,814 mice—with specific numbers varying slightly depending on the quality control procedure for each phenotype (Supporting Information, Section 6). Here, SNP-set annotations are based on gene boundaries defined by the Mouse Genome Informatics database (see URLs in the main text). Unannotated SNPs located within the same genomic region were labeled as being within the "intergenic region" between two genes. This file gives the posterior inclusion probabilities (PIPs) for the input and hidden layer neural network weights after fitting the BANNs model on the individual-level data. We assess significance for both SNPs and SNP-sets according to the "median probability model" threshold [51] (i.e., PIP $\geq$ 0.5). Page #1 provides the variant-level association mapping results with columns corresponding to: (1) chromosome; (2) SNP ID; (3) chromosomal position in base-pair (bp) coordinates; (4) SNP PIP; and (5) SuSiE PIP, which corresponds to SNP-level posterior inclusion probabilities computed by SuSiE [64]. Page #2 provides the SNP-set level enrichment results with columns corresponding to: (1) chromosome; (2) SNP-set ID; (3-4) the starting and ending position of the SNP-set chromosomal boundaries; (5) SNP-set PIP; (6) RSS PIP, which corresponds to the posterior inclusion probabilities computed by RSS [7]; (7) the number of SNPs that have been annotated within each SNP-set; (8) the "top" associated SNP within each SNP-set; (9) the PIP of each top SNP.

**Table S13. SNP and SNP-set results for percentage of CD8+ cells in the heterogenous stock of mice from the Wellcome Trust Centre for Human Genetics.** We analyze $J \approx$ 10,000 SNPs and $G = 1,925$ SNP-sets from $N =$ 1,814 mice—with specific numbers varying slightly depending on the quality control procedure for each phenotype (Supporting Information, Section 6). Here, SNP-set annotations are based on gene boundaries defined by the Mouse Genome Informatics database (see URLs listed in the main text). Unannotated SNPs located within the same genomic region were labeled as being within the "intergenic region" between two genes. This file gives the posterior inclusion probabilities (PIPs) for the input and hidden layer neural network weights after fitting the BANNs model on the individual-level data. We assess significance for both SNPs and SNP-sets according to the "median probability model" threshold [51] (i.e., PIP $\geq$ 0.5). Page #1 provides the variant-level association mapping results with columns corresponding to: (1) chromosome; (2) SNP ID; (3) chromosomal position in base-pair (bp) coordinates; (4) SNP PIP; and (5) SuSiE PIP, which corresponds to SNP-level posterior inclusion probabilities computed by SuSiE [64]. Page #2 provides the SNP-set level enrichment results with columns corresponding to: (1) chromosome; (2) SNP-set ID; (3-4) the starting and ending position of the SNP-set chromosomal boundaries; (5) SNP-set PIP; (6) RSS PIP, which corresponds to the posterior inclusion probabilities computed by RSS [7]; (7) the number of SNPs that have been annotated within each SNP-set; (8) the "top" associated SNP within each SNP-set; (9) the PIP of each top SNP.

**Table S14. SNP and SNP-set results for high-density lipoprotein (HDL) cholesterol in the heterogenous stock of mice from the Wellcome Trust Centre for Human Genetics.** We analyze $J \approx$ 10,000 SNPs and $G = 1,925$ SNP-sets from $N =$ 1,814 mice—with specific numbers varying slightly depending on the quality control procedure for each phenotype (Supporting Information, Section 6). Here, SNP-set annotations are based on gene boundaries defined by the Mouse Genome Informatics database (see URLs listed in the main text). Unannotated SNPs located within the same genomic region were labeled as being within the "intergenic region" between two genes. This file gives the posterior inclusion probabilities (PIPs) for the input and hidden layer neural network weights after fitting the BANNs model on the individual-level data. We assess significance for both SNPs and SNP-sets according to the "median probability model" threshold [51] (i.e., PIP $\geq$ 0.5). Page #1 provides the variant-level association mapping results with columns corresponding to: (1) chromosome; (2) SNP ID; (3) chromosomal position in base-pair (bp) coordinates; (4) SNP PIP; and (5) SuSiE PIP, which corresponds to SNP-level posterior inclusion probabilities computed by SuSiE [64]. Page #2 provides the SNP-set level enrichment results with columns corresponding to: (1) chromosome; (2) SNP-set ID; (3-4) the starting and ending position of the SNP-set chromosomal boundaries; (5) SNP-set PIP; (6) RSS PIP, which corresponds to the posterior inclusion probabilities computed by RSS [7]; (7) the number of SNPs that have been annotated within each SNP-set; (8) the "top" associated SNP within each SNP-set; (9) the PIP of each top SNP.

**Table S15. SNP and SNP-set results for low-density lipoprotein (LDL) cholesterol in the heterogenous stock of mice from the Wellcome Trust Centre for Human Genetics.** We analyze $J \approx 10{,}000$ SNPs and $G = 1,925$ SNP-sets from $N = 1{,}814$ mice—with specific numbers varying slightly depending on the quality control procedure for each phenotype (Supporting Information, Section 6). Here, SNP-set annotations are based on gene boundaries defined by the Mouse Genome Informatics database (see URLs listed in the main text). Unannotated SNPs located within the same genomic region were labeled as being within the "intergenic region" between two genes. This file gives the posterior inclusion probabilities (PIPs) for the input and hidden layer neural network weights after fitting the BANNs model on the individual-level data. We assess significance for both SNPs and SNP-sets according to the "median probability model" threshold [51] (i.e., PIP $\geq 0.5$). Page #1 provides the variant-level association mapping results with columns corresponding to: (1) chromosome; (2) SNP ID; (3) chromosomal position in base-pair (bp) coordinates; (4) SNP PIP; and (5) SuSiE PIP, which corresponds to SNP-level posterior inclusion probabilities computed by SuSiE [64]. Page #2 provides the SNP-set level enrichment results with columns corresponding to: (1) chromosome; (2) SNP-set ID; (3-4) the starting and ending position of the SNP-set chromosomal boundaries; (5) SNP-set PIP; (6) RSS PIP, which corresponds to the posterior inclusion probabilities computed by RSS [7]; (7) the number of SNPs that have been annotated within each SNP-set; (8) the "top" associated SNP within each SNP-set; (9) the PIP of each top SNP.

**Table S16. SNP and SNP-set results for mean corpuscular hemoglobin (MCH) in the heterogenous stock of mice from the Wellcome Trust Centre for Human Genetics.** We analyze $J \approx 10{,}000$ SNPs and $G = 1,925$ SNP-sets from $N = 1{,}814$ mice—with specific numbers varying slightly depending on the quality control procedure for each phenotype (Supporting Information, Section 6). Here, SNP-set annotations are based on gene boundaries defined by the Mouse Genome Informatics database (see URLs listed in the main text). Unannotated SNPs located within the same genomic region were labeled as being within the "intergenic region" between two genes. This file gives the posterior inclusion probabilities (PIPs) for the input and hidden layer neural network weights after fitting the BANNs model on the individual-level data. We assess significance for both SNPs and SNP-sets according to the "median probability model" threshold [51] (i.e., PIP $\geq 0.5$). Page #1 provides the variant-level association mapping results with columns corresponding to: (1) chromosome; (2) SNP ID; (3) chromosomal position in base-pair (bp) coordinates; (4) SNP PIP; and (5) SuSiE PIP, which corresponds to SNP-level posterior inclusion probabilities computed by SuSiE [64]. Page #2 provides the SNP-set level enrichment results with columns corresponding to: (1) chromosome; (2) SNP-set ID; (3-4) the starting and ending position of the SNP-set chromosomal boundaries; (5) SNP-set PIP; (6) RSS PIP, which corresponds to the posterior inclusion probabilities computed by RSS [7]; (7) the number of SNPs that have been annotated within each SNP-set; (8) the "top" associated SNP within each SNP-set; (9) the PIP of each top SNP.

| Trait | SNP-Set | Chr | PIP($g$) | Rank | RSS PIP | RSS Rank | Top SNP | PIP($j$) | SuSiE PIP | SuSiE Rank | Ref(s) |
|---|---|---|---|---|---|---|---|---|---|---|---|
| HDL | *C10rf213-TCEA3* | 1 | 0.998 | 2 | 0.121 | 162 | rs176141 | 0.298 | 0.120 | 19 | [78, 79] |
| | *ST18-FAM150A* | 8 | 0.9169 | 4 | 0.959 | 4 | rs11990693 | 0.999 | 0.277 | 8 | [80] |
| | *LPL-SLC18A1* | 8 | 0.734 | 5 | 0.999 | 2 | rs17482753♣ | 0.999 | 0.191 | 11 | [81, 82] |
| LDL | *AQP9-LIPC* | 15 | 0.913 | 1 | 0.198 | 29 | rs4775041 | 0.422 | 0.238 | 4 | [83, 84] |
| | *TRMU-CELSR1* | 22 | 0.678 | 2 | 0.895 | 3 | rs5768939 | 0.379 | 0.185 | 5 | [85–88] |
| | *C10orf92-C10orf93* | 10 | 0.537 | 3 | 0.125 | 157 | rs10781583 | 0.215 | 0.0377 | 26 | — |

**Table S17. Notable enriched SNP-sets after applying the BANNs framework to high-density and low-density lipoprotein (HDL and LDL, respectively) traits in the Framingham Heart Study [47] where each SNP-set annotation has been augmented with a ±500 kilobase (kb) buffer to account for possible regulatory elements.** Here, SNP-set annotations are based on gene boundaries defined by the NCBI's RefSeq database in the UCSC Genome Browser [49]. Unannotated SNPs located within the same genomic region were labeled as being within the "intergenic region" between two genes. These regions are labeled as *Gene1-Gene2* in the table. Posterior inclusion probabilities (PIP) for the input and hidden layer weights are derived by fitting the BANNs model on individual-level data. A SNP-set is considered enriched if it has a PIP($g$) $\geq 0.5$ (i.e., the "median probability model" threshold [51]). We report the "top" associated SNP within each region and its corresponding PIP($j$). We also report the corresponding SNP and SNP-set level results after running SuSiE [64] and RSS [7] on these same traits, respectively. The last column details references and literature sources that have previously suggested some level of association or enrichment between the each genomic region and the traits of interest. See Tables S18 and S19 for the complete list of SNP and SNP-set level results. ♣: SNPs and SNP-sets replicated in an independent analysis of ten thousand randomly sampled individuals of European ancestry from the UK Biobank [50].

**Table S18. SNP and SNP-set results for high-density lipoprotein (HDL) cholesterol in individuals assayed within the Framingham Heart Study.** We analyze $J$ = 394,174 SNPs and $G = 18,364$ SNP-sets from $N$ = 6,950 people. Here, SNP-set annotations are based on gene boundaries defined by the NCBI's RefSeq database in the UCSC Genome Browser [49]. Unannotated SNPs located within the same genomic region were labeled as being within the "intergenic region" between two genes. This file gives the posterior inclusion probabilities (PIPs) for the input and hidden layer neural network weights after fitting the BANNs model on the individual-level data. We assess significance for both SNPs and SNP-sets according to the "median probability model" threshold [51] (i.e., PIP $\geq$ 0.5). Page #1 provides the variant-level association mapping results with columns corresponding to: (1) chromosome; (2) SNP ID; (3) chromosomal position in base-pair (bp) coordinates; (4) SNP PIP; and (5) SuSiE PIP, which corresponds to SNP-level posterior inclusion probabilities computed by SuSiE [64]. Page #2 provides the SNP-set level enrichment results with columns corresponding to: (1) chromosome; (2) SNP-set ID; (3-4) the starting and ending position of the SNP-set chromosomal boundaries; (5) SNP-set PIP; (6) RSS PIP, which corresponds to the posterior inclusion probabilities computed by RSS [7]; (7) the number of SNPs that have been annotated within each SNP-set; (8) the "top" associated SNP within each SNP-set; (9) the PIP of each top SNP. Pages #3 and #4 provide similar results based on analyses where each SNP-set annotation has been augmented with a ±500 kilobase (kb) buffer to account for possible regulatory elements.

**Table S19. SNP and SNP-set results for low-density lipoprotein (LDL) cholesterol in individuals assayed within the Framingham Heart Study.** We analyze $J$ = 394,174 SNPs and $G = 18,364$ SNP-sets from $N$ = 6,950 people. Here, SNP-set annotations are based on gene boundaries defined by the NCBI's RefSeq database in the UCSC Genome Browser [49]. Unannotated SNPs located within the same genomic region were labeled as being within the "intergenic region" between two genes. This file gives the posterior inclusion probabilities (PIPs) for the input and hidden layer neural network weights after fitting the BANNs model on the individual-level data. We assess significance for both SNPs and SNP-sets according to the "median probability model" threshold [51] (i.e., PIP $\geq$ 0.5). Page #1 provides the variant-level association mapping results with columns corresponding to: (1) chromosome; (2) SNP ID; (3) chromosomal position in base-pair (bp) coordinates; (4) SNP PIP; and (5) SuSiE PIP, which corresponds to SNP-level posterior inclusion probabilities computed by SuSiE [64]. Page #2 provides the SNP-set level enrichment results with columns corresponding to: (1) chromosome; (2) SNP-set ID; (3-4) the starting and ending position of the SNP-set chromosomal boundaries; (5) SNP-set PIP; (6) RSS PIP, which corresponds to the posterior inclusion probabilities computed by RSS [7]; (7) the number of SNPs that have been annotated within each SNP-set; (8) the "top" associated SNP within each SNP-set; (9) the PIP of each top SNP. Pages #3 and #4 provide similar results based on analyses where each SNP-set annotation has been augmented with a ±500 kilobase (kb) buffer to account for possible regulatory elements.

**Table S20. Complete summary of the results after applying BANNs, SuSiE [64], and RSS [7] to high-density lipoprotein (HDL) and low-density lipoprotein (LDL) cholesterol in both individuals assayed within the Framingham Heart Study and ten thousand randomly sampled individuals of European ancestry from the UK Biobank.** The first page compares the overlap of significant SNPs and SNP-sets found by each method according to the "median probability model" threshold [51] (i.e., PIP $\geq 0.5$) in the Framingham Heart Study. The second page lists how many SNPs and SNP-sets were replicated for each method when analyzing the independent dataset from the UK Biobank. Results are based on defining gene boundaries in two ways: ($a$) we use the UCSC gene boundary definitions directly, and ($b$) we augment the gene boundaries by adding SNPs within a $\pm 500$ kilobase (kb) buffer to account for possible regulatory elements.

**Table S21. SNP and SNP-set results for high-density lipoprotein (HDL) cholesterol in ten thousand randomly sampled individuals of European ancestry from the UK Biobank.** We analyze the same $J$ = 394,174 SNPs and $G = 18,364$ SNP-sets used in the Framingham Heart Study analyses. Here, SNP-set annotations are based on gene boundaries defined by the NCBI's RefSeq database in the UCSC Genome Browser [49]. Unannotated SNPs located within the same genomic region were labeled as being within the "intergenic region" between two genes. This file gives the posterior inclusion probabilities (PIPs) for the input and hidden layer neural network weights after fitting the BANNs model on the individual-level data. We assess significance for both SNPs and SNP-sets according to the "median probability model" threshold [51] (i.e., PIP $\geq 0.5$). Page #1 provides the variant-level association mapping results with columns corresponding to: (1) chromosome; (2) SNP ID; (3) chromosomal position in base-pair (bp) coordinates; (4) SNP PIP; and (5) SuSiE PIP, which corresponds to SNP-level posterior inclusion probabilities computed by SuSiE [64]. Page #2 provides the SNP-set level enrichment results with columns corresponding to: (1) chromosome; (2) SNP-set ID; (3-4) the starting and ending position of the SNP-set chromosomal boundaries; (5) SNP-set PIP; (6) RSS PIP, which corresponds to the posterior inclusion probabilities computed by RSS [7]; (7) the number of SNPs that have been annotated within each SNP-set; (8) the "top" associated SNP within each SNP-set; (9) the PIP of each top SNP. Pages #3 and #4 provide similar results based on analyses where each SNP-set annotation has been augmented with a $\pm 500$ kilobase (kb) buffer to account for possible regulatory elements.

**Table S22. SNP and SNP-set results for low-density lipoprotein (LDL) cholesterol in ten thousand randomly sampled individuals of European ancestry from the UK Biobank.** We analyze the same $J = 394{,}174$ SNPs and $G = 18,364$ SNP-sets used in the Framingham Heart Study analyses. Here, SNP-set annotations are based on gene boundaries defined by the NCBI's RefSeq database in the UCSC Genome Browser [49]. Unannotated SNPs located within the same genomic region were labeled as being within the "intergenic region" between two genes. This file gives the posterior inclusion probabilities (PIPs) for the input and hidden layer neural network weights after fitting the BANNs model on the individual-level data. We assess significance for both SNPs and SNP-sets according to the "median probability model" threshold [51] (i.e., PIP $\geq 0.5$). Page #1 provides the variant-level association mapping results with columns corresponding to: (1) chromosome; (2) SNP ID; (3) chromosomal position in base-pair (bp) coordinates; (4) SNP PIP; and (5) SuSiE PIP, which corresponds to SNP-level posterior inclusion probabilities computed by SuSiE [64]. Page #2 provides the SNP-set level enrichment results with columns corresponding to: (1) chromosome; (2) SNP-set ID; (3-4) the starting and ending position of the SNP-set chromosomal boundaries; (5) SNP-set PIP; (6) RSS PIP, which corresponds to the posterior inclusion probabilities computed by RSS [7]; (7) the number of SNPs that have been annotated within each SNP-set; (8) the "top" associated SNP within each SNP-set; (9) the PIP of each top SNP. Pages #3 and #4 provide similar results based on analyses where each SNP-set annotation has been augmented with a $\pm 500$ kilobase (kb) buffer to account for possible regulatory elements.
